# Cancer Prognosis According to Parthanatos Features

**DOI:** 10.1101/2021.05.24.445484

**Authors:** Alessandra Messikommer, Bruktawit Maru, Katja Seipel, Peter J. M. Valk, Alexandre P.A. Theocharides, Thomas Pabst, Maureen McKeague, Nathan W. Luedtke

## Abstract

For nearly 50 years, translational research studies aimed at improving chemotherapy-induced killing of cancer cells have focused on the induction of apoptosis. Here we show that a PARP-1-mediated programmed cell death mechanism “parthanatos” is associated with the successful, front-line treatment of a common cancer. Peripheral blood mononuclear cells (PBMCs) from healthy human donors (10 of 10 tested), as well as primary cancer cells from approximately 50% of acute myeloid leukemia (AML) patients (n = 18 of 39 tested, French-American-British (FAB) subtypes M4 and M5) exhibited two distinctive features of parthanatos upon treatment with a front-line drug combination of cytarabine and an anthracycline. Statistically significant improvements in survival rates were observed in the parthanatos positive versus parthanatos negative AML patient groups (HR = 0.22 – 0.38, *p* = 0.002 – 0.05). Near-median expression of *PARP1* mRNA was associated with a 50% longer survival time (HR = 0.66, *p* = 0.01), and the poly [ADP-ribose] polymerase (PARP) inhibitor Olaparib exhibited antagonistic activities against ara-C and idarubicin in primary blood monocytes from healthy donors as well as primary cancer isolates from ~50% of AML patients. Together these results suggest that PARP activity is a prognostic biomarker for AML subtypes M4 and M5 and support the relevance of parthanatos in curative chemotherapy of AML.

**In Brief:** Messikommer and co-workers report that PARP-1-mediated programmed cell death is associated with successful, front-line treatment of acute myeloid leukemia (AML).

**Highlights:**

- The first-line cancer drug cytarabine (ara-C) induces parthanatos or apoptosis, depending on the specific AML cell line being treated.
- OCI-AML3 cells undergo parthanatos or apoptosis, depending on the specific drug being added.
- The presence of two parthanatos features in primary cancer cells from AML patients (n = 18 of 39 tested) having French-American-British (FAB) subclassifications M4 or M5 is associated with four-fold improved survival (HR = 0.23, *p* = 0.01) following curative chemotherapy with ara-C and an anthracycline.
- The poly [ADP-ribose] polymerase (PARP) inhibitor Olaparib exhibits antagonistic activities against ara-C and idarubicin in primary blood monocytes from healthy donors as well as primary cancer isolates from ~50% of AML patients.
- Near-median expression of *PARP1* mRNA is associated with a 50% increase in survival time (HR = 0.66, p = 0.01) of AML patients following chemotherapy with ara-C and idarubicin.

**GRAPHICAL ABSTRACT:** 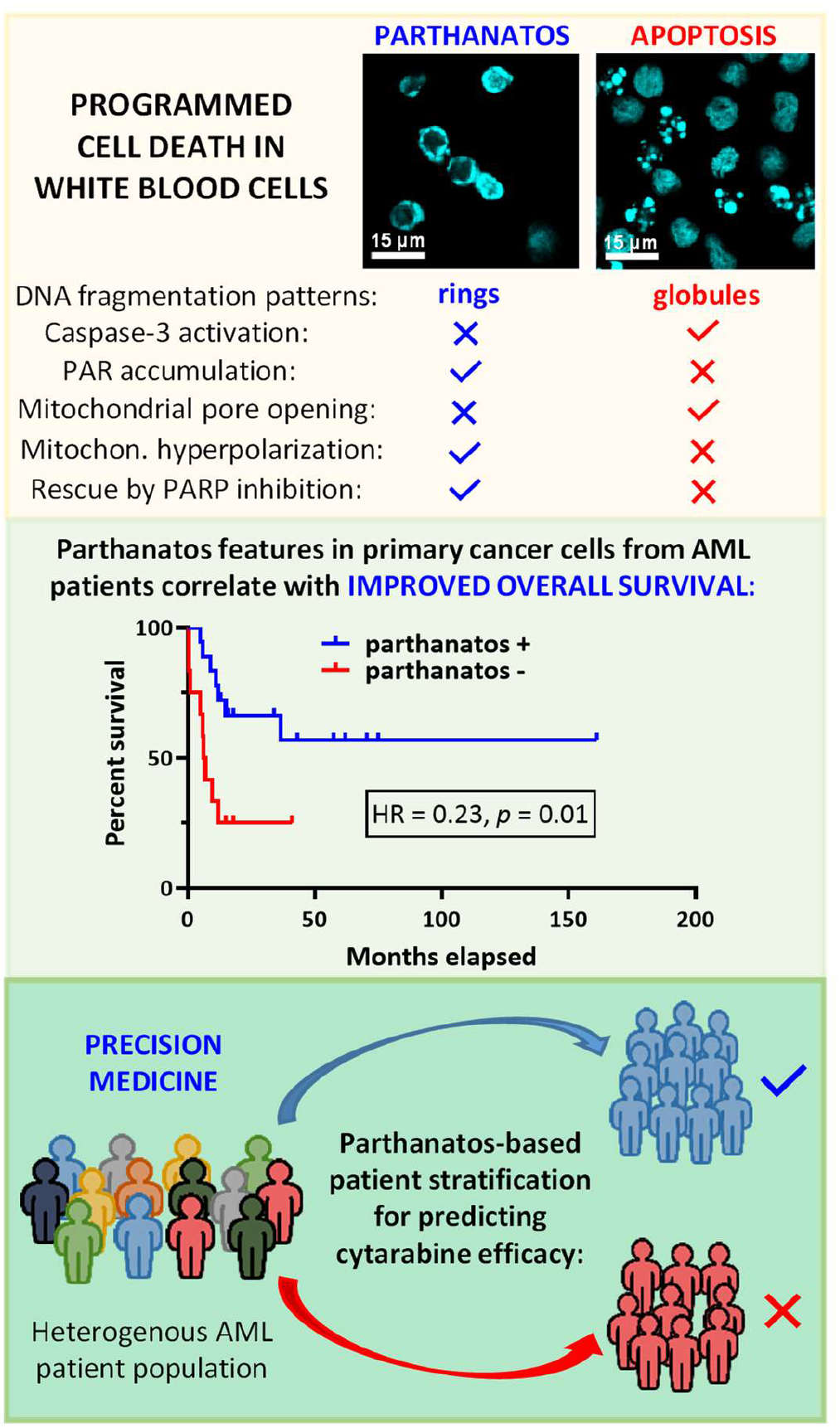

## INTRODUCTION

Acute myeloid leukemia (AML) is the most common and malicious form of leukemia in adults, with an approximate 30% five-year overall survival rate for patients receiving curative therapies (Costa et al., 2017). The histology-based French-American-British (FAB) subclassification system helped identify acute promyelocytic leukemia (APL, subtype M3) as a separate and highly curable subtype of AML, but it only accounts for ~10% of AML patients (Cicconi and Lo-Coco, 2016). The remaining 90% of AML patients are classified by a prognosis-based system established by the World Health Organization and European LeukemiaNet (ELN) that divides patients into three risk groups (favorable, intermediate, and adverse) based on the presence of cytogenetic and mutational biomarkers. Despite a growing number of new auxiliary and palliative AML therapies becoming available (Kopmar and Estey, 2019), the frontline, curative treatment of eligible patients from all three risk groups commences with a 7-day infusion of cytarabine (ara-C) that is typically augmented with an anthracycline for three days known as “7+3” induction chemotherapy (Estey, 2018). Even after 50 years of widespread clinical use, the mechanisms responsible for ara-C’s selective killing of white blood cells remain poorly understood. Previous studies have focused on metabolic incorporation of ara-C into DNA and downstream apoptosis (Cassier et al., 2017; Deng et al., 2017; Harrison et al., 2016; Schneider et al., 2017; Vincelette and Yun, 2014). However, recent reports suggest that ara-C incorporation into primer RNA is responsible for its therapeutic efficacy (Messikommer et al., 2020; Triemer et al., 2018), and active killing of AML primary isolates by ara-C can be a caspase-independent process (Carter et al., 2003).

DNA damage and stimulation of apoptosis have been focal areas of chemotherapeutic drug development for leukemia and other cancers for nearly 50 years (Carneiro and El-Deiry, 2020). Cells that lose tumor suppressor activities by p53 mutation (Kandoth et al., 2013) can gain resistance against standard chemotherapeutic drugs by dysregulation of apoptosis (Mohammad et al., 2015; Wong et al., 2015). Targeted therapies that re-sensitize cells towards apoptosis such as B-cell leukemia/lymphoma-2 (Bcl-2) family inhibitors (Jonas and Pollyea, 2019) as well as p53 “reactivators”(Pan et al., 2017) can overcome apoptosis resistance. However, primary AML cells treated with ara-C can undergo caspase-independent programmed cell death (Carter et al., 2003), and caspase-3 activation failed to predict ara-C and anthracycline drug sensitivity of primary AML isolates from 42 patients (Staib et al., 2005). A potential explanation for these observations is that healthy and diseased white blood cells may undergo non-apoptotic forms of programmed cell death during chemotherapy such as autophagic cell death (Auberger and Puissant, 2017; Kajiume and Kobayashi, 2018), necroptosis (Huang et al., 2018; Mezzatesta and Bornhauser, 2019), pyroptosis (Xia et al., 2019), and/or parthanatos (Cloux et al., 2019).

Parthanatos has been recognized as a distinct mode of programmed cell death by the Nomenclature Committee on Cell Death since 2012 (Galluzzi et al., 2018; Galluzzi et al., 2012). Poly [ADP-ribose] polymerase 1 (PARP-1) is the primary orchestrator of parthanatos (Wang et al., 2011). Abnormal expression of *PARP1* is associated with poor clinical outcomes in breast cancer (Rojo et al., 2012) and AML (Li et al., 2018; Pashaiefar et al., 2018; Wang et al., 2015). PARP-1-mediated cell death has been reported in white blood cells treated with reactive oxygen species (Cloux et al., 2019; Robinson et al., 2019). Parthanatos proceeds via activation of PARP-1 by chromatin stress (Prokhorova et al., 2020), accumulation of poly-ADP ribose (PAR), and translocation of apoptosis-inducing factor (AIF) from the mitochondria into the nucleus where it activates nuclease macrophage migration inhibitory factor (MIF) (Andrabi et al., 2008; David et al., 2009; Wang et al., 2016). Once activated, MIF selectively cleaves single-stranded DNA to generate very large chromatin fragments of ~50 kb (Fatokun et al., 2014; Wang et al., 2016). In contrast to both necroptosis (Mezzatesta and Bornhauser, 2019) and pyroptosis (Xia et al., 2019), parthanatos proceeds with retention of plasma membrane integrity and flipping of phosphatidylserine onto the outer surface of the cell (Leong et al., 2010; Zuber et al., 2009). As such, parthanatos cannot be distinguished from apoptosis using standard assay kits, for example by using only annexin V and propidium iodide staining (Fatokun et al., 2014; Wang et al., 2009). Additional experiments are required to distinguish between parthanatos and apoptosis, such as chromatin fragment analysis (Huang et al., 1995; Wang et al., 2009) or evaluating the impact of PARP-1 inhibition on drug sensitivity (Cloux et al., 2019).

Here, we report that PARP-1-mediated programmed cell death is associated with successful, frontline treatment of a common cancer. The presence/absence of two key parthanatos features: PARP-dependent changes in drug sensitivity and chromatin fragmentation morphologies are highly correlated with each other in samples taken from 39 AML patients (χ^2^_(3)_ = 21.8, p = 0.0001). Among the 35 patients receiving intensive chemotherapy including ara-C, the group exhibiting one or both parthanatos features had a four-fold improved survival rate (HR = 0.24, p = 0.008) as compared to the parthanatos negative group. Furthermore, near-median expression of *PARP1* was associated with a two-fold lower risk of death (HR = 0.55, p = 0.003) in AML patients of FAB subgroups M4 and M5 with wild-type FLT3 receiving curative chemotherapy with ara-C and idarubicin. Together, these results demonstrate that parthanatos is a clinically relevant mechanism of programmed cell death in a first-line cancer therapy. The development of drugs that directly activate parthanatos by inhibiting putative parthanatos suppressors such as PARG or ARH3 (Andrabi et al., 2008; Houl et al., 2019; Leung, 2014; Mashimo et al., 2013) therefore provides an attractive new strategy to treat refractory/relapsed AML and possibly other cancers.

## RESULTS

### Cytarabine induces parthanatos in OCI-AML3 and apoptosis in OCI-AML2 cells

OCI-AML2 and OCI-AML3 cell lines were selected for initial studies since they are both derived from a common FAB subtype (M4, ~25% of AML) and they express wild type *FLT3* (Seipel et al., 2018), *PARP1* (Sanger, 2021), and *TP53* (Tiacci et al., 2012). Ara-C treatment of both OCI-AML2 and OCI-AML3 cells caused phosphatidylserine exposure on cell surfaces without loss of membrane integrity (Fig 1A). At ara-C concentrations near the EC50 value for each cell line, caspase-3 activation was present in OCI-AML2, but absent in OCI-AML3 cells (Fig 1B). As a positive control, the topoisomerase inhibitor camptothecin (CPT) (Rodriguez-Hernandez et al., 2006) was found to activate caspase-3 in both cell lines (Fig 1B). These results suggest that caspase-independent programmed cell death is operational in OCI-AML3 cells treated with ara-C. To further investigate differences in programmed cell death mechanisms, we measured changes in mitochondrial properties of treated cells. Consistent with the features of classical apoptosis, the opening of mitochondrial pores (arrow, Fig. 1C) as well as loss of mitochondrial membrane potentials (Fig. 1D) were observed in OCI-AML2 cells treated with ara-C. In contrast, OCI-AML3 cells did not form mitochondrial transition pores and ara-C treatment caused hyperpolarization of the mitochondrial membranes. These changes, together with the concurrent maintenance of cell membrane integrity, demonstrate that ara-C induces neither classical apoptosis nor necrosis in OCI-AML3 cells. To evaluate the possibility that OCI-AML3 cells undergo parthanatos upon ara-C treatment, we quantified changes in PAR accumulation (Andrabi et al., 2006) and AIF translocation (Yu et al., 2006). Indeed, OCI-AML3 cells accumulated PAR polymers (Fig 1E) and released AIF following ara-C treatment (Fig. 1F-G). Together these results demonstrate that OCI-AML2 cells undergo classical apoptosis upon addition of ara-C, whereas OCI-AML3 cells undergo parthanatos.

**Fig. 1.**
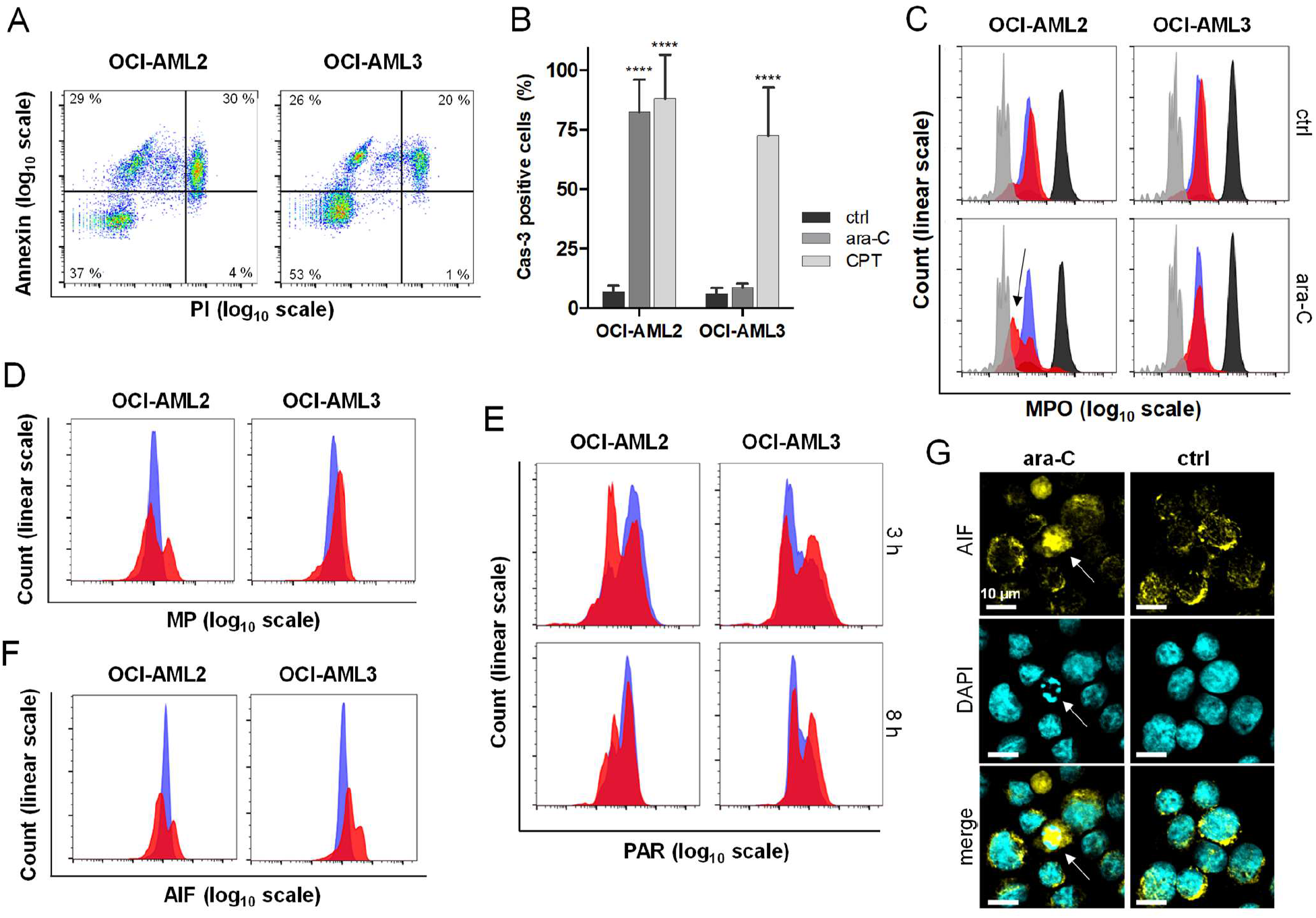
Cytarabine induces apoptosis in OCI-AML2 cells and parthanatos in OCI-AML3 cells. **A)** Phosphatidylserine exposure on the cell surface was detected by Annexin V staining, and membrane integrity was probed using propidium iodide (PI) following a 24 h treatment with cytarabine (ara-C). **B)** Ara-C treatment caused caspase-3 activity in OCI-AML2 (5 μM) but not OCI-AML3 cells (15 μM) after 8 hours. Ara-C concentrations were selected based on EC_50_ values for toxicity in each cell line. Camptothecin (CPT) was used as positive control for stimulating caspase-3 activity (Rodriguez-Hernandez et al., 2006). **C)** Upon treatment of OCI-AML2 cells (1 μM) or OCI-AML3 cells (10 μM) with ara-C for 24 h, fluorescence quenching indicating mitochondrial pore opening (MPO) in OCI-AML2 (black arrow) and not in OCI-AML3 cells according to the MitoProbe Transition Pore Assay and flow cytometry. Blue: untreated, red: ara-C treated, black: staining control calcein AM, grey: positive control ionomycin. **D)** Mitochondrial membrane potentials (MP) following ara-C treatment for 24 h. Blue: untreated, red: ara-C treated. **E)** After 3 h of ara-C treatment, PAR accumulated in OCI-AML3 cells but was diminished in OCI-AML2 cells according to immunofluorescent staining. Blue: untreated, red: ara-C treated. **F)** After 4 hours of ara-C treatment, higher levels of AIF staining were observed in treated OCI-AML3 cells as compared to OCI-AML2 cells. Blue: untreated, red: ara-C treated. **G)** Immunostaining of AIF and microscopy revealed the release of AIF in OCI-AML3 cells. The arrow highlights a cell with a characteristic “ring” structure of DNA and high abundance of released AIF. **** *p* < 0.0001. Control (“ctrl”) samples were not treated with ara-C.

### Different modes of programmed cell death are reflected in distinctive nuclear morphologies

DAPI staining and microscopy revealed the formation of globular chromatin fragments throughout the nuclei of OCI-AML2 cells treated with ara-C (Fig 2A). In contrast, chromatin fragments in a distinctive “ring-shape” pattern were observed at the nuclear periphery of treated OCI-AML3 cells (Fig 2A). We speculated that this unique chromatin morphology is associated with the cleavage of chromatin into a relatively small number of large fragments in late parthanatos (Huang et al., 1995; Wang et al., 2009). To evaluate this possibility, the TUNEL assay was used to compare the relative number of DNA cleavage sites in OCI-AML2 and OCI-AML3 cells treated with ara-C (Fig 2B). Indeed, a much larger number of DNA strand breaks was observed in treated OCI-AML2 versus OCI-AML3 cells, consistent with a higher frequency of DNA cleavage events in late apoptosis versus parthanatos. When treated with a clinically relevant, 17:1 ratio mixture of ara-C and the anthracycline drug idarubicin (Dohner et al., 2010), the OCI-AML3 cells again exhibited ring-shaped chromatin patterns (Fig 2C). In contrast, nuclear morphologies consistent with apoptosis were observed (Fig 2C) when the same OCI-AML3 cells were treated with the BH3 mimetic AT-101 (Gossypol) which inhibits anti-apoptotic Bcl family proteins to activate apoptosis (Pan et al., 2014; Paulus et al., 2014). These results suggest that OCI-AML3 cells can undergo parthanatos or apoptosis depending on the specific drug being added and that characteristic changes in nuclear morphologies reflect the type of programmed cell death mechanism in operation.

**Fig. 2.**
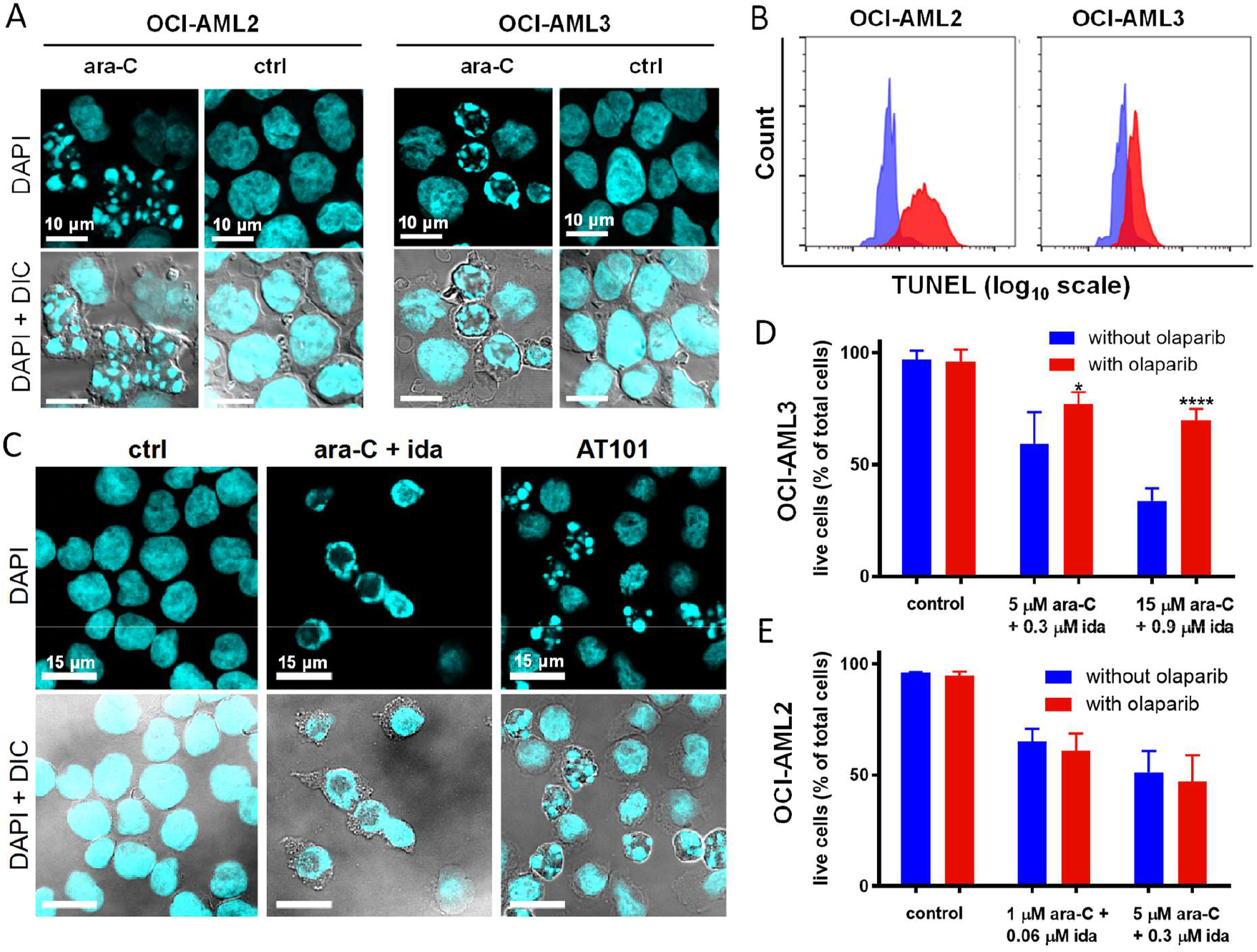
Additional parthanatos features in OCI-AML3 cells. **A**) DAPI staining revealed globular chromatin fragments of highly variable sizes throughout the nuclei of OCI-AML2 cells treated with 1 μM ara-C for 24 hours, whereas a relatively small number of large chromatin fragments are distributed in a ring-shaped pattern at the nuclear periphery of OCI-AML3 cells treated with 10 μM of ara-C for 24 hours. Ara-C concentrations were selected based on EC_50_ values for toxicity in each cell line. **B)** The TUNEL assay confirmed a relatively broad distribution of small DNA fragments in OCI-AML2 as compared to OCI-AML3 cells following treatment with ara-C. Blue: untreated, red: ara-C treated. **C)** Nuclear morphologies in OCI-AML3 cells following treatment with a clinically relevant 17:1 ratio mixture of ara-C and the anthracycline drug idarubicin (ida), as compared to the BH3 mimetic AT-101 (Gossypol) that selectively stimulates apoptosis. **D)** Live/dead staining and flow cytometry analysis of OCI-AML3 cells treated with 1 μM of the PARP inhibitor Olaparib for 24 h prior to treatment with ara-C and idarubicin in a fixed 17:1 ratio for 24 hours. **E)** Live/dead staining and flow cytometry analysis of OCI-AML2 cells treated with 1 μM of the PARP inhibitor Olaparib for 24 h prior to treatment with ara-C and idarubicin in a fixed 17:1 ratio for 24 hours. Control (“ctrl”) samples were not treated with ara-C or idarubicin; * p < 0.1; **** p < 0.0001; “DIC” = differential interference contrast image.

### PARP inhibition protects OCI-AML3 cells from ara-C and idarubicin-induced cell death

To further evaluate the presence of PARP-mediated programmed cell death, we applied 1 μM of the PARP inhibitor Olaparib (Gunderson and Moore, 2015) for 24 h followed by escalating doses of ara-C (1 – 15 μM) and idarubicin (0.06 – 0.9 μM) in a fixed, clinically-relevant mixture ratio of 17:1 (Dohner et al., 2010). Olaparib caused a statistically significant 18 – 36% increase in the number of living OCI-AML3 cells following treatment with ara-C/idarubicin as compared to cells receiving no Olaparib (Fig 2D). In contrast, Olaparib caused a small decrease in the number of viable OCI-AML2 cells when treated with the same mixture (Fig 2E). These results further support the conclusion that OCI-AML3 cells undergo parthanatos whereas OCI-AML2 cells undergo apoptosis upon addition of ara-C and idarubicin.

### PBMCs from healthy human donors exhibited parthanatos features upon treatment with ara-C and idarubicin ex vivo

Peripheral blood mononuclear cells (PBMCs) from healthy donors (n = 10) were treated with the PARP inhibitor Olaparib (1 μM, 24 h) prior to adding a clinically relevant 17:1 ratio mixture of ara-C and idarubicin for 24 h. The number of live cells in each sample was determined using a differential staining cytotoxicity assay (Perfetto et al., 2006) and flow cytometry. Olaparib pre-treatment of the primary PBMCs caused a 6.4 – 33% increase in the number of live cells as compared to ara-C and idarubicin treatment alone (Fig 3A, Supplementary Fig. A1-A5, Supplementary Tables S1-S2). Furthermore, all 10 samples of treated PBMCs from healthy donors exhibited the same ring-shaped nuclear morphologies (Fig. 3A, Supplementary Fig. A1-A5) as those observed for OCI-AML3 cells undergoing parthanatos (Fig. 2A,C). PBMCs from healthy donors treated with either ara-C or idarubicin alone also exhibited the same DNA ring morphologies (Supplementary Fig. B1-B2). To evaluate the specificity of this morphology-based analysis, five additional samples of PBMCs from healthy donors were treated with ara-C and/or idarubicin, or AT-101 for 24 h and subjected to DAPI staining and microscopy. Nuclear morphologies consistent with parthanatos were observed in all five of five PBMC samples upon addition of the ara-C/idarubicin mixture, as well as ara-C or idarubicin alone. In contrast, globular chromatin morphologies consistent with apoptosis were observed in all five of five samples treated with AT-101 (Supplementary Fig. B1-B2). These results demonstrate that PBMCs from healthy human donors exhibit two distinctive features of parthanatos when treated with a clinically relevant mixture of ara-C and idarubicin.

**Fig. 3.**
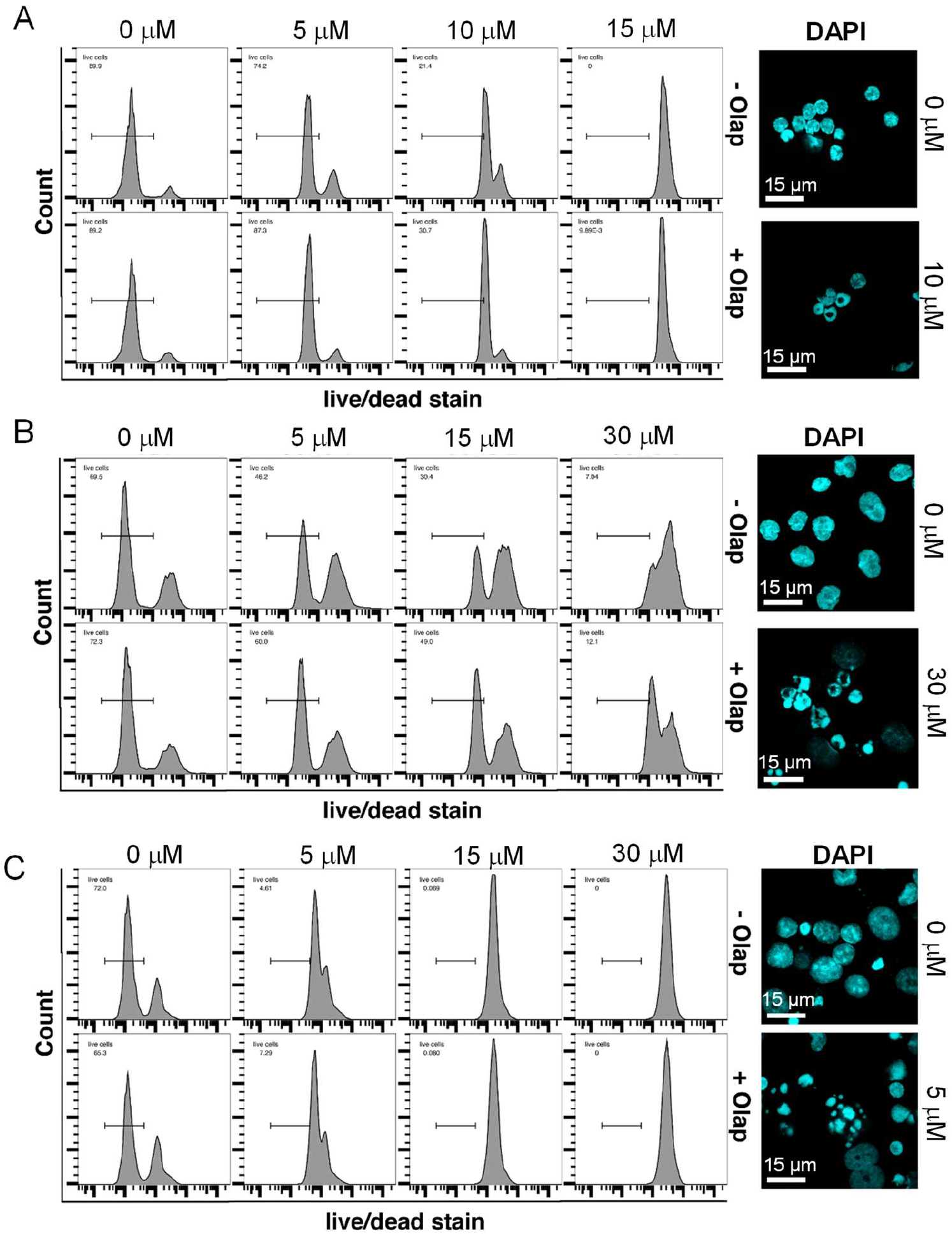
Analysis of primary AML blasts and healthy PBMCs for two parthanatos features upon treatment with ara-C and idarubicin ex vivo. **A)** PBMCs from healthy donors (10 of 10 tested) exhibited two distinct parthanatos features. The first feature is toxicity rescue according to a “live/dead” differential staining cytotoxicity assay (Perfetto et al., 2006) of cells treated with 1 μM of the PARP inhibitor Olaparib (Olap) for 24 hours prior to treatment with ara-C and idarubicin in a fixed 17:1 ratio for 24 hours. The second characteristic feature is the presence of ring-shaped nuclei according to DAPI staining and fluorescence microscopy (see Supplementary Fig. A1-A5 for all 10 examples). Semi-quantitative image analyses indicate the ring-shaped nuclear patterns were observed in 14 – 46% of the treated PBMCs, as compared to only 0.3 – 2.7 % of the untreated PBMCs. Cells treated with either ara-C or idarubicin alone also exhibited these characteristic DNA “ring” morphologies (Supplementary Fig. B1-B2). **B)** Primary cells from AML patient (ID 17-008) exhibiting both parthanatos features following treatment with a fixed, 17:1 ratio of ara-C and idarubicin for 24 hours (see Supplementary Fig. D1-D5 for all 18 examples). Concentrations are given in terms of ara-C. **C)** Primary cells from AML patient (ID 15-130) exhibiting neither parthanatos feature following treatment with ara-C and idarubicin for 24 hours. See Supplementary Fig. E1-E7 for all 21 examples of primary AML isolates exhibiting zero or one parthanatos feature.

### Parthanatos morphologies were observed in the nuclei of primary PBMCs from AML patients undergoing chemotherapy with ara-C and idarubicin

An evaluation of cell death characteristics during front-line AML chemotherapy in vivo was conducted by microscopic analysis of PBMCs collected from eight AML patients at 6 – 24 h after the onset of a standard “7 + 3” day continuous IV infusion of ara-C at 200 mg/m^2^ and idarubicin at 12 mg/m^2^ (Dohner et al., 2010). DAPI staining revealed characteristic ring chromatin patterns in samples from all eight AML patients tested (Supplementary Fig C). The patterns were consistent with those observed following ex vivo drug treatment of healthy PBMCs and primary AML cells (Fig. 3, Supplementary Figs A-B, D) as well as OCI-AML3 cell cultures undergoing parthanatos (Fig. 2). Other parthanatos assays, such as measuring PAR formation or the impact of PARP inhibition on drug toxicity were impractical for in vivo drug treatment studies. We therefore used primary AML isolates to assess the formation of chromatin ring structures and impact of PARP inhibition on drug sensitivity ex vivo.

### The presence of parthanatos features in primary AML isolates was associated with longer overall and event-free survival

All studies involving primary AML samples were conducted in a single-blind fashion, where all bioanalyses and assignments were conducted prior to the lead authors receiving any information about the diagnostic profile or clinical outcome of each patient (Supplementary Tables S3-S4). To facilitate comparisons with the results obtained using OCI-AML3 cells, primary cell samples from M4 and M5 FAB subgroup patients (acute myelomonocytic and acute monocytic leukemia) were included in this study due to their close morphological and cytochemical similarities (Bennett et al., 1985; Schiffer and Stone, 2003). Primary isolates from 39 AML patients were collected from blood (n = 33) or bone marrow (n = 6) at the time of diagnosis. The samples were purified using a Ficoll gradient and frozen in media containing 10% DMSO until use. To mimic in vivo growth conditions present in bone marrow, the thawed cells were co-cultured with a monolayer of human HS-5 stromal cells for 24 hours (Garrido et al., 2001). The co-cultured cells were treated with Olaparib (1 μM, 24 h) or carrier only (DMSO). On the second day after thawing, the cells were treated with a clinically-relevant mixture of ara-C (1 – 30 μM) and idarubicin (0.06 – 1.8 μM) in a fixed, 17:1 ratio for an additional 24 h. The samples were then analyzed for viability using a differential staining cytotoxicity assay (Perfetto et al., 2006) and flow cytometry. The AML cells were analytically distinguished from other cells using side light scattering and CD45 immunostaining (Garrido et al., 2001). In parallel, the same samples were analyzed for parthanatos-associated nuclear morphologies by DAPI staining and fluorescence microscopy. Approximately half of all AML patient samples (n = 18) exhibited both characteristic parthanatos features of toxicity rescue by Olaparib and ring-shaped nuclear morphologies (Fig 3B, Table 1, Supplementary Table S5, and Supplementary Fig D1-D9). The other half of AML patient samples (Supplementary Fig E1-E7) exhibited neither feature (n = 16, Fig 3C), or only one feature (n = 5). These results demonstrate a highly significant correlation (χ^2^_(3)_ = 21.8, *p* = 0.0001) between the presence/absence of toxicity rescue by Olaparib and the presence/absence of the DNA ring morphologies. Most of the parthanatos negative samples exhibited globular nuclear morphologies consistent with OCI-AML2 cells undergoing apoptosis (Supplementary Fig E1-E7). Given the ambiguous assignment of patients exhibiting only one parthanatos feature (n = 5), three separate survival analyses were conducted, where the 5 patients were considered to be parthanatos negative, parthanatos positive (Supplementary Table S6), or excluded from the analysis (Table 1). In all three analyses, the parthanatos positive subgroup exhibited a three-fold higher overall survival rate (Fig 4A, Supplementary Fig F, HR = 0.22 – 0.38, p = 0.002 – 0.05) and event-free survival (Supplementary Fig G, HR = 0.21 – 0.39; p = 0.001 – 0.05) as compared to the parthanatos negative group. The inclusion/exclusion of the four patients who were too weak to receive intensive chemotherapy (Supplementary Tables S4) did not impact these conclusions (Supplementary Figs. F-G). Notably, all four of these older patients (69 – 79 yo) were parthanatos negative according to both assays (Supplementary Tables S3,S5). Indeed, age and sex are general risk factors for AML (Acharya et al., 2018). Accordingly, the average age and masculinity of the parthanatos positive subgroup (53 years, 39 % male) were lower than those of the parthanatos negative subgroup (63 years, 75 % male, Table 1, Supplementary Table S6).

**Fig. 4.**
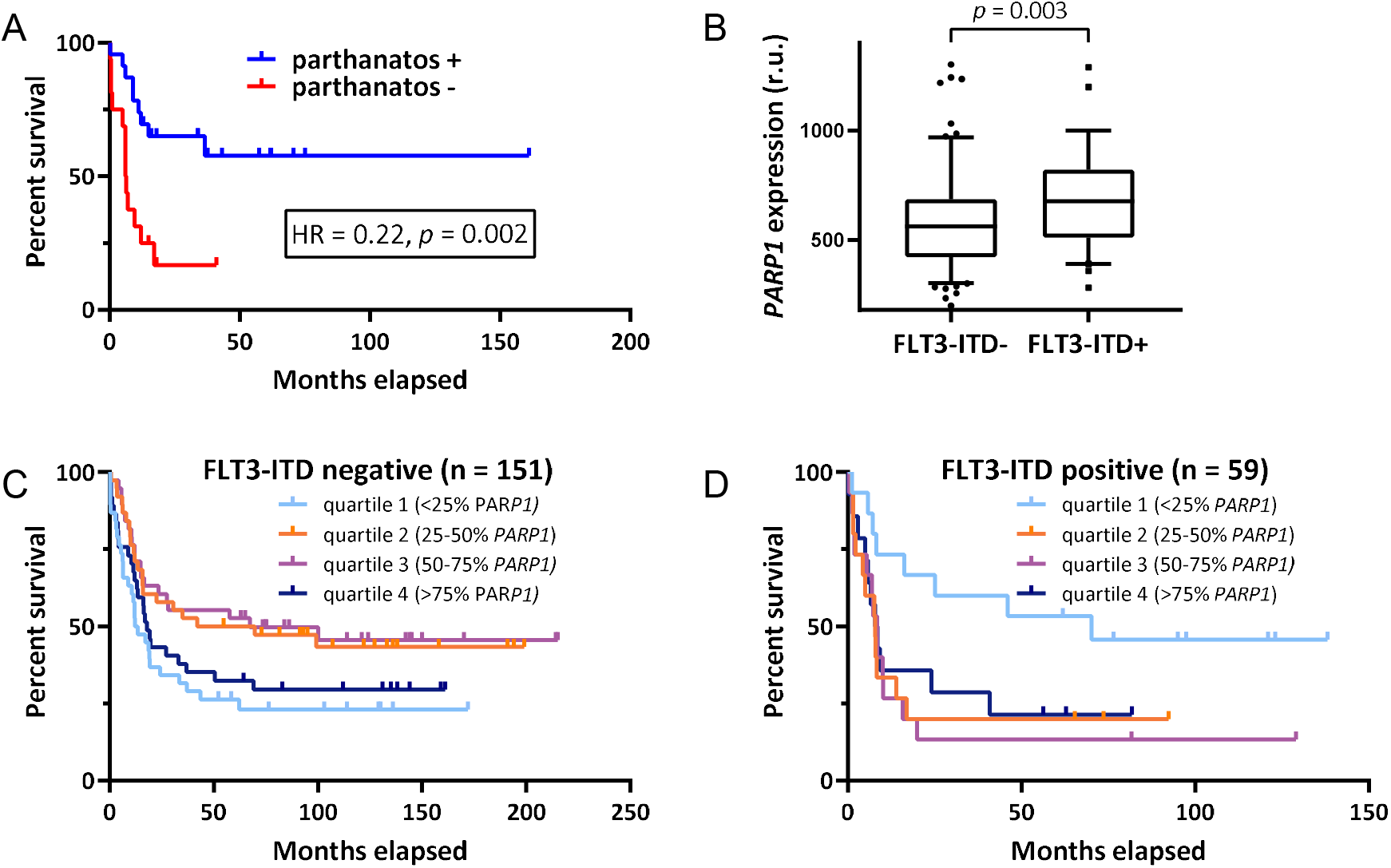
Kaplan-Meier curves of parthanatos positive (+) versus negative (−) groups as well as PARP1 mRNA expression versus survival. **A)** Overall % survival (OS) versus time of parthanatos +/- groups exhibiting both parthanatos features (n = 18) or neither feature (n = 16, Table 1). **B)** Boxplot with whiskers (25 – 75% interquartile range) illustrating higher relative *PARP1* mRNA levels in patients with a FLT3-ITD mutation (FLT3-ITD+) versus without (FLT3-ITD-). **C)** OS curves of FLT3-ITD negative M4 and M5 AML patients versus PARP1 mRNA quantities. Patients were grouped into four equal quartiles according to the relative expression of *PARP1* (quartile 1, <25%; quartile 2, 25-50%; quartile 3, 50-75%; and quartile 4, >75%) and OS were plotted using Kaplan-Meier survival estimates. **D)** OS of FLT3-ITD positive M4 and M5 AML patients versus relative expression of *PARP1*. See Supplementary Fig. H-I for analyses of *PARP2, PARG, ARH3*, inv(16) status, and NPM1 mutational status.

**Table 1.** Samples from 39 AML patients (FAB subtypes M4 and M5) were tested for the presence of the two parthanatos features: chromatin fragmentation “ring” morphologies and PARP-dependent changes in drug sensitivity upon addition of Olaparib. Only five of the 39 samples exhibited one feature in the absence of the other. The inclusion/exclusion of these five patients in survival analyses had no impact on the conclusions drawn in this study. See Supplementary Table S6 for alternative analyses including these five patients.

| Sample type | Total | Exhibited both parthanatos features | Exhibited one feature | Exhibited neither feature |
| --- | --- | --- | --- | --- |
| Blood | 33 | 15 | 4 | 14 |
| Bone marrow | 6 | 3 | 1 | 2 |
| Total | 39 | 18 | 5 | 16 |
|  |  | Positive | Excluded | Negative |
| Assigned patients (n) |  | 18 |  | 16 |
| Average age (years) |  | 53 ± 16 |  | 63 +/- 9 |
| Sex (% female) |  | 61 |  | 25 |
| FAB (% M4) |  | 50 |  | 50 |
| FLT3 positive (%) |  | 23 |  | 19 |
| Normal cytogenetics (%) |  | 45 |  | 63 |
| ELN Risk group | Adverse (%) | 33 |  | 31 |
|  | Intermediate (%) | 28 |  | 25 |
|  | Favorable (%) | 39 |  | 44 |
| Two-year overall survival (%) |  | 66 % |  | 17 % |
| Log-rank (Mantel-Cox) test: hazard ratio (HR) and statistical significance (p) |  | HR = 0.22, p = 0.002 |  |  |

### *PARP1* expression is a potential biomarker for intensive chemotherapy and the impact of FLT3-ITD status

Given the low frequency of *PARP1* mutations in hematopoietic and lymphoid blood cancers (41 of 5,212 tested (Sanger, 2021)), together with the ability of PARP inhibition to decrease AML drug sensitivity (Supplementary Table S5), we hypothesized that differing levels of *PARP1* expression are associated with AML survival. We therefore investigated *PARP1* expression as a predictive biomarker of chemotherapy response in AML patients by analyzing microarray data from clinical trials conducted by the Haemato Oncology Foundation for Adults in the Netherlands (Walter et al., 2015). For direct comparison purposes, mRNA expression data from AML patients with FAB subtypes M4 and M5 (n = 210) receiving curative treatment including “7 + 3 induction” therapy with ara-C and idarubicin were stratified into four equal quartile groups based on the relative mRNA expression (quartile 1: <25%; quartile 2: 25-50%; quartile 3: 50-75%; and quartile 4: >75%) of genes associated with parthanatos including *PARP1, PARP2, PARG*, and *ARH3*. Kaplan-Meier survival curves were analyzed using a log-rank (Mantel-Cox) test. No significant differences in overall survival (OS) were observed between any of the quartile groups for *PARP2, PARG*, or *ARH3*. In contrast, patients with near-median expression levels (quartiles 2 and 3) of *PARP1* exhibited a 50% improved OS rate (HR = 0.66, *p* = 0.01) as compared to patients with high (quartile 4) or low (quartile 1) *PARP1* expression (Supplementary Fig. H). Further evidence that dysregulated *PARP1* expression is associated with poor outcomes was obtained by evaluating its co-occurrence with the common, poor-prognosis mutational biomarker FLT3-ITD that is present in ~25% of AML cases (Stirewalt and Radich, 2003). The FLT3-ITD positive patients exhibited a significantly higher relative expression of *PARP1* as compared to the FLT3-ITD negative patients (Fig 4B). In contrast, *PARP1* expression was found to be independent of other common biomarkers including inv(16) translocation and mutation of *NPM1* (Supplementary Fig. I). The FLT3-ITD negative patients (n = 151) with near-median *PARP1* expression (quartiles 2 and 3) exhibited nearly a 2-fold improved overall survival rate as compared to quartiles 1 and 4 (HR = 0.55, *p* = 0.003, Fig. 4C). In contrast, the FLT3-ITD positive patients (n = 59) exhibited a different pattern, where low expression of *PARP1* (quartile 1) was associated with a >two-fold higher OS rate (HR = 0.45, *p* = 0.013) as compared to quartiles 2-4 (Fig. 4D). This latter pattern has been reported in previous studies and has motivated clinical development of PARP-1 inhibitors (Gil-Kulik et al., 2020; Li et al., 2018; Pashaiefar et al., 2018). Together these results suggest that *PARP1* expression is an important prognostic biomarker for M4 and M5 AML subtypes, and further support the relevance of parthanatos in curative chemotherapy for AML.

## DISCUSSION

For nearly 50 years, chemotherapy-induced cell death mechanistic studies and associated design of new cancer drugs have focused almost exclusively on the induction of apoptosis (Carneiro and El-Deiry, 2020). The translational importance of such studies is evidenced by the development of new drugs based on Bcl-2 family inhibitors such as Venetoclax (DiNardo et al., 2019). We hypothesized that non-apoptotic form(s) of programmed death play a role in the treatment of AML due to previous observations that primary AML cells treated with ara-C can undergo caspase-independent programmed cell death (Carter et al., 2003) as well as caspase-3 activation failing to predict ara-C chemosensitivity (Staib et al., 2005). These observations were reported before parthanatos was recognized as a distinct form of programmed cell death (Galluzzi et al., 2012).

Here, we report that a therapeutically-relevant, front-line AML regimen of ara-C and an anthracycline causes characteristic features of parthanatos in samples from approximately 50% of AML patients (FAB groups M4 and M5). The presence of one or more parthanatos features is associated with four-fold improved AML survival rates following curative chemotherapy (HR = 0.24, p = 0.008, Supplementary Fig F). Surprisingly, the primary isolates from the parthanatos positive subgroup exhibited the same or even lower average drug sensitivity ex vivo (EC_50_ = 13 ± 8 μM (- Olaparib), EC_50_ = 21 ± 14 μM (+Olaparib)) as compared to the parthanatos negative subgroup (EC_50_ = 11 ± 6 μM (-Olaparib), EC_50_ = 11 ± 4 μM (+Olaparib), Supplementary Table S5). These results demonstrate that the mechanism of cell death can have more prognostic value than the absolute drug sensitivity exhibited by primary isolates ex vivo.

In this study, we focused on AML FAB subgroups M4 + M5 since they are biologically distinct from the other FAB subgroups, and together these represent approximately 35% of all AML cases (Bennett et al., 1985; Schiffer and Stone, 2003). Depending on the exact cell line being studied, a clinically relevant 17:1 mixture of ara-C and idarubicin (Dohner et al., 2010) induced either parthanatos or apoptosis. Our results with OCI-AML2 cells are consistent with previous studies (Deng et al., 2017; Vincelette and Yun, 2014) showing that certain leukemia cell lines can undergo classical apoptosis following ara-C treatment. However, the closely related cell line OCI-AML3 exhibited parthanatos features under the same conditions. These features included a lack of caspase-3 activation, PAR accumulation (Andrabi et al., 2006), AIF translocation (Yu et al., 2006), and the production of large DNA fragments at the nuclear periphery. Interestingly, we also observed a lack of mitochondrial transition pores and hyperpolarization of mitochondrial membranes (Fig. 1C-D). These two features have not been previously reported during PARP-1-mediated programmed cell death, and they are in stark contrast to the opening of mitochondrial pores and loss of mitochondrial membrane potentials associated with apoptosis. Of note, the parthanatos we observed in OCI-AML3 cells was a specific result of ara-C and idarubicin because the addition of camptothecin or AT-101 caused apoptosis features including caspase 3 activation and globular nuclear morphologies (Fig. 1B, Fig. 2C). Likewise, PMBCs from healthy human donors also exhibited either parthanatos or apoptosis features upon addition of ara-C/idarubicin or AT-101, respectively (Supplementary Fig. B1-B2). Together with the low (~30%) overall cure rate of the disease, these results lead to our hypothesis that the presence of different programmed cell death pathways in individual AML patients is associated with different clinical outcomes following intensive chemotherapy with ara-C and idarubicin. To test this hypothesis, we evaluated different assays capable of identifying parthanatos features using a relatively small number of primary AML cells that survive for only a few days in culture. We found that one such feature is a distinct nuclear morphology where the fragments of cleaved chromatin are distributed in a ring-shaped pattern (Fig. 2C). This pattern is likely a result of MIF cleavage of single-stranded DNA (Wang et al., 2016) into a relatively small number of large chromatin fragments as confirmed here by the TUNEL assay (Fig. 2B). Our results are consistent with an early report of DNA fragmentation patterns of primary leukemia cells treated with various nucleoside-based drugs that failed to give apoptotic nucleosomal ladders, but rather large molecular weight DNA fragments (Huang et al., 1995). The nuclei of monocytes and their precursors (including M4 and M5 AML) are known to exhibit highly distinct nuclear morphologies (Skinner and Johnson, 2017) that may expose MIF-hypersensitive cleavage sites to give large chromatin fragments arranged in a sphere at the periphery of the nucleus. Our findings are further supported by previous studies of human breast epithelial cell lines MCF-10A and SKBR-3 undergoing parthanatos which also exhibited DNA fragments distributed in a characteristic ring-shaped pattern (Soriano et al., 2017).

Primary white blood cells from all healthy human donors (10 of 10 tested) and approximately 50% of M4/M5 AML (18 of 39 tested) exhibited ring-like nuclear morphologies as well as drug resistance upon inhibition of PARP. AML patients with primary isolates exhibiting one or both of these characteristic parthanatos features had an overall two-year survival of 66 – 67% in contrast to the 17 – 28% survival of the parthanatos negative subgroup (Table 1, Supplementary Table S6). The parthanatos positive and negative subgroups contained members from a wide variety of risk groups with diverse cytogenetic markers. Surprisingly, the ELN risk group distribution (% adverse, % intermediate, and % favorable) of the parthanatos positive and negative subgroups was essentially identical. These results suggest that the presence/absence of parthanatos features has prognostic value independent of the ELN risk assessment criteria. The presence/absence of Olaparib rescue and the presence/absence of the nuclear DNA ring morphologies for the 39 AML isolates were highly correlated with one another (χ^2^_(3)_ = 21.8, *p* = 0.0001), and thus would serve as robust ex vivo prognostic assay for cytarabine administration in the clinic. In the future, newly diagnosed AML patients who are predicted to have poor responsiveness towards intensive chemotherapy with cytarabine will be prescribed alternative therapies such as a hypomethylating agent (De Kouchkovsky and Abdul-Hay, 2016; Dombret et al., 2015), Bcl-2 inhibitor (DiNardo et al., 2019), and/or FLT3 kinase inhibitor (Antar et al., 2020). Improved AML patient stratification is especially important given the devastating side effects of intensive chemotherapy with ara-C including “tumor lysis syndrome” in ~20 % of treated patients that is responsible for approximately 2 – 5 % of AML mortality (Montesinos et al., 2008). Such treatment-related mortality further highlights the heterogeneity of the disease and the need for new diagnostic strategies to identify which patients are not suitable candidates for intensive “7 + 3” chemotherapy.

With a general trend towards whole transcriptome RNA sequencing (Arindrarto et al., 2020) and other high-throughput biomarker discovery (Bai et al., 2013; Drenberg et al., 2019; Prashad et al., 2016; Saad et al., 2018), the profiling of genes such as *PARP1* that is associated with cytarabine-induced parthanatos may provide a practical means for patient stratification. Indeed, PARP-1 is known to play a central role in parthanatos, DNA damage repair, and hematopoietic differentiation (Gil-Kulik et al., 2020; Hsieh et al., 2017). Previous studies reported that *PARP1* overexpression predicts poor AML patient survival (Gil-Kulik et al., 2020; Li et al., 2018; Pashaiefar et al., 2018). Here we observed the same trend for FLT3-ITD positive AML (Fig. 4D) that is a well-known prognostic marker associated with poor survival in ~25% of AML cases (Stirewalt and Radich, 2003). Our results further support the application of PARP inhibitors in FLT3 mutated AML, particularly in combination with synthetic lethal partners such as specific genetic alterations (Dellomo et al., 2019). Surprisingly, our results revealed that abnormally low expression of *PARP1* in FLT3-ITD negative AML (~75% of cases) is associated with poor survival (Fig. 4C). Taken together with the antagonistic effects of Olaparib towards ara-C and idarubicin in primary AML cancer cells, it is possible that low levels of PARP1 expression enable drug resistance by evasion of parthanatos. On the other hand, abnormally high levels of PARP-1 may result in over-active repair and thus resistance to DNA-damaging chemotherapy. This “double-edged sword” of PARP-1 activity has been previously noted with respect to carcinogenesis (Dorsam et al., 2018). Here we report this type of dual activity is relevant to AML treatment stratification. In addition to providing a means for specifically identifying which patients are suitable candidates for intensive “7 + 3” chemotherapy, our results contraindicate the use of PARP inhibitors in FLT3 wild-type AML patients receiving ara-C. In contrast, the addition of a PARP1 inhibitor to the 7 + 3 regimen should be further evaluated for the treatment of FLT3 mutated AML.

## MATERIALS AND METHODS

### Ethical approval

This study was approved by a decision of the local ethics committee in Berne, Switzerland (decision number #207/14), and Zurich, Switzerland (EK-ZH-NR: 2009-0062/1 and BASEC-NR: 2018-00539). All donors provided written, informed consent.

### Primary cell collection

Mononuclear cells were isolated from the peripheral blood of AML patients at the time of diagnosis or from healthy donors using a Lymphoprep^™^ (Axis-Shield) and were frozen in media with 10% DMSO as a cryoprotectant and stored in liquid nitrogen. AML cells were purified from the bone marrow using Ficoll density gradient centrifugation and cryopreserved in media containing 10% fetal calf serum (FCS) and 10% DMSO.

### Cell culture

OCI-AML3 and OCI-AML2 cell lines were obtained from the DSMZ and monthly tested for mycoplasma infections using the MycoFluor^™^ Mycoplasma Detection Kit (Invitrogen). AML cell lines and primary cells were cultured in RPMI at 37°C in a humidified incubator with 5 % CO2. The media was supplemented with 1 % MEM non-essential amino acid solution (Sigma-Aldrich), 50,000 units of penicillin and 50 mg of streptomycin per liter (Sigma-Aldrich). Additionally, 10 % FBS (Thermo Fisher Scientific) was added to the media for OCI-AML2 cells while 20 % was supplemented for OCI-AML3, HS-5, and primary cells. Cell lines were grown to confluency and passaged every 2 – 4 days. A Scepter cell counter (Merck Millipore) was used for cell counting. Primary cell samples were thawed and transferred to a 50 ml tube containing 200 μl of PBS/DNase (2000 U/ml) solution (Sigma-Aldrich). The cryotube was rinsed with 1 ml of thawing medium (5 % FBS/PBS/DNase (20 U/ml)). Afterwards, 18 ml of thawing media was added in a stepwise manner over 4 min. The cells were centrifuged for 15 min at 300 x g and resuspended in 5 ml media. Cell viability was determined with Trypan Blue staining (Sigma-Aldrich) and analyzed by Cell Counter Model R1 (Olympus). For the Olaparib rescue experiments, blood cells were co-cultured with the feeder cell line HS-5 (Garrido et al., 2001). Accordingly, the HS-5 cells were seeded in 24-well plates at a density of 1 x 10^5^ / 500 μl media per well 24 h prior to seeding of the primary blood cells at a density of 2 – 6 x 10^5^ / 500 μl media per well.

### Reagents

Cytosine β-D-arabinofuranoside (cytarabine, ara-C), idarubicin, AT101, camptothecin (CPT), 4’,6-diamidino-2-phenylindole (DAPI), and dimethyl sulfoxide DMSO were purchased from Sigma-Aldrich. Olaparib was obtained from Selleck Chemicals. Primary antibodies used: monoclonal rabbit anti-AIF antibody (1:50, Abcam), monoclonal mouse APC CD45 conjugate (1:50, BioLegend), monoclonal mouse anti-PAR antibody (10H, 1:300, Adipogen). Secondary antibodies used: goat-anti-mouse Alexa Fluor 488 antibody conjugate (1:100, Thermo Fisher Scientific), goat-anti-rabbit Alexa Fluor 555 antibody conjugate (1:50, Thermo Fisher Scientific).

### Differential staining cytotoxicity assay (Perfetto et al., 2006)

Cells were seeded in 6-well plates at a density of 5 x 10^5^ / 2 ml media per well. After incubation with relevant doses of drugs, cells were transferred to centrifuge tubes, washed with PBS, and stained for live and dead cells with the LIVE/DEAD fixable green dead cell stain kit (Thermo Fisher Scientific) according to the instructions of the manufacturer. In the case of primary AML isolates, the cells were stained at the same time with the APC conjugated surface marker antibody CD45 after a 5 min blocking step with Human TruStain FcX solution (20x, BioLegend) diluted in 1 % BSA/PBS. Fixation was performed in 300 μl paraformaldehyde (cell lines: 3.7 % in PBS, primary cells: 2 % in PBS) for 15 min at RT. Cells were washed with PBS and 1 % BSA/PBS. As a next step, cells were permeabilized in 200 μl 0.2 % Triton-X-100/PBS for 15 min on ice and washed with PBS. Afterwards, cells were blocked with 1 % BSA/PBS for 15 min at RT. Primary antibodies were diluted in 1 % BSA/PBS to their appropriate concentrations and cells were incubated with the solution for 2 h at RT. Cells were washed with PBS, 1 % BSA/PBS and finally incubated in 50 μl 1 % BSA/PBS containing the appropriate secondary antibodies for 30 min at RT. After two more washing steps with PBS, the cells were resuspended in 500 μl DAPI solution (5 μM in PBS) and incubated overnight at 4°C. Cell suspensions were analyzed by LSR II Fortessa (BD Biosciences) and the results were evaluated with the FlowJo software (version 10.0.8, FlowJo LLC).

### Cell imaging

For further analysis with confocal microscopy, the cells were transferred to microscope slides using a Thermo Shandon Cytospin 3 Centrifuge, coverslipped with ProLong Gold Antifade mountant (Thermo Fisher Scientific), and analyzed by confocal laser scanning microscopy using a Leica SP5 Mid UV-VIS or TCS Leica SP8 Multiphoton (63x oil immersion objective, NA 1.4). Images were analyzed with LAS AF 2.6.0 (Leica Microsystems) and ImageJ 1.47c (National Institutes of Health).

### Annexin V staining

Cells were seeded in 6-well plates at a density of 1.2 x 10^6^ / 2.5 ml media per well. After 24 h incubation with relevant doses of drugs, phosphatidyl serine externalization was determined with Dead Cell Apoptosis Kit with Annexin V Alexa Fluor 488 and propidium iodide (Thermo Fisher Scientific) according to the instructions of the manufacturer.

### Caspase-3 Assay

Cells were seeded in 6-well plates at a density of 5 x 10^5^ / 2 ml media per well and incubated with relevant doses of drugs for 8 h. Activity of caspase-3 was detected with NucView 488 Caspase-3 Assay kit (Biotium) according to the instructions of the manufacturer.

### Mitochondrial Membrane Potential Assay

Cells were seeded in 6-well plates at a density of 5 x 10^5^ / 2 ml media per well and incubated with relevant doses of drugs for 24 h. Cells were transferred to centrifuge tubes and stained with 500 μl Mitotracker Red CMXRos staining solution (200 nM in PBS, Thermo Fisher Scientific) for 30 min at 37°C. After one washing step with PBS, cells were fixed with 300 μl paraformaldehyde (3.7 % in PBS) for 15 min at RT. The cells were washed twice with PBS and resuspended in 500 μl DAPI solution (5 μM in PBS) and incubated overnight at 4°C. Cell suspensions were analyzed by LSR II Fortessa.

### MitoProbe Transition Pore Assay

Cells were seeded in 6-well plates at a density of 5 x 10^5^ / 2 ml media per well and incubated with relevant doses of drugs for 24 h. Mitochondrial permeability transition pore opening was measured using MitoProbe Transition Pore Assay Kit (Thermo Fisher Scientific) according to the instructions of the manufacturer.

### TUNEL Assay

Cells were seeded in 6-well plates at a density of 5 x 10^5^ / 2 ml media per well and incubated with relevant doses of drugs for 24 h. The CF^™^ 488 TUNEL Assay (Biotium) was performed according to the instructions of the manufacturer.

### Statistical analysis

Kaplan-Meier survival curves were analyzed by a log-rank (Mantel-Cox) test and the hazard ratios (Mantel-Haenszel).

## SUPPORTING INFORMATION

The Supporting Information is available free of charge. This material includes anonymized patient information, microscopy, flow cytometry, EFS, and RNA expression data.

## Acknowledgements

We thank Dr. François Mercier, Dr. Paul MacEoin and Dr. Therese Trimer for helpful discussions. We thank Kaixiang Wang, Joshua O’Grady, and Ayodele Edinboro for proofreading. We thank Dr. Urs Ziegler and Dr. Nicolas Audet for technical assistance with microscopy, and Dr. Claudia Dumrese for technical assistance with flow cytometry. We acknowledge the Swiss National Science Foundation (grant 165949 to N.W.L.), the Natural Sciences and Engineering Research Council of Canada (grant 05048 to N.W.L.), and CFI John R. Evans Leaders Fund (grant 39168 to N.W.L.) for funding. A.P.A.T. is supported by the Professor Dr. Max Cloëtta foundation.

## Author Contributions

N.W.L. conceived the project and together with A.M., T.P., K.S. and M.M. designed the experiments; A.M., B.M. and K.S. performed the experiments; T.P., A.P.A.T., and P.J.M.V. provided human cell samples; P.J.M.V. provided the microarray data; N.W.L., A.M., B.M. and M. M. analyzed the data, prepared the figures, and wrote the manuscript.

## References

Acharya, U.H., Halpern, A.B., Wu, Q.V., Voutsinas, J.M., Walter, R.B., Yun, S., Kanaan, M., and Estey, E.H. (2018). Impact of region of diagnosis, ethnicity, age, and gender on survival in acute myeloid leukemia (AML). J Drug Assess 7, 51–53.

Andrabi, S.A., Dawson, T.M., and Dawson, V.L. (2008). Mitochondrial and nuclear cross talk in cell death: parthanatos. Annals of the New York Academy of Sciences 1147, 233–241.

Andrabi, S.A., Kim, N.S., Yu, S.W., Wang, H., Koh, D.W., Sasaki, M., Klaus, J.A., Otsuka, T., Zhang, Z., Koehler, R.C., et al. (2006). Poly(ADP-ribose) (PAR) polymer is a death signal. Proceedings of the National Academy of Sciences of the United States of America 103, 18308–18313.

Antar, A.I., Otrock, Z.K., Jabbour, E., Mohty, M., and Bazarbachi, A. (2020). FLT3 inhibitors in acute myeloid leukemia: ten frequently asked questions. Leukemia 34, 682–696.

Arindrarto, W., Borras, D.M., de Groen, R.A.L., van den Berg, R.R., Locher, I.J., van Diessen, S., van der Holst, R., van der Meijden, E.D., Honders, M.W., de Leeuw, R.H., et al. (2020). Comprehensive diagnostics of acute myeloid leukemia by whole transcriptome RNA sequencing. Leukemia 35, 47–61.

Auberger, P., and Puissant, A. (2017). Autophagy, a key mechanism of oncogenesis and resistance in leukemia. Blood 129, 547–552.

Bai, J., He, A., Zhang, W., Huang, C., Yang, J., Yang, Y., Wang, J., and Zhang, Y. (2013). Potential biomarkers for adult acute myeloid leukemia minimal residual disease assessment searched by serum peptidome profiling. Proteome science 11, 39.

Bennett, J.M., Catovsky, D., Daniel, M.T., Flandrin, G., Galton, D.A., Gralnick, H.R., and Sultan, C. (1985). Proposed revised criteria for the classification of acute myeloid leukemia. A report of the French-American-British Cooperative Group. Ann Intern Med 103, 620–625.

Carneiro, B.A., and El-Deiry, W.S. (2020). Targeting apoptosis in cancer therapy. Nat Rev Clin Oncol 17, 395–417.

Carter, B.Z., Kornblau, S.M., Tsao, T., Wang, R.Y., Schober, W.D., Milella, M., Sung, H.G., Reed, J.C., and Andreeff, M. (2003). Caspase-independent cell death in AML: caspase inhibition in vitro with pancaspase inhibitors or in vivo by XIAP or Survivin does not affect cell survival or prognosis. Blood 102, 4179–4186.

Cassier, P.A., Castets, M., Belhabri, A., and Vey, N. (2017). Targeting apoptosis in acute myeloid leukaemia. Br J Cancer 117, 1089–1098.

Cicconi, L., and Lo-Coco, F. (2016). Current management of newly diagnosed acute promyelocytic leukemia. Annals of oncology: official journal of the European Society for Medical Oncology 27, 1474–1481.

Cloux, A.J., Aubry, D., Heulot, M., Widmann, C., ElMokh, O., Piacente, F., Cea, M., Nencioni, A., Bellotti, A., Bouzourene, K., et al. (2019). Reactive oxygen/nitrogen species contribute substantially to the antileukemia effect of APO866, a NAD lowering agent. Oncotarget 10, 6723–6738.

Costa, A.F.O., Menezes, D.L., Pinheiro, L.H.S., Sandes, A.F., Nunes, M.A.P., Lyra Junior, D.P., and Schimieguel, D.M. (2017). Role of new Immunophenotypic Markers on Prognostic and Overall Survival of Acute Myeloid Leukemia: a Systematic Review and Meta-Analysis. Scientific reports 7, 4138.

David, K.K., Andrabi, S.A., Dawson, T.M., and Dawson, V.L. (2009). Parthanatos, a messenger of death. Frontiers in bioscience 14, 1116–1128.

De Kouchkovsky, I., and Abdul-Hay, M. (2016). ‘Acute myeloid leukemia: a comprehensive review and 2016 update’. Blood cancer journal 6, e441.

Dellomo, A.J., Baer, M.R., and Rassool, F.V. (2019). Partnering with PARP inhibitors in acute myeloid leukemia with FLT3-ITD. Cancer Lett 454, 171–178.

Deng, R., Fan, F.Y., Yi, H., Fu, L., Zeng, Y., Wang, Y., Miao, X.J., Shuai, Y.R., He, G.C., and Su, Y. (2017). Cytotoxic T lymphocytes promote cytarabine-induced acute myeloid leukemia cell apoptosis via inhibiting Bcl-2 expression. Experimental and therapeutic medicine 14, 1081–1085.

DiNardo, C.D., Pratz, K., Pullarkat, V., Jonas, B.A., Arellano, M., Becker, P.S., Frankfurt, O., Konopleva, M., Wei, A.H., Kantarjian, H.M., et al. (2019). Venetoclax combined with decitabine or azacitidine in treatment-naive, elderly patients with acute myeloid leukemia. Blood 133, 7–17.

Dohner, H., Estey, E.H., Amadori, S., Appelbaum, F.R., Buchner, T., Burnett, A.K., Dombret, H., Fenaux, P., Grimwade, D., Larson, R.A., et al. (2010). Diagnosis and management of acute myeloid leukemia in adults: recommendations from an international expert panel, on behalf of the European LeukemiaNet. Blood 115, 453–474.

Dombret, H., Seymour, J.F., Butrym, A., Wierzbowska, A., Selleslag, D., Jang, J.H., Kumar, R., Cavenagh, J., Schuh, A.C., Candoni, A., et al. (2015). International phase 3 study of azacitidine vs conventional care regimens in older patients with newly diagnosed AML with >30% blasts. Blood 126, 291–299.

Dorsam, B., Seiwert, N., Foersch, S., Stroh, S., Nagel, G., Begaliew, D., Diehl, E., Kraus, A., McKeague, M., Minneker, V., et al. (2018). PARP-1 protects against colorectal tumor induction, but promotes inflammation-driven colorectal tumor progression. Proceedings of the National Academy of Sciences of the United States of America 115, E4061–e4070.

Drenberg, C.D., Shelat, A., Dang, J., Cotton, A., Orwick, S.J., Li, M., Jeon, J.Y., Fu, Q., Buelow, D.R., Pioso, M., et al. (2019). A high-throughput screen indicates gemcitabine and JAK inhibitors may be useful for treating pediatric AML. Nat Commun 10, 2189.

Estey, E.H. (2018). Acute myeloid leukemia: 2019 update on risk-stratification and management. Am J Hematol 93, 1267–1291.

Fatokun, A.A., Dawson, V.L., and Dawson, T.M. (2014). Parthanatos: mitochondrial-linked mechanisms and therapeutic opportunities. British journal of pharmacology 171, 2000–2016.

Galluzzi, L., Vitale, I., Aaronson, S.A., Abrams, J.M., Adam, D., Agostinis, P., Alnemri, E.S., Altucci, L., Amelio, I., Andrews, D.W., et al. (2018). Molecular mechanisms of cell death: recommendations of the Nomenclature Committee on Cell Death 2018. Cell Death Differ 25, 486–541.

Galluzzi, L., Vitale, I., Abrams, J.M., Alnemri, E.S., Baehrecke, E.H., Blagosklonny, M.V., Dawson, T.M., Dawson, V.L., El-Deiry, W.S., Fulda, S., et al. (2012). Molecular definitions of cell death subroutines: recommendations of the Nomenclature Committee on Cell Death 2012. Cell Death Differ 19, 107–120.

Garrido, S.M., Appelbaum, F.R., Willman, C.L., and Banker, D.E. (2001). Acute myeloid leukemia cells are protected from spontaneous and drug-induced apoptosis by direct contact with a human bone marrow stromal cell line (HS-5). Exp Hematol 29, 448–457.

Gil-Kulik, P., Dudzinska, E., Radzikowska-Buchner, E., Wawer, J., Jojczuk, M., Nogalski, A., Wawer, G.A., Feldo, M., Kocki, W., Cioch, M., et al. (2020). Different regulation of PARP1, PARP2, PARP3 and TRPM2 genes expression in acute myeloid leukemia cells. BMC Cancer 20, 435.

Gunderson, C.C., and Moore, K.N. (2015). Olaparib: an oral PARP-1 and PARP-2 inhibitor with promising activity in ovarian cancer. Future oncology 11, 747–757.

Harrison, J.S., Wang, X., and Studzinski, G.P. (2016). The role of VDR and BIM in potentiation of cytarabine-induced cell death in human AML blasts. Oncotarget 7, 36447–36460.

Houl, J.H., Ye, Z., Brosey, C.A., Balapiti-Modarage, L.P.F., Namjoshi, S., Bacolla, A., Laverty, D., Walker, B.L., Pourfarjam, Y., Warden, L.S., et al. (2019). Selective small molecule PARG inhibitor causes replication fork stalling and cancer cell death. Nat Commun 10, 5654.

Hsieh, M.-H., Chen, Y.-T., Chen, Y.-T., Lee, Y.-H., Lu, J., Chien, C.-L., Chen, H.-F., Ho, H.-N., Yu, C.-J., Wang, Z.-Q., et al. (2017). PARP1 controls KLF4-mediated telomerase expression in stem cells and cancer cells. Nucleic Acids Research 45, 10492–10503.

Huang, P., Robertson, L.E., Wright, S., and Plunkett, W. (1995). High molecular weight DNA fragmentation: a critical event in nucleoside analogue-induced apoptosis in leukemia cells. Clinical cancer research: an official journal of the American Association for Cancer Research 1, 1005–1013.

Huang, X., Xiao, F., Li, Y., Qian, W., Ding, W., and Ye, X. (2018). Bypassing drug resistance by triggering necroptosis: recent advances in mechanisms and its therapeutic exploitation in leukemia. J Exp Clin Cancer Res 37, 310.

Jonas, B.A., and Pollyea, D.A. (2019). How we use venetoclax with hypomethylating agents for the treatment of newly diagnosed patients with acute myeloid leukemia. Leukemia 33, 2795–2804.

Kajiume, T., and Kobayashi, M. (2018). Human granulocytes undergo cell death via autophagy. Cell Death Discov 4, 111.

Kandoth, C., McLellan, M.D., Vandin, F., Ye, K., Niu, B., Lu, C., Xie, M., Zhang, Q., McMichael, J.F., Wyczalkowski, M.A., et al. (2013). Mutational landscape and significance across 12 major cancer types. Nature 502, 333–339.

Kopmar, N.E., and Estey, E.H. (2019). New drug approvals in acute myeloid leukemia: an unprecedented paradigm shift. Clin Adv Hematol Oncol 17, 569–575.

Leong, S.M., Tan, B.X., Bte Ahmad, B., Yan, T., Chee, L.Y., Ang, S.T., Tay, K.G., Koh, L.P., Yeoh, A.E., Koay, E.S., et al. (2010). Mutant nucleophosmin deregulates cell death and myeloid differentiation through excessive caspase-6 and −8 inhibition. Blood 116, 3286–3296.

Leung, A.K. (2014). Poly(ADP-ribose): an organizer of cellular architecture. The Journal of cell biology 205, 613–619.

Li, X., Li, C., Jin, J., Wang, J., Huang, J., Ma, Z., Huang, X., He, X., Zhou, Y., Xu, Y., et al. (2018). High PARP-1 expression predicts poor survival in acute myeloid leukemia and PARP-1 inhibitor and SAHA-bendamustine hybrid inhibitor combination treatment synergistically enhances anti-tumor effects. EBioMedicine 38, 47–56.

Mashimo, M., Kato, J., and Moss, J. (2013). ADP-ribosyl-acceptor hydrolase 3 regulates poly (ADP-ribose) degradation and cell death during oxidative stress. Proceedings of the National Academy of Sciences of the United States of America 110, 18964–18969.

Messikommer, A., Seipel, M., Byrne, S., Valk, P.J.M., Pabst, T., and Luedtke, N.W. (2020). RNA targeting in acute myeloid leukemia. ACS Pharmacol Transl Sci 3, 1225–1232.

Mezzatesta, C., and Bornhauser, B.C. (2019). Exploiting Necroptosis for Therapy of Acute Lymphoblastic Leukemia. Front Cell Dev Biol 7, 40.

Mohammad, R.M., Muqbil, I., Lowe, L., Yedjou, C., Hsu, H.Y., Lin, L.T., Siegelin, M.D., Fimognari, C., Kumar, N.B., Dou, Q.P., et al. (2015). Broad targeting of resistance to apoptosis in cancer. Semin Cancer Biol 35 Suppl, S78–S103.

Montesinos, P., Lorenzo, I., Martin, G., Sanz, J., Perez-Sirvent, M.L., Martinez, D., Orti, G., Algarra, L., Martinez, J., Moscardo, F., et al. (2008). Tumor lysis syndrome in patients with acute myeloid leukemia: identification of risk factors and development of a predictive model. Haematologica 93, 67–74.

Pan, R., Hogdal, L.J., Benito, J.M., Bucci, D., Han, L., Borthakur, G., Cortes, J., DeAngelo, D.J., Debose, L., Mu, H., et al. (2014). Selective BCL-2 inhibition by ABT-199 causes on-target cell death in acute myeloid leukemia. Cancer discovery 4, 362–375.

Pan, R., Ruvolo, V., Mu, H., Leverson, J.D., Nichols, G., Reed, J.C., Konopleva, M., and Andreeff, M. (2017). Synthetic Lethality of Combined Bcl-2 Inhibition and p53 Activation in AML: Mechanisms and Superior Antileukemic Efficacy. Cancer Cell 32, 748–760 e746.

Pashaiefar, H., Yaghmaie, M., Tavakkoly-Bazzaz, J., Ghaffari, S.H., Alimoghaddam, K., Momeny, M., Izadi, P., Izadifard, M., Kasaeian, A., and Ghavamzadeh, A. (2018). PARP-1 Overexpression as an Independent Prognostic Factor in Adult Non-M3 Acute Myeloid Leukemia. Genetic testing and molecular biomarkers 22, 343–349.

Paulus, A., Chitta, K., Akhtar, S., Personett, D., Miller, K.C., Thompson, K.J., Carr, J., Kumar, S., Roy, V., Ansell, S.M., et al. (2014). AT-101 downregulates BCL2 and MCL1 and potentiates the cytotoxic effects of lenalidomide and dexamethasone in preclinical models of multiple myeloma and Waldenstrom macroglobulinaemia. British journal of haematology 164, 352–365.

Perfetto, S.P., Chattopadhyay, P.K., Lamoreaux, L., Nguyen, R., Ambrozak, D., Koup, R.A., and Roederer, M. (2006). Amine reactive dyes: an effective tool to discriminate live and dead cells in polychromatic flow cytometry. J Immunol Methods 313, 199–208.

Prashad, S.L., Drusbosky, L., Sibai, H., Minden, M.D., Western, S.J., Biondi, C., Shah, R., Liu, D., Nguyen, T., Warnock, C., et al. (2016). Ex Vivo High-Throughput Flow Cytometry Screening Identifies Subsets of Responders to Differentiation Agents in Individual AML Patient Samples. Blood 128, 5206–5206.

Prokhorova, E.A., Egorshina, A.Y., Zhivotovsky, B., and Kopeina, G.S. (2020). The DNA-damage response and nuclear events as regulators of nonapoptotic forms of cell death. Oncogene 39, 1–16.

Robinson, N., Ganesan, R., Hegedus, C., Kovacs, K., Kufer, T.A., and Virag, L. (2019). Programmed necrotic cell death of macrophages: Focus on pyroptosis, necroptosis, and parthanatos. Redox biology 26, 101239.

Rodriguez-Hernandez, A., Brea-Calvo, G., Fernandez-Ayala, D.J., Cordero, M., Navas, P., and Sanchez-Alcazar, J.A. (2006). Nuclear caspase-3 and caspase-7 activation, and poly(ADP-ribose) polymerase cleavage are early events in camptothecin-induced apoptosis. Apoptosis: an international journal on programmed cell death 11, 131–139.

Rojo, F., Garcia-Parra, J., Zazo, S., Tusquets, I., Ferrer-Lozano, J., Menendez, S., Eroles, P., Chamizo, C., Servitja, S., Ramirez-Merino, N., et al. (2012). Nuclear PARP-1 protein overexpression is associated with poor overall survival in early breast cancer. Annals of oncology: official journal of the European Society for Medical Oncology 23, 1156–1164.

Saad, J.J., Miettinen, J., Tsallos, D., Eldfors, S., Kontro, M., Wennerberg, K., Kallioniemi, O., Porkka, K., Tang, J., and Heckman, C.A. (2018). Predictive Response Biomarkers for BET Inhibitors in AML. Blood 132, 2749–2749.

Sanger Institute (2021). https://cancer.sanger.ac.uk/cosmic/gene/analysis?ln=PARP1. Cancer Genome Project.

Schiffer, C., and Stone, R. (2003). Morphologic classification and clinical and laboratory correlates. Holland-Frei Cancer Medicine 6th ed Hamilton, Canada: BC Decker.

Schneider, C., Oellerich, T., Baldauf, H.-M., Schwarz, S.-M., Thomas, D., Flick, R., Bohnenberger, H., Kaderali, L., Stegmann, L., Cremer, A., et al. (2017). SAMHD1 is a biomarker for cytarabine response and a therapeutic target in acute myeloid leukemia. Nature Medicine 23, 250–255.

Seipel, K., Marques, M.A.T., Sidler, C., Mueller, B.U., and Pabst, T. (2018). MDM2- and FLT3-inhibitors in the treatment of FLT3-ITD acute myeloid leukemia, specificity and efficacy of NVP-HDM201 and midostaurin. Haematologica 103, 1862–1872.

Skinner, B.M., and Johnson, E.E.P. (2017). Nuclear morphologies: their diversity and functional relevance. Chromosoma 126, 195–212.

Soriano, J., Mora-Espi, I., Alea-Reyes, M.E., Perez-Garcia, L., Barrios, L., Ibanez, E., and Nogues, C. (2017). Cell Death Mechanisms in Tumoral and Non-Tumoral Human Cell Lines Triggered by Photodynamic Treatments: Apoptosis, Necrosis and Parthanatos. Scientific reports 7, 41340.

Staib, P., Tiehen, J., Strunk, T., and Schinkothe, T. (2005). Determination of caspase-3 activation fails to predict chemosensitivity in primary acute myeloid leukemia blasts. BMC Cancer 5, 60.

Stirewalt, D.L., and Radich, J.P. (2003). The role of FLT3 in haematopoietic malignancies. Nature reviews Cancer 3, 650–665.

Tiacci, E., Spanhol-Rosseto, A., Martelli, M.P., Pasqualucci, L., Quentmeier, H., Grossmann, V., Drexler, H.G., and Falini, B. (2012). The NPM1 wild-type OCI-AML2 and the NPM1-mutated OCI-AML3 cell lines carry DNMT3A mutations. Leukemia 26, 554–557.

Triemer, T., Messikommer, A., Glasauer, S.M.K., Alzeer, J., Paulisch, M.H., and Luedtke, N.W. (2018). Superresolution imaging of individual replication forks reveals unexpected prodrug resistance mechanism. Proceedings of the National Academy of Sciences of the United States of America 115, E1366–E1373.

Vincelette, N.D., and Yun, S. (2014). Assessing the Mechanism of Cytarabine-Induced Killing in Acute Leukemia. Blood 124, 5210–5210.

Walter, R.B., Othus, M., Burnett, A.K., Lowenberg, B., Kantarjian, H.M., Ossenkoppele, G.J., Hills, R.K., Ravandi, F., Pabst, T., Evans, A., et al. (2015). Resistance prediction in AML: analysis of 4601 patients from MRC/NCRI, HOVON/SAKK, SWOG and MD Anderson Cancer Center. Leukemia 29, 312–320.

Wang, L., Cai, W., Zhang, W., Chen, X., Dong, W., Tang, D., Zhang, Y., Ji, C., and Zhang, M. (2015). Inhibition of poly(ADP-ribose) polymerase 1 protects against acute myeloid leukemia by suppressing the myeloproliferative leukemia virus oncogene. Oncotarget 6, 27490–27504.

Wang, Y., An, R., Umanah, G.K., Park, H., Nambiar, K., Eacker, S.M., Kim, B., Bao, L., Harraz, M.M., Chang, C., et al. (2016). A nuclease that mediates cell death induced by DNA damage and poly(ADP-ribose) polymerase-1. Science 354, aad6872.

Wang, Y., Dawson, V.L., and Dawson, T.M. (2009). Poly(ADP-ribose) signals to mitochondrial AIF: a key event in parthanatos. Experimental neurology 218, 193–202.

Wang, Y., Kim, N.S., Haince, J.F., Kang, H.C., David, K.K., Andrabi, S.A., Poirier, G.G., Dawson, V.L., and Dawson, T.M. (2011). Poly(ADP-ribose) (PAR) binding to apoptosis-inducing factor is critical for PAR polymerase-1-dependent cell death (parthanatos). Science signaling 4, ra20.

Wong, T.N., Ramsingh, G., Young, A.L., Miller, C.A., Touma, W., Welch, J.S., Lamprecht, T.L., Shen, D., Hundal, J., Fulton, R.S., et al. (2015). Role of TP53 mutations in the origin and evolution of therapy-related acute myeloid leukaemia. Nature 518, 552–555.

Xia, X., Wang, X., Cheng, Z., Qin, W., Lei, L., Jiang, J., and Hu, J. (2019). The role of pyroptosis in cancer: pro-cancer or pro-“host”? Cell Death Dis 10, 650.

Yu, S.W., Andrabi, S.A., Wang, H., Kim, N.S., Poirier, G.G., Dawson, T.M., and Dawson, V.L. (2006). Apoptosis-inducing factor mediates poly(ADP-ribose) (PAR) polymer-induced cell death. Proceedings of the National Academy of Sciences of the United States of America 103, 18314–18319.

Zuber, J., Radtke, I., Pardee, T.S., Zhao, Z., Rappaport, A.R., Luo, W., McCurrach, M.E., Yang, M.M., Dolan, M.E., Kogan, S.C., et al. (2009). Mouse models of human AML accurately predict chemotherapy response. Genes & development 23, 877–889.

