## Supplemental Information for "Cancer Prognosis According to Parthanatos Features"

**Table S1: Summary of parthanatos features in PBMCs from healthy human donors.** Ring-shaped DNA fragmentation, and % of cells rescued from a clinically-relevant mixture of ara-C (5 μM) + idarubicin (0.3 μM) by pre-treatment with Olaparib (1 μM, 24 h).

| Person | Sex | Birth Year | Ring-Shaped Chromatin | Olaparib Rescue / % |
|---|---|---|---|---|
| **A** | m | 1951 | **yes** | **yes / 15%** |
| **B** | - | 1953 | **yes** | **yes / 12%** |
| **C** | f | 1960 | **yes** | **yes / 14%** |
| **D** | m | 1970 | **yes** | **yes / 15%** |
| **E** | m | 1971 | **yes** | **yes / 16%** |
| **F** | m | 1973 | **yes** | **yes / 14%** |
| **G** | f | 1982 | **yes** | **yes / 15%** |
| **H** | m | 1986 | **yes** | **yes / 33%** |
| **I** | f | 1990 | **yes** | **yes / 8%** |
| **J** | - | E | **yes** | **yes / 6%** |

**Table S2: The % of living cells in PBMC samples from healthy human donors.** Cells were pre-treated with Olaparib (+Olap, 1 μM, 24 h) or carrier only (-Olap), followed by incubation with a clinically relevant, 17:1 mixture of ara-C (5 μM) + idarubicin (0.3 μM) for 24 h. Cells were then treated with a live/dead fixable stain, fixed, stained with DAPI and analyzed by flow cytometry.

| | No Ara-C/Idarubicin | | +5 μM Ara-C/0.3 μM Ida | | +5 μM Ara-C/0.3 μM Ida | | |
|---|---|---|---|---|---|---|---|
| **Donor** | -Olap (raw %) | +Olap (raw %) | - Olap (raw %) | **- Olap normalized** | + Olap (raw %) | **+ Olap normalized** | **% rescue** |
| **A** | 85.4 | 87.0 | 71.4 | **82.8** | 84.6 | **98.1** | **15.3** |
| **B** | 91.0 | 89.9 | 74.7 | **82.6** | 85.3 | **94.3** | **11.7** |
| **C** | 87.0 | 88.6 | 73.8 | **84.1** | 86.2 | **98.2** | **14.1** |
| **D** | 88.4 | 89.4 | 70.9 | **79.8** | 84.1 | **94.6** | **14.8** |
| **E** | 90.7 | 89.2 | 65.8 | **73.2** | 79.9 | **88.8** | **15.7** |
| **F** | 80.5 | 82.2 | 67.1 | **82.5** | 77.9 | **95.8** | **13.3** |
| **G** | 89.9 | 89.2 | 74.2 | **82.9** | 87.3 | **97.5** | **14.6** |
| **H** | 90.7 | 90.4 | 39.4 | **43.5** | 69.3 | **76.5** | **33.0** |
| **I** | 86.7 | 89.1 | 79.7 | **90.7** | 86.5 | **98.4** | **7.7** |
| **J** | 70.5 | 72.2 | 63.5 | **89.0** | 68.1 | **95.4** | **6.4** |

**Table S3: Profiles of AML patients in this study.**

| Patient # | ID | Age | Sex | FAB | Risk Group | Molecular Diagnosis | Cytogenetics |
|---|---|---|---|---|---|---|---|
| **1** | **04-032** | 32 | f | M4 | favorable | inv(16) | inv(16) |
| **2** | **04-045** | 26 | f | M5 | intermediate | FLT3-ITD, NPM1mut | normal |
| **3** | **05-002** | 46 | m | M4 | favorable | inv(16) | inv(16) |
| **4** | **15-084** | 65 | m | M5 | intermediate | normal | normal |
| **5** | **15-105** | 68 | f | M5 | favorable | NPM1mut | normal |
| **6** | **16-092** | 62 | m | M4 | adverse | RUNX1mut | inv(7) |
| **7** | **17-008** | 34 | m | M4eo | favorable | inv(16), c-kit | inv(16) |
| **8** | **17-016** | 69 | f | M4 | favorable | NPM1mut | normal |
| **9** | **17-036** | 43 | f | sec. M4 (MDS) | adverse | FLT3-ITD, NPM1mut | normal |
| **10** | **17-063** | 71 | f | M4 | intermediate | IDH2 mut | normal |
| **11*** | **PID 145** | 56 | f | M4 | favorable | NPM1A | 46,XX |
| **12*** | **PID 160** | 70 | m | M4 | intermediate | normal | 46,XY |
| **13*** | **PID 198** | 69 | f | M4 | adverse | NPM1A | 46,XX,t(2;22) |
| **14** | **PID 752** | 28 | m | M4eo | favorable | CBFB-MYH11, Type A, KIT p.Asp816Val | inv(16),(p13q22) |
| **15** | **PID 52** | 43 | f | M5 | adverse | ASXL1 | tet2 |
| **16** | **PID 528** | 64 | f | M5 | intermediate | FLT3-ITD, NPM1 | tet2 |
| **17** | **PID 242** | 45 | m | M5 | adverse | FLT3-ITD, FLT3-TKD | normal |
| **18** | **PID 127** | 63 | f | M5 | adverse | IDH2, DNMT3A | normal |
| **19** | **04-015** | 69 | f | M4 | adverse | t(11,17) | t(11,17) |
| **20** | **15-119** | 74 | f | M5 | favorable | NPM1mut | normal |
| **21** | **15-130** | 68 | m | M5 | favorable | NPM1mut | normal |
| **22** | **16-007** | 79 | m | M4 | intermediate | unknown | unknown |
| **23** | **16-062** | 70 | m | M4 | favorable | inv(16) | inv(16) |
| **24** | **16-068** | 73 | f | M5 | favorable | NPM1mut | normal |
| **25** | **17-003** | 57 | m | M5a | intermediate | NPM1mut, U2AF1, ASXL1 | t(9;11) |
| **26** | **17-024** | 64 | m | M4 | favorable | inv(16), IDH2mut | inv(16), +8 |
| **27*** | **17-039** | 58 | m | M4 | favorable | NPM1mut | normal |
| **28** | **17-045** | 68 | f | M4 | intermediate | FLT3-ITD, NPM1 | normal |
| **29*** | **17-014** | 62 | m | M4 | adverse | RUNX1mut | inv(7) |
| **30*** | **PID 624** | 35 | m | M5 a | adverse | MECOM (EVI1) | 46,XY,t(10;11) |
| **31** | **PID 625** | 48 | m | M4 | favorable | CBFB-MYH11, Type A | inv(16) (p13q22) |
| **32** | **PID 766** | 28 | f | M5 | adverse | FLT3-TKD, EVI1, RUNX1 | wt1 |
| **33** | **PID 154** | 46 | m | M5 | adverse | EVI1 | normal |
| **34** | **PID 218** | 58 | m | M5 | favorable | NPM1, DNMT3A | normal |
| **35** | **PID 469** | 50 | m | M5 | intermediate | FLT3-ITD, NPM1 | normal |
| **36** | **PID 103** | 59 | f | M5 | favorable | FLT3-TKD, NPM1, DNMT3A, PTKN11 | normal |
| **37** | **PID 176** | 59 | m | M5 | adverse | FLT3-ITD/TKD, NPM1, DNMT3A | normal |
| **38** | **PID 171** | 64 | m | M4 | adverse | EVI1, RUNX1, PTKN11 | normal |
| **39** | **PID 517** | 45 | m | M5 | intermediate | FLT3-TKD | normal |

*Bone marrow samples.

**Table S4: Overall survival (OS) and event-free survival (EFS) of AML patients in this study.**

| | Patient # | ID | Therapy | Stem Cell Transplantation | OS (months) | EFS (months) | Alive |
|---|---|---|---|---|---|---|---|
| Exhibiting BOTH Parthanatos Features | 1 | 04-032 | curative | autologous in 1. CR | 6.0 | 6.0 | no |
| | 2 | 04-045 | curative | allogenic in 2. CR | 15 | 7.0 | no |
| | 3 | 05-002 | curative | autologous in 1. CR | 161 | 161 | yes |
| | 4 | 15-084 | curative | autologous in 1. CR | 34 | 34 | yes |
| | 5 | 15-105 | curative | - | 11 | 8.0 | no |
| | 6 | 16-092 | curative | autologous in 1. CR | 5.0 | 2.0 | no |
| | 7 | 17-008 | curative | autologous in 1. CR | 18 | 10 | yes |
| | 8 | 17-016 | curative | autologous in 1. CR | 16 | 16 | yes |
| | 9 | 17-036 | curative | autologous in 1. CR | 12 | 3.0 | no |
| | 10 | 17-063 | curative | - | 13 | 12 | yes |
| | 11* | PID 145 | curative | allogenic | 75 | 75 | yes |
| | 12* | PID 160 | curative | - | 9.0 | 9.0 | no |
| | 13* | PID 198 | curative | - | 62 | 62 | yes |
| | 14 | PID 752 | curative | allogenic | 16 | 7.0 | yes |
| | 15 | PID 52 | curative | allogenic | 71 | 71 | yes |
| | 16 | PID 528 | curative | allogenic | 43 | 43 | yes |
| | 17 | PID 242 | curative | allogenic | 57 | 57 | yes |
| | 18 | PID 127 | curative | allogenic | 37 | 34 | no |
| Exhibiting ONE or ZERO Parthanatos Features | 19 | 04-015 | palliative | - | 6 | 6 | no |
| | 20 | 15-119 | palliative | - | 6 | 6 | no |
| | 21 | 15-130 | curative | - | 0.5 | 0.5 | no |
| | 22 | 16-007 | none | - | 0.1 | 0.1 | no |
| | 23 | 16-062 | curative | - | 1 | 1 | no |
| | 24 | 16-068 | palliative | - | 17 | 17 | no |
| | 25 | 17-003 | curative | allogenic in 1. CR | 18 | 18 | yes |
| | 26 | 17-024 | curative | - | 0.5 | 0.5 | no |
| | 27* | 17-039 | curative | autologous in 1. CR | 15 | 15 | yes |
| | 28 | 17-045 | curative | - | 12 | 2.0 | no |
| | 29* | 17-014 | curative | autologous in 1. CR | 5 | 2.5 | no |
| | 30* | PID 624 | curative | - | 0.2 | 0.2 | no |
| | 31 | PID 625 | curative | allogenic | 9.0 | 9.0 | no |
| | 32 | PID 766 | curative | allogenic | 38 | 38 | yes |
| | 33 | PID 154 | curative | allogenic | 9.6 | 5.6 | no |
| | 34 | PID 218 | curative | allogenic | 6.3 | 5.6 | no |
| | 35 | PID 469 | curative | allogenic | 34 | 34 | yes |
| | 36 | PID 103 | curative | - | 62 | 62 | yes |
| | 37 | PID 176 | curative | - | 7.0 | 6.4 | no |
| | 38 | PID 171 | curative | allogenic | 6.0 | 5.2 | no |
| | 39 | PID 517 | curative | allogenic | 41 | 41 | yes |

*Bone marrow samples.

**Table S5: The % of living AML cancer cells (CD45+, SS+) in primary isolates treated with a variable concentration of ara-C (1 – 30 μM) in a fixed 17:1 ratio with idarubicin (0.06 – 1.8 μM) following pre-treatment with Olaparib (+Olap, 1 μM, 24 h) or carrier only (-Olap)**. Cells were then treated with a live/dead fixable stain, fixed, stained with a CD45 antibody and DAPI. The blasts were distinguished from other cells by flow cytometry using side light scattering (SS+) and CD45 immunostaining (CD45+). Estimated $EC_{50}$ values are given in terms of ara-C concentrations. The % values are normalized to the number of living cells in samples receiving no ara-C or idarubicin.

| | # | Live AML Cells (%) -Olap | | | | Est. $EC_{50}$ (μM) | Live AML Cells (%) +Olap | | | | Est. $EC_{50}$ (μM) | Ring DNA | Olaparib Rescue / % |
|---|---|---|---|---|---|---|---|---|---|---|---|---|---|
| | | 1μM | 5μM | 15μM | 30μM | | 1μM | 5μM | 15μM | 30μM | | | |
| Exhibiting BOTH Parthanatos Features | 1 | | 96.8 | 62.4 | 15.3 | 17.6 | | 95.6 | 76.0 | 68.1 | 59.0 | yes | yes / 14% |
| | 2 | 86.2 | | 19.9 | 20.0 | 5.2 | 86.5 | | 60.6 | 53.8 | 38.5 | yes | yes / 41% |
| | 3 | | 82.1 | 11.9 | 2.2 | 8.1 | | 82.6 | 51.7 | 9.0 | 13.7 | yes | yes / 40% |
| | 4 | | 78.8 | 62.1 | 10.7 | 15.5 | | 85.9 | 73.9 | 16.5 | 19.6 | yes | yes / 7% |
| | 5 | | 90.9 | 80.8 | 51.1 | 31.7 | | 96.9 | 86.9 | 59.8 | 36.4 | yes | yes / 6% |
| | 6 | | 95.9 | 93.9 | 35.2 | 26.4 | | 101.1 | 100.7 | 43.4 | 29.7 | yes | yes / 7% |
| | 7 | | 65.2 | 42.9 | 9.9 | 9.1 | | 84.6 | 69.1 | 17.1 | 18.8 | yes | yes / 26% |
| | 8 | 86.2 | 70.4 | 56.2 | | 22.9 | 92.7 | 74.3 | 65.3 | | 38.2 | yes | yes / 7% |
| | 9 | 64.5 | 32.2 | 0.1 | | 1.9 | 86.7 | 72.8 | 15.7 | | 7.5 | yes | yes / 41% |
| | 10 | 83.6 | 63.4 | 33.0 | | 7.7 | 87.1 | 69.1 | 25.2 | | 7.7 | yes | yes / 6% |
| | 11* | | 75.5 | 28.7 | 0.0 | 8.9 | | 89.6 | 23.9 | 0.0 | 10.2 | yes | yes / 14% |
| | 12* | | 91.3 | 72.8 | 0.0 | 15.9 | | 101.0 | 54.1 | 0.0 | 15.1 | yes | yes / 10% |
| | 13* | | 71.2 | 37.3 | 0.0 | 9.1 | | 89.1 | 24.1 | 0.0 | 10.2 | yes | yes / 18% |
| | 14 | 74.7 | | 23.1 | 1.1 | 3.1 | 95.5 | | 34.3 | 1.5 | 13.2 | yes | yes / 21% |
| | 15 | | 90.7 | 74.9 | 1.1 | 17.2 | | 98.7 | 80.5 | 2.1 | 18.1 | yes | yes / 8% |
| | 16 | | 65.6 | 5.7 | 0.5 | 6.1 | | 73.9 | 6.0 | 0.3 | 6.8 | yes | yes / 8% |
| | 17 | | 91.0 | 71.4 | 3.7 | 17.5 | | 101.7 | 92.2 | 9.8 | 21.6 | yes | yes / 11% |
| | 18 | | 89.8 | 71.7 | 0.8 | 16.8 | | 101.2 | 73.2 | 1.5 | 17.1 | yes | yes / 11% |
| | **AVE ± ST** | **79 ± 9 %** | **78 ± 17 %** | **47 ± 28 %** | **10 ± 15 %** | **13 ± 8 μM** | **90 ± 4 %** | **89 ± 11 %** | **56 ± 28 %** | **19 ± 25 %** | **21 ± 14 μM** | | |
| Exhibiting ONE or ZERO Parthanatos Features | 19 | | 6.7 | 0.1 | 0.0 | 2.5 | | 10.6 | 0.1 | 0.0 | 3.0 | no | no / 4% |
| | 20 | | 91.5 | 28.5 | 0.0 | 11.1 | | 92.3 | 28.3 | 0.0 | 11.2 | no | no / 1% |
| | 21 | | 26.2 | 0.1 | 0.0 | 4.2 | | 27.4 | 0.0 | 0.0 | 4.3 | no | no / 1% |
| | 22 | | 92.2 | 4.2 | 0.0 | 8.1 | | 79.6 | 0.1 | 0.0 | 6.0 | no | no / 0% |
| | 23 | | 64.8 | 88.8 | 0.0 | 17.3 | | 64.2 | 4.6 | 0.0 | 6.0 | no | no / 1% |
| | 24 | | 91.3 | 59.7 | 0.0 | 15.6 | | 94.5 | 61.0 | 0.0 | 16.1 | no | no / 3% |
| | 25 | 94.9 | 72.7 | 51.0 | | 15.2 | 100.5 | 70.6 | 37.4 | | 10.1 | no | no / 2% |
| | 26 | | 82.5 | 14.7 | 0.7 | 8.3 | | 84.7 | 18.5 | 1.6 | 9.0 | no | no / 2% |
| | 27* | 77.6 | 0.3 | 0.0 | | 1.3 | 77.9 | 0.3 | 0.0 | | 1.3 | no | no / 0% |
| | 28 | 91.7 | 70.7 | 4.4 | | 6.4 | 91.6 | 62.8 | 5.0 | | 5.9 | no | no / 0% |
| | 29* | 85.3 | 15.9 | 0.1 | | 2.3 | 90.9 | 17.3 | 0.1 | | 2.6 | no | no / 1% |
| | 30* | 87.2 | 45.4 | 7.7 | | 4.1 | 88.5 | 41.1 | 7.9 | | 3.9 | yes | no / 1% |
| | 31 | | 77.4 | 28.1 | 1.4 | 9.1 | | 82.2 | 40.3 | 4.0 | 11.4 | no | yes / 5% |
| | 32 | | 92.0 | 86.7 | 21.0 | 22.5 | | 96.7 | 96.2 | 26.5 | 25.4 | no | yes / 10% |
| | 33 | | 92.8 | 52.9 | 5.9 | 15.3 | | 93.5 | 56.6 | 11.9 | 16.1 | no | no / 1% |
| | 34 | 99.2 | 96.3 | | 0.9 | 10.4 | 100.4 | 99.9 | | 6.4 | 18.7 | no | no / 4% |
| | 35 | | 89.7 | 76.8 | 2.1 | 17.7 | | 104.1 | 82.9 | 8.7 | 19.8 | no | yes / 14% |
| | 36 | 98.2 | 97.1 | | 8.4 | 14.5 | 101.1 | 99.6 | | 10.6 | 18.1 | yes | no / 2% |
| | 37 | 100.5 | 98.7 | | 0.9 | 11.9 | 103.3 | 97.3 | | 1.9 | 11.8 | no | no / 3% |
| | 38 | 99.9 | | 54.3 | 4.8 | 15.6 | 101.6 | | 59.0 | 4.5 | 16.2 | no | no / 2% |
| | 39 | | 91.7 | 29.8 | 0.1 | 11.3 | | 90.3 | 33.7 | 0.4 | 11.6 | no | no / 1% |
| | **AVE ± ST** | **93 ± 8 %** | **70 ± 33 %** | **33 ± 31 %** | **3 ± 5 %** | **11 ± 6 μM** | **95 ± 8 %** | **70 ± 33 %** | **31 ± 33 %** | **5 ± 7 %** | **11 ± 7 μM** | | |

*Bone marrow samples.

**Table S6. Summary of AML Patient Data.** Samples from 39 AML patients were tested for the presence of chromatin fragmentation "ring" morphologies and PARP-dependent changes in drug sensitivity upon addition of Olaparib. Only five of the 39 samples exhibited only one feature in the absence of the other. Assignment of these five patients as being parthanatos positive (Assignment 1), or parthanatos negative (Assignment 2), or being excluded from the analysis (Assignment 3) had little-to-no impact on the conclusions drawn in this study.

| Sample type | Total | Exhibited both parthanatos features | Exhibited one feature | Exhibited neither feature |
|---|---|---|---|---|
| **Blood** | 33 | 15 | 4 | 14 |
| **Bone marrow** | 6 | 3 | 1 | 2 |
| **Total** | **39** | **18** | **5** | **16** |

| **Assignment 1** | | **Positive** | **Negative** |
|---|---|---|---|
| **Assigned patients (N)** | | **18** | **21** |
| **Average age (years)** | | 53 +/- 16 | 59 +/- 13 |
| **Sex (% female)** | | 61 | 29 |
| **FAB (% M4)** | | 50 | 46 |
| **ELN Risk group** | **Adverse (%)** | 33 | 33 |
| | **Intermediate (%)** | 28 | 24 |
| | **Favorable (%)** | 39 | 43 |
| **Two-year overall survival (%)** | | **66 %** | **28 %** |
| **Log-rank (Mantel-Cox) test: hazard ratio (HR) and p value** | | **HR = 0.34, p = 0.02** | |

| **Assignment 2** | | **Positive** | **Negative** |
|---|---|---|---|
| **Assigned patients (N)** | | **23** | **16** |
| **Average Age (years)** | | 51 +/- 15 | 63 +/- 9 |
| **Sex (% female)** | | 61 | 25 |
| **FAB (% M4)** | | 52 | 50 |
| **ELN Risk group** | **Adverse (%)** | 35 | 31 |
| | **Intermediate (%)** | 26 | 25 |
| | **Favorable (%)** | 39 | 44 |
| **Two-year overall survival (%)** | | **67 %** | **17 %** |
| **Log-rank (Mantel-Cox) test: hazard ratio (HR) and p value** | | **HR = 0.22, p = 0.002** | |

| **Assignment 3** | | **Positive** | **Negative** |
|---|---|---|---|
| **Assigned patients (N)** | | **18** | **16** |
| **Average age (years)** | | 53 ± 16 | 63 +/- 9 |
| **Sex (% female)** | | 61 | 25 |
| **FAB (% M4)** | | 50 | 50 |
| **ELN Risk group** | **Adverse (%)** | 33 | 31 |
| | **Intermediate (%)** | 28 | 25 |
| | **Favorable (%)** | 39 | 44 |
| **Two-year overall survival (%)** | | **66 %** | **17 %** |
| **Log-rank (Mantel-Cox) test: hazard ratio (HR) and p value** | | **HR = 0.22, p = 0.002** | |

### Supplementary Figure A: PBMCs from healthy donors exhibiting both parthanatos features

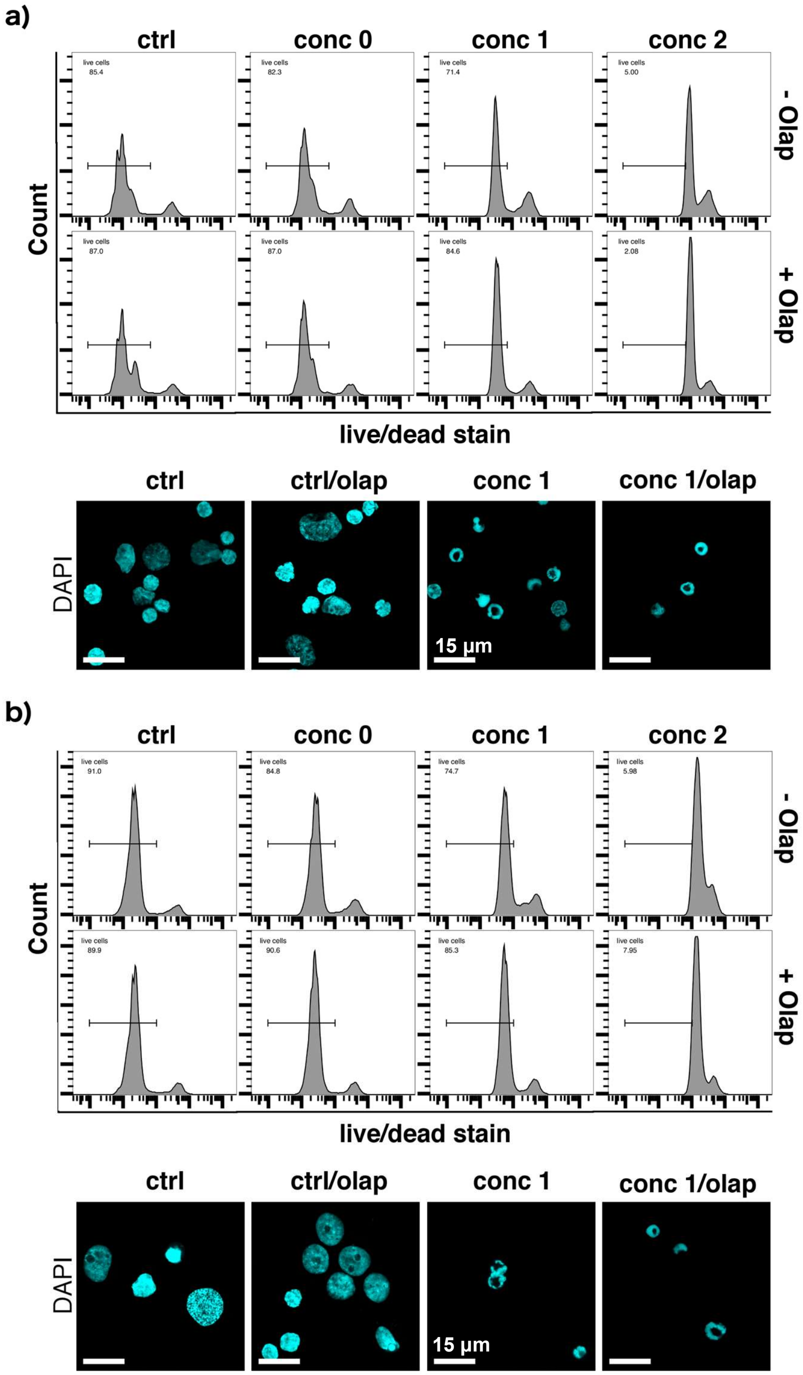


**Fig. A1** Parthanatos features in PBMCs from healthy donors according to toxicity rescue by Olaparib (Olap) and the presence of ring-shaped nuclei examined by DAPI staining. Primary cells: **a)** Human donor 1 (1951); **b)** Human donor 2 (1953). Pretreatment: 1 µM Olaparib o/n; drug treatment: 24 h. Conc 0: 1 µM ara-C + 0.06 µM ida, conc 1: 5 µM ara-C + 0.3 µM ida, conc 2: 15 µM ara-C + 0.9 µM ida.

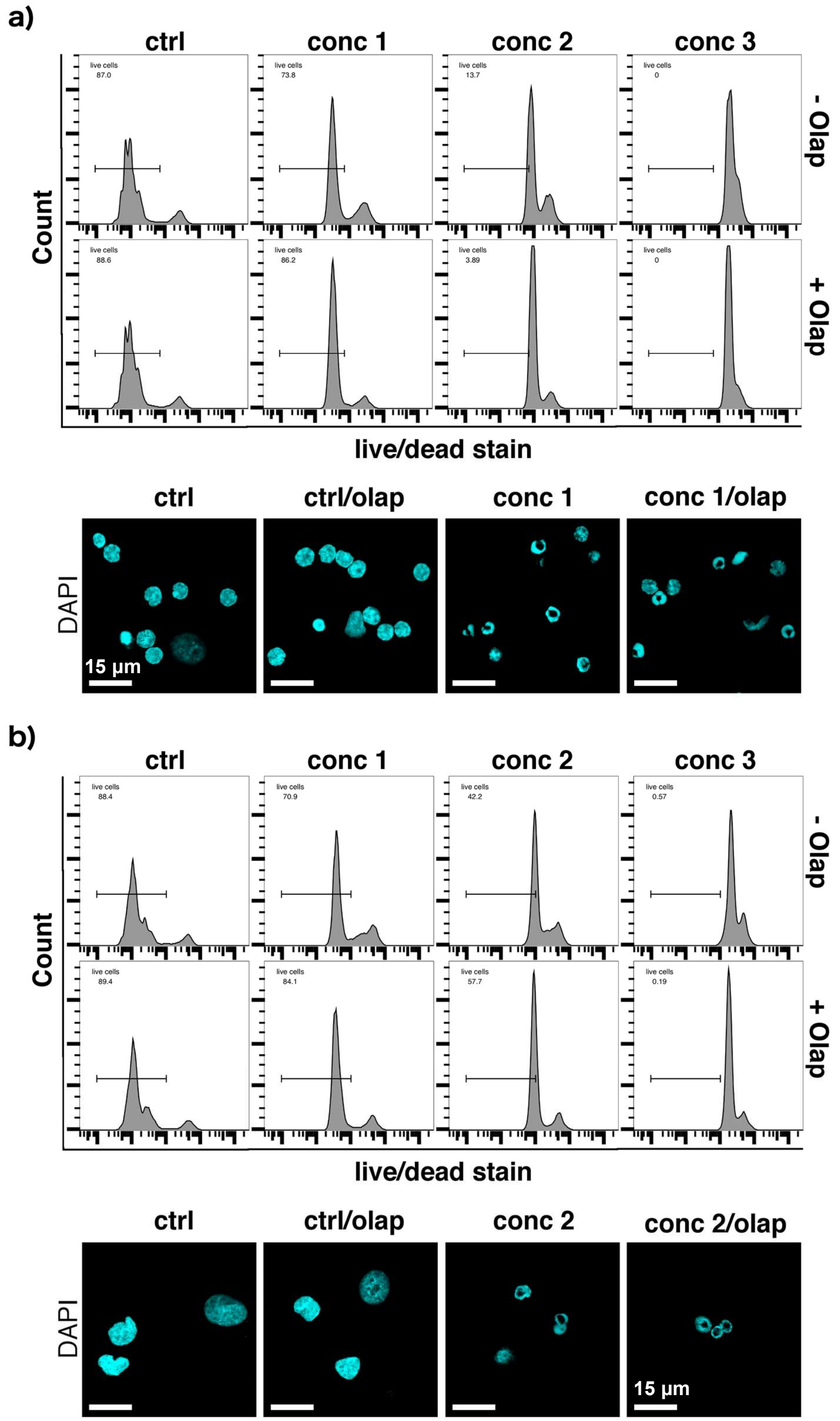


**Fig. A2** Parthanatos features in PBMCs from healthy donors according to toxicity rescue by Olaparib (Olap) and the presence of ring-shaped nuclei examined by DAPI staining. Primary cells: **a)** 3 / 1960 and **b)** 4 / 1970. Pretreatment: 1 µM Olaparib o/n; drug treatment: 24 h. Conc 1: 5 µM ara-C + 0.3 µM ida, conc 2: 15 µM ara-C+ 0.9 µM ida, conc 3: 30 µM ara-C + 1.8 µM ida.

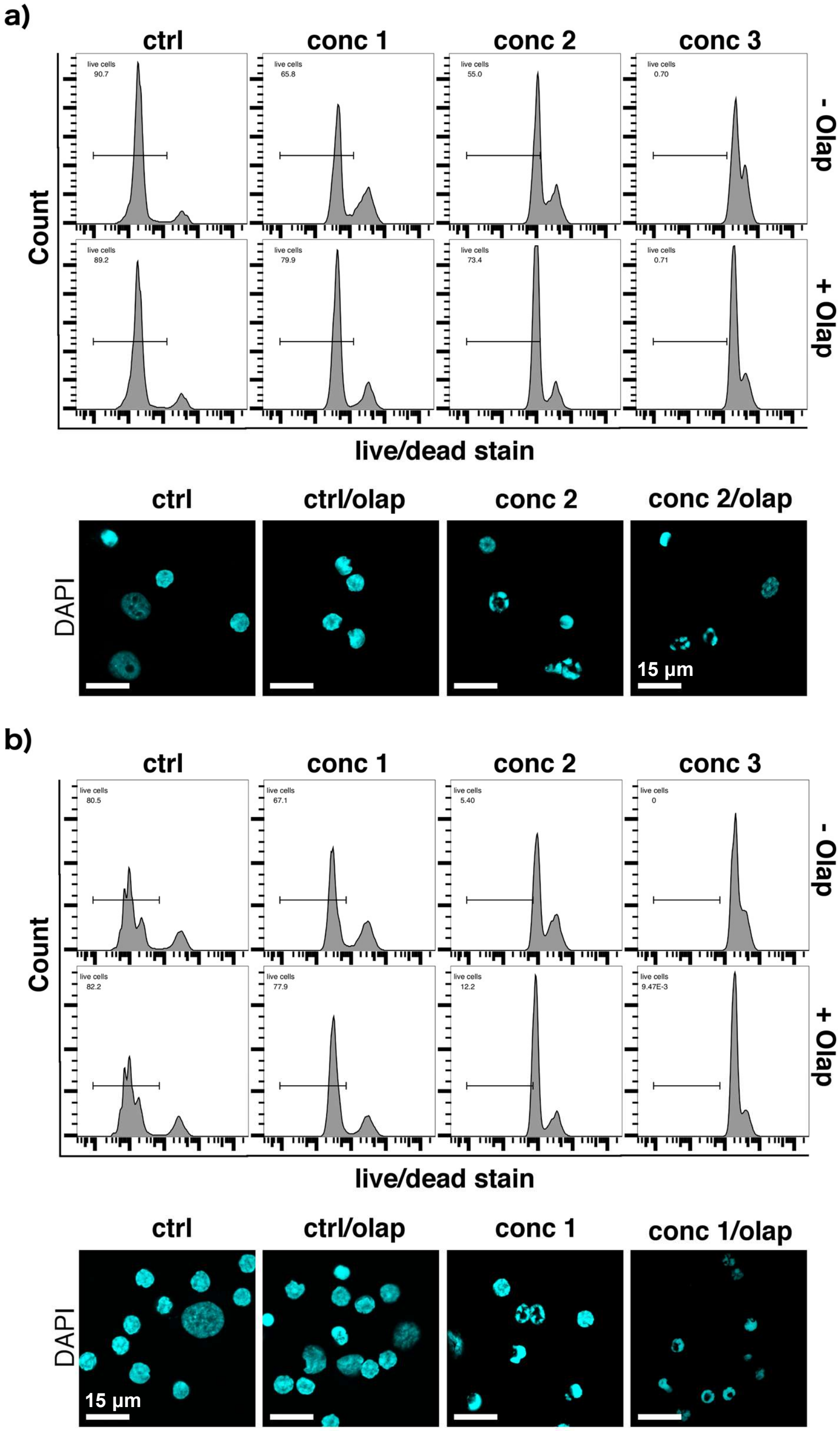


**Fig. A3** Parthanatos features in PBMCs from healthy donors according to toxicity rescue by Olaparib (Olap) and the presence of ring-shaped nuclei examined by DAPI staining. Primary cells: **a)** 5 / 1971 and **b)** 6 / 1973. Pretreatment: 1 µM Olaparib o/n; drug treatment: 24 h. Conc 1: 5 µM ara-C + 0.3 µM ida, conc 2: 15 µM ara-C + 0.9 µM ida, conc 3: 30 µM ara-C + 1.8 µM ida.

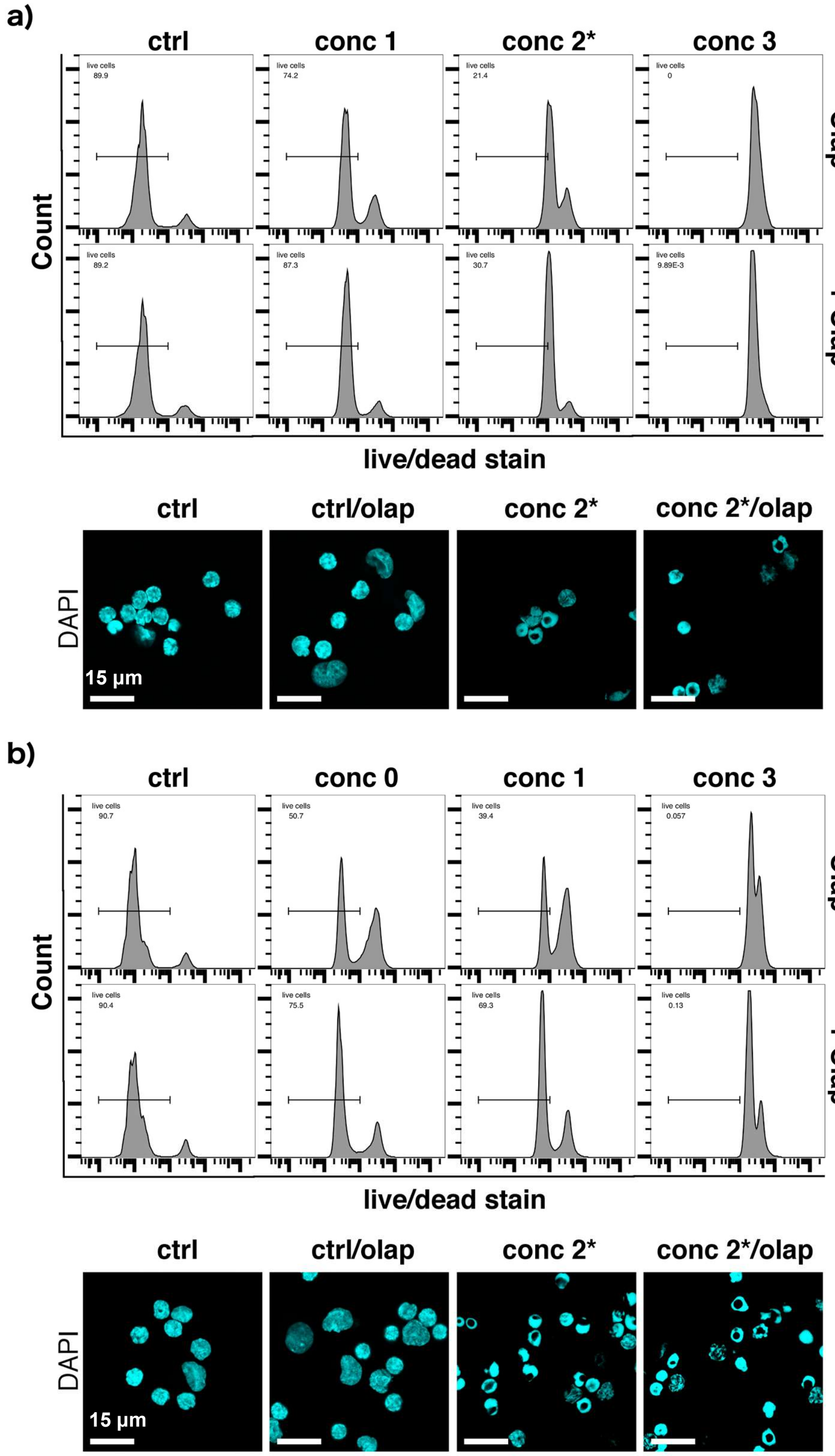


**Fig. A4** Parthanatos features in PBMCs from healthy donors according to toxicity rescue by Olaparib (Olap) and the presence of ring-shaped nuclei examined by DAPI staining. Primary cells: **a)** 7 / 1982 and **b)** 8 / 1986. Pretreatment: 1 µM Olaparib o/n; drug treatment: 24 h. Conc 0: 1 µM ara-C + 0.06 µM ida, conc 1: 5 µM ara-C + 0.3 µM ida, conc 2*: 10 µM ara-C + 0.6 µM ida, conc 3: 30 µM ara-C + 1.8 µM ida.

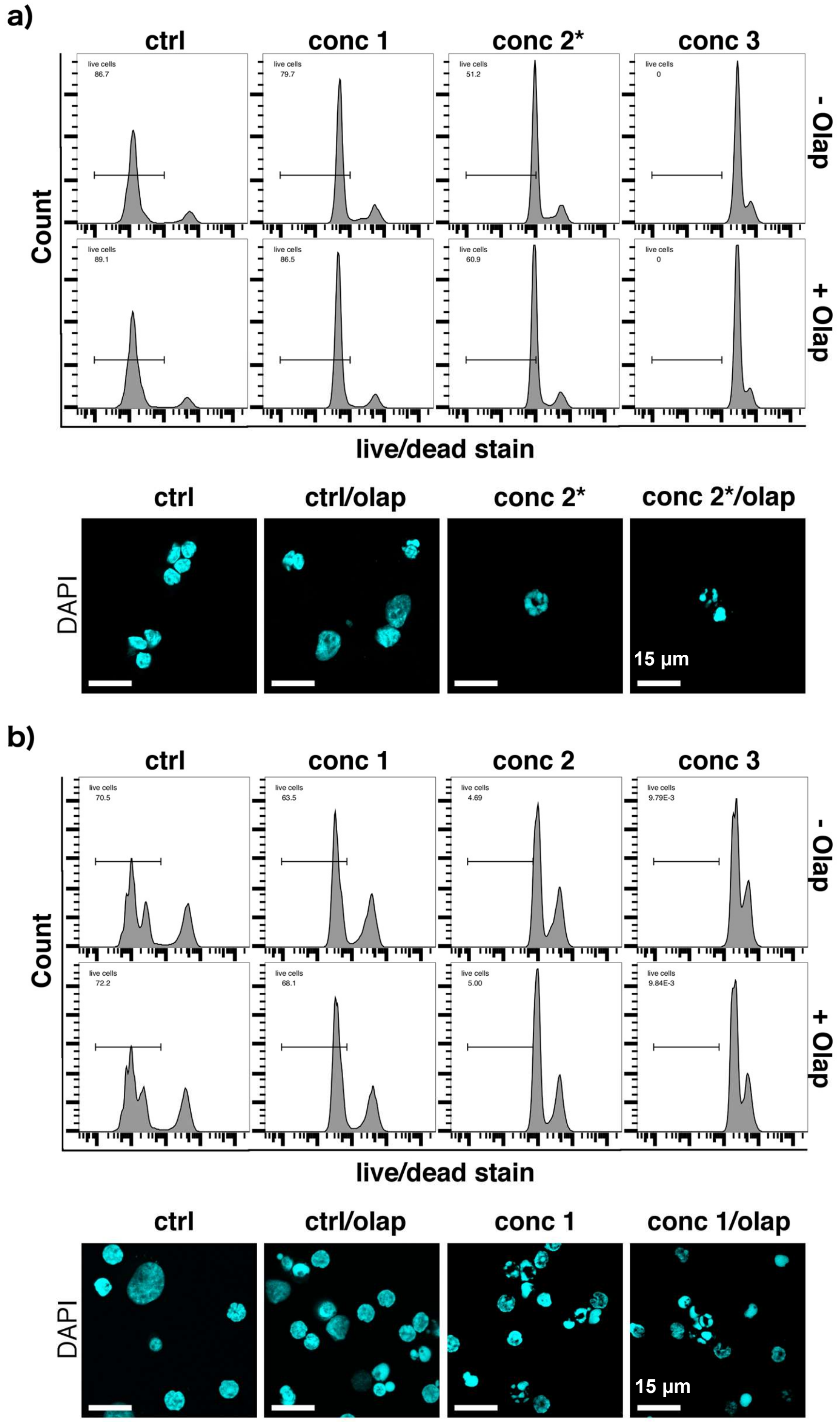


**Fig. A5** Parthanatos features in PBMCs from healthy donors according to toxicity rescue by Olaparib (Olap) and the presence of ring-shaped nuclei examined by DAPI staining. Primary cells: **a)** 9 / 1990 and **b)** 10 / E. Pretreatment: 1 µM Olaparib o/n; drug treatment: 24 h. Conc 1: 5 µM ara-C + 0.3 µM ida, conc 2*: 10 µM ara-C + 0.6 µM ida, conc 2: 15 µM ara-C + 0.9 µM ida, conc 3: 30 µM ara-C + 1.8 µM ida.

**Supplementary Figure B: PBMCs from healthy donors exhibit drug-specific morphologies consistent with parthantos or apoptosis**

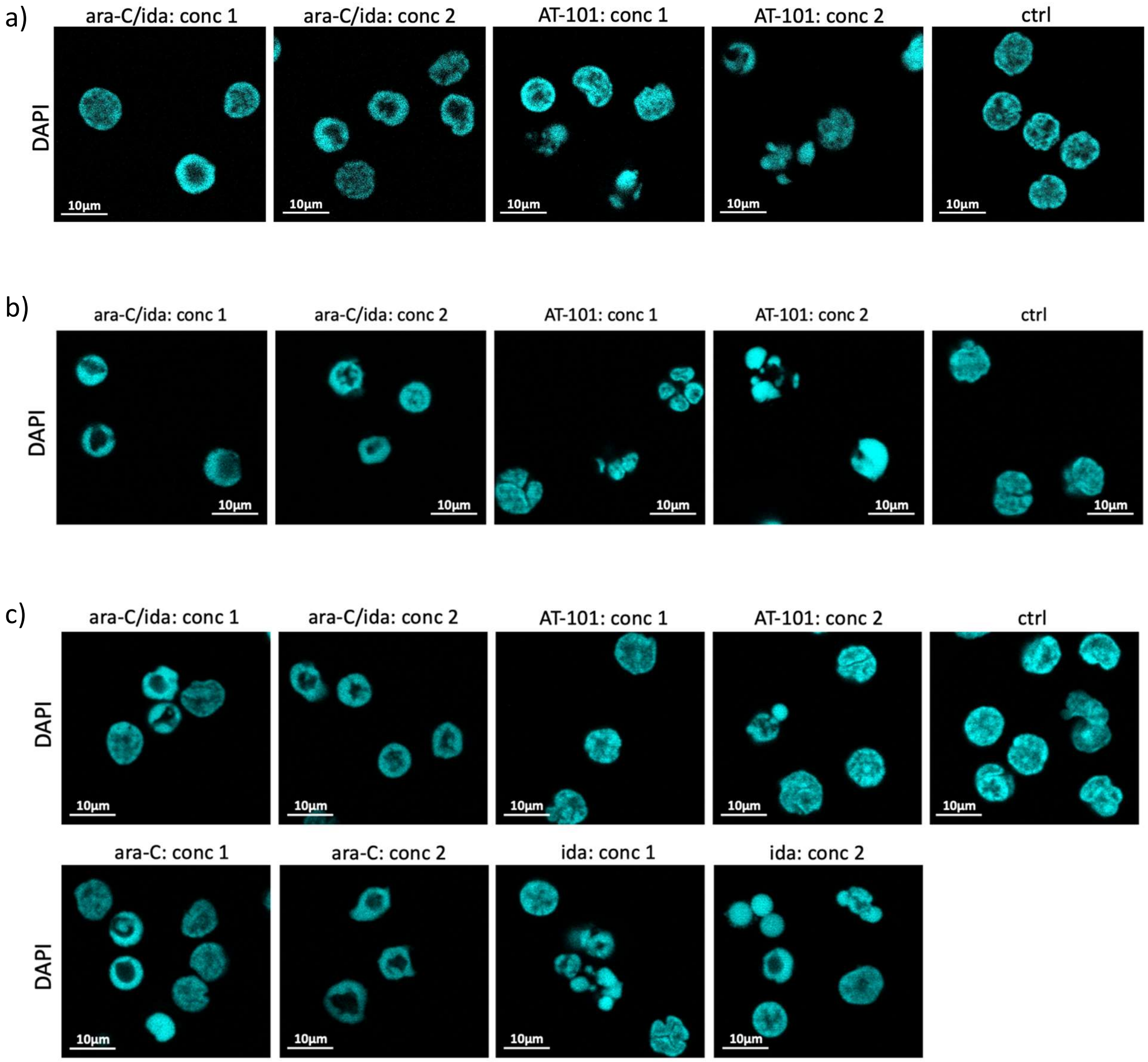


**Fig. B1.** PBMCs from healthy donors upon ex vivo treatment with cytarabine and/or idarubicin, or AT-101. The presence of ring-shaped fragmented nuclei was examined by DAPI staining **a)** HB m 1951, **b)** UL f 1948 **c)** SM f 1954. Drug treatment: 24 h. **ara-C/ida conc 1**: 5 µM ara-C + 0.3 µM ida; **ara-C/ida conc 2**: 15 µM ara-C + 0.9 µM ida; **ara-C conc 1**: 5 µM ara-C; **ara-C conc 2:** 15 µM ara-C; **ida conc 1:** 0.3 µM ida; **ida conc 2:** 0.9 µM ida; **AT-101 conc 1**: 10 µM AT-101; **AT-101 conc 2**: 30 µM AT-101.

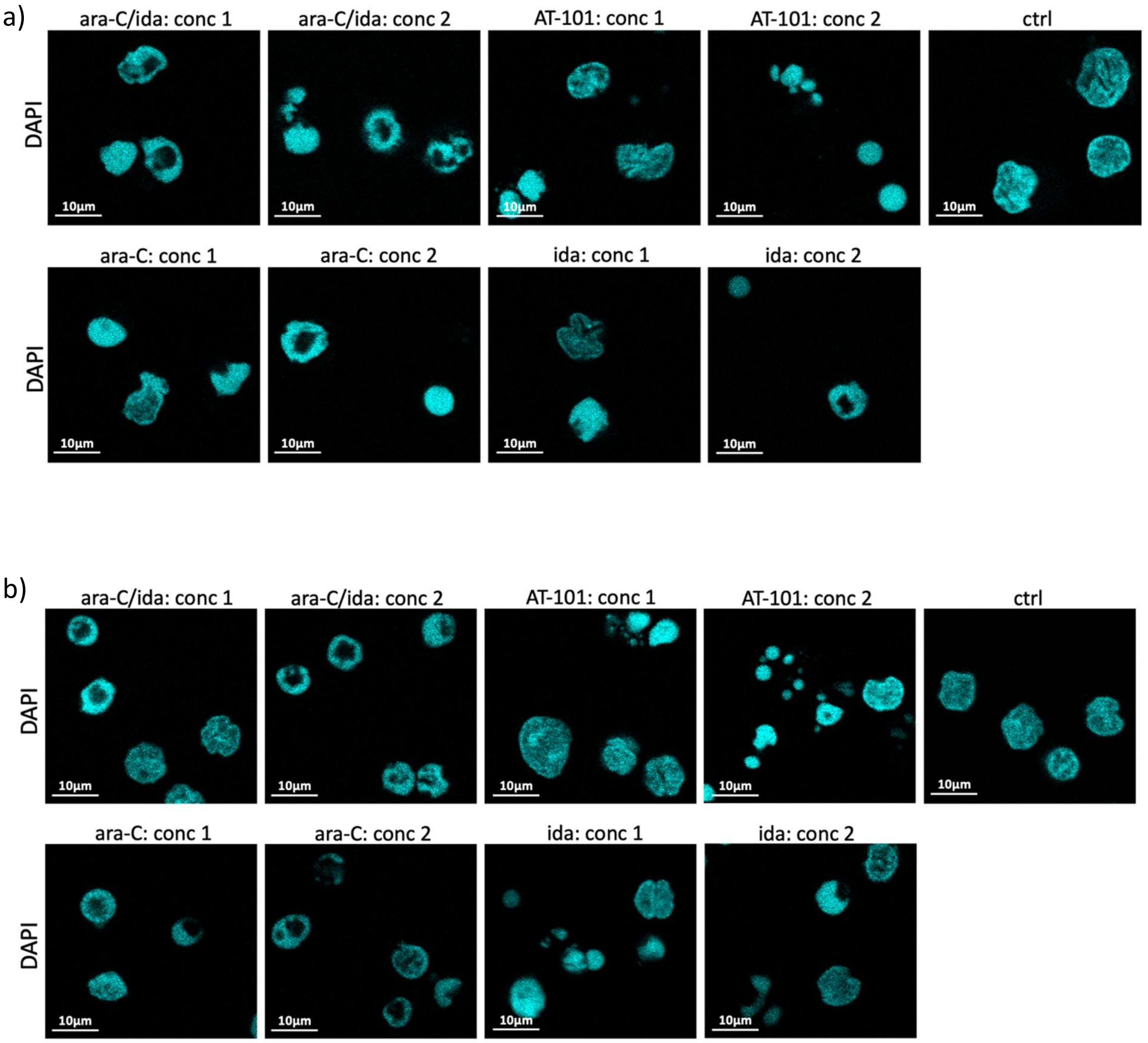


**Fig. B2.** PBMCs from healthy donors upon ex vivo treatment with cytarabine and/or idarubicin, or AT-101. The presence of ring-shaped fragmented nuclei was examined by DAPI staining **a)** HB m 1951, **b)** UL f 1948 **c)** SM f 1954. Drug treatment: 24 h. **ara-C/ida conc 1**: 5 µM ara-C + 0.3 µM ida; **ara-C/ida conc 2**: 15 µM ara-C + 0.9 µM ida; **ara-C conc 1**: 5 µM ara-C; **ara-C conc 2:** 15 µM ara-C; **ida conc 1:** 0.3 µM ida; **ida conc 2:** 0.9 µM ida; **AT-101 conc 1**: 10 µM AT-101; **AT-101 conc 2**: 30 µM AT-101.

**Supplementary Figure C: Evidence for parthanatos features following in vivo drug treatment of AML patients with 7 + 3 induction chemotherapy**

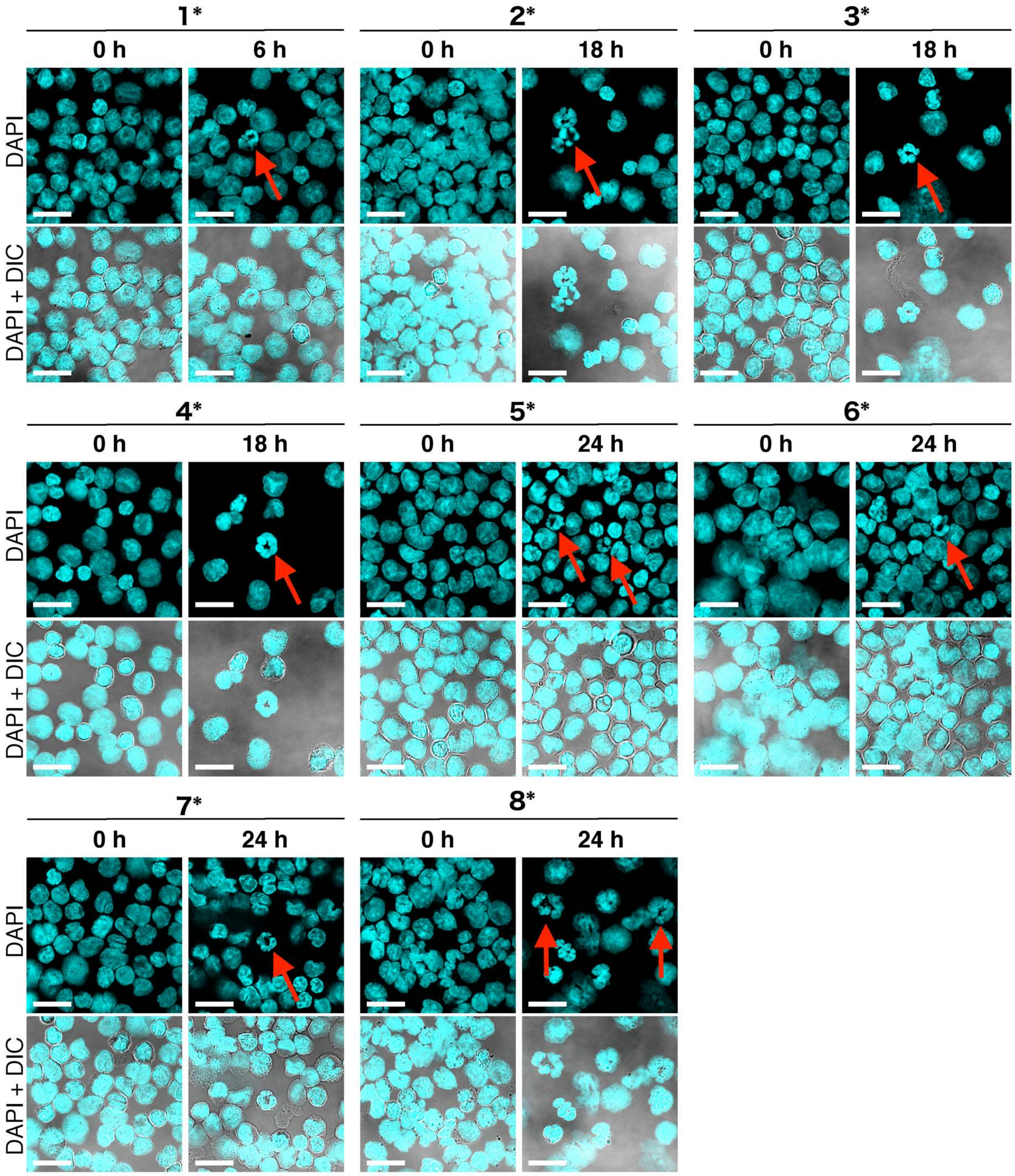


**Fig. C. DAPI staining of monocytes from AML patients collected immediately before or 6 – 24 hours after the onset of “7 + 3” induction chemotherapy with cytarabine (200 mg/m$^2$/d) and idarubicin (12 mg/m$^2$/d). 1*** (patient ID# 17-008, 93% blasts, FAB M4eo, 6 h); **2*** (patient ID# 17-024, 52% blasts, FAB M4eo, 18 h); **3*** (patient ID# 17-038, 38% blasts, FAB M4, 24 h); **4*** (patient ID# 17-040, 15% blasts, FAB M4, 24 h); **5*** (patient ID# 17-064, 1% blasts, FAB M5, 24 h); **6*** (patient ID# 17-068, 26% blasts, FAB M4, 24 h); **7*** (patient ID# 17-110, 47% blasts, MDS-AML, 24 h); **8*** (patient ID# 17-115, FAB M5, 24 h). Scale bar: 15 µm.

### Supplementary Figure D: Primary samples from AML patients exhibiting both parthanatos features

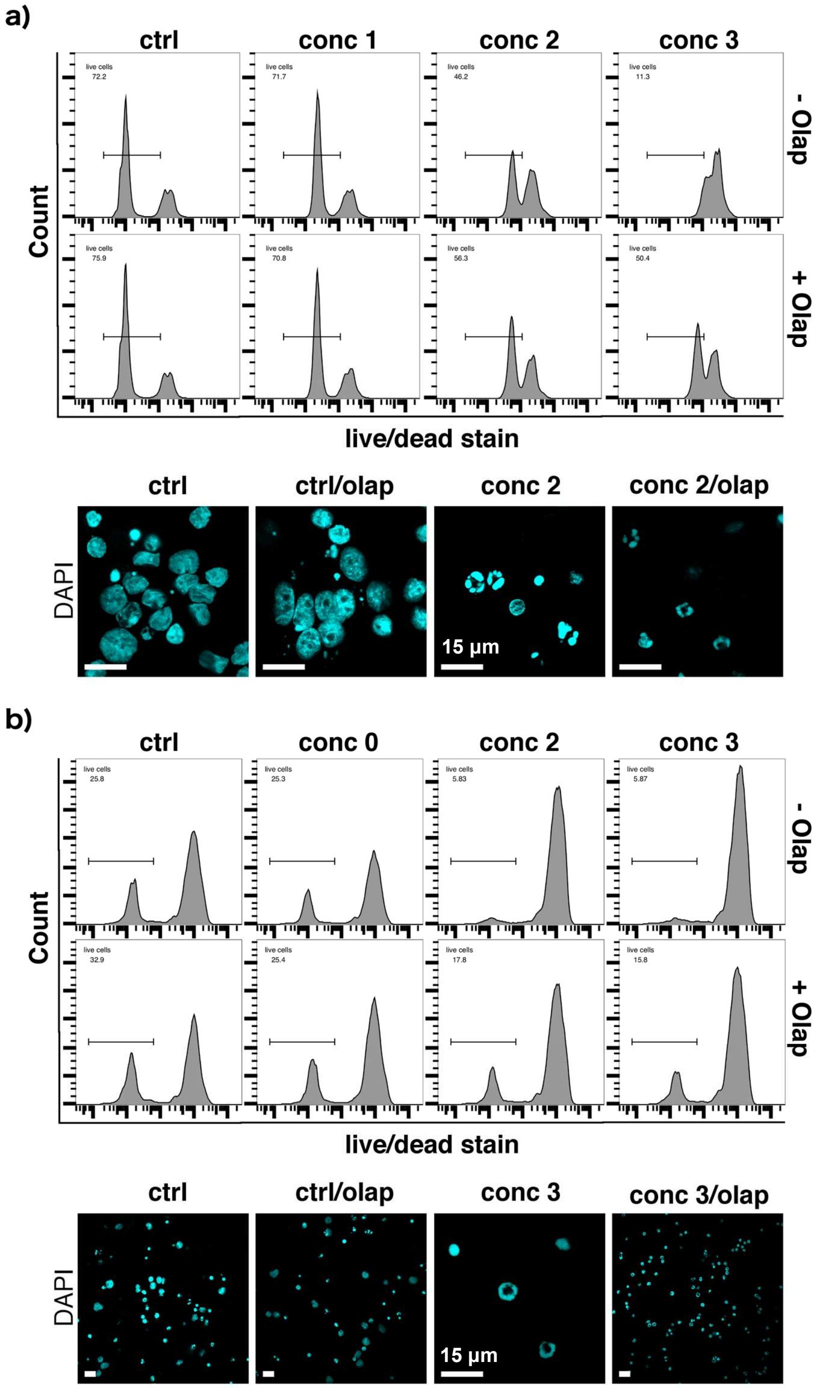


**Fig. D1** Parthanatos features in primary cells from AML donors according to toxicity rescue by Olaparib (Olap) and the presence of ring-shaped nuclei examined by DAPI staining. **a)** 1 / 04-032 and **b)** 2 / 04-045. Pretreatment: 1 µM Olaparib o/n; drug treatment: 24 h. Conc 0: 1 µM ara-C + 0.06 µM ida, conc 1: 5 µM ara-C + 0.3 µM ida, conc 2: 15 µM ara-C + 0.9 µM ida, conc 3: 30 µM ara-C + 1.8 µM ida.

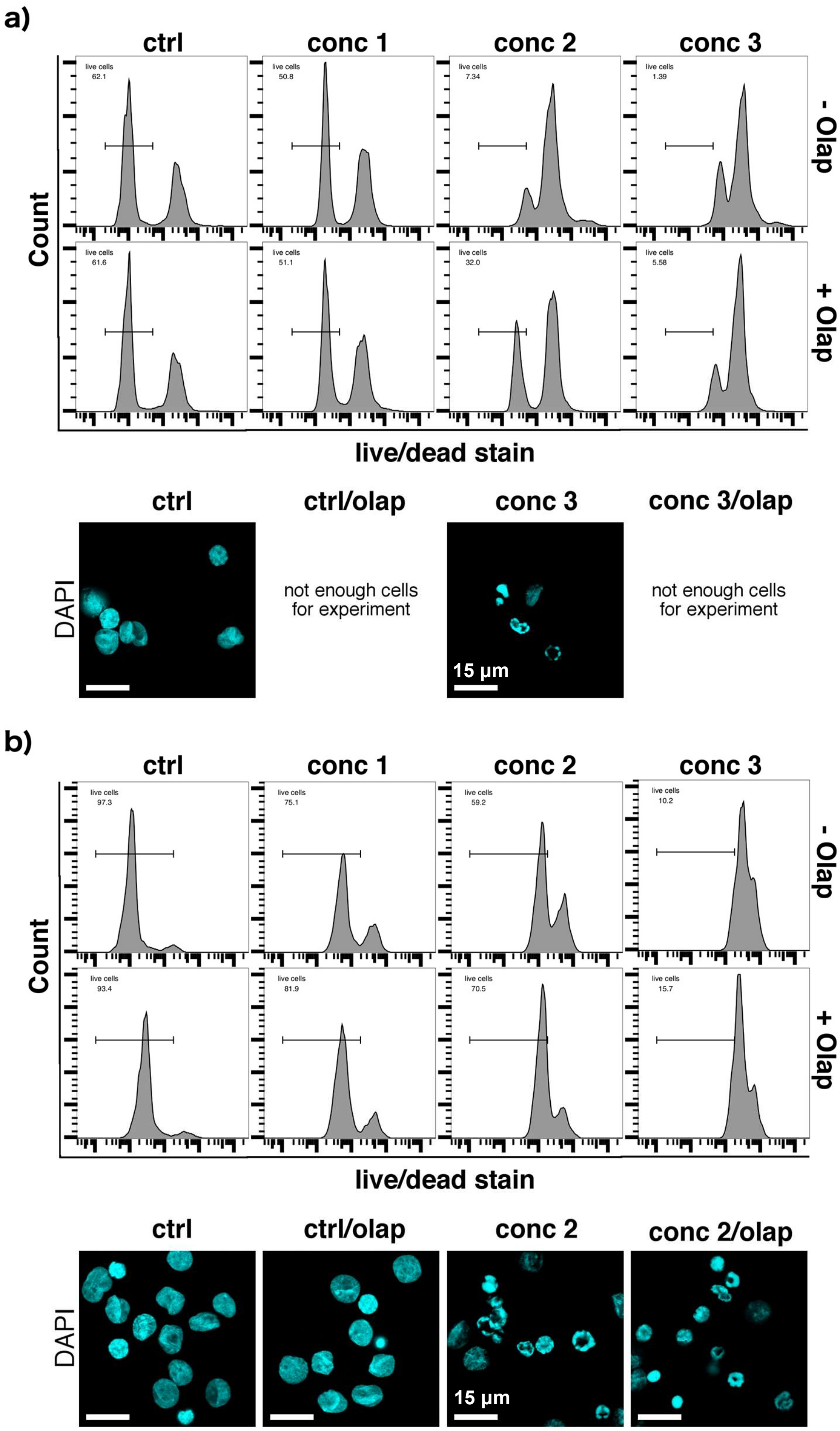


**Fig. D2** Parthanatos features in primary cells from AML donors according to toxicity rescue by Olaparib (Olap) and the presence of ring-shaped nuclei examined by DAPI staining. **a)** 3 / 05-002 and **b)** 4 / 15-084. Pretreatment: 1 µM Olaparib o/n; drug treatment: 24 h. Conc 1: 5 µM ara-C + 0.3 µM ida, conc 2: 15 µM ara-C + 0.9 µM ida, conc 3: 30 µM ara-C + 1.8 µM ida.

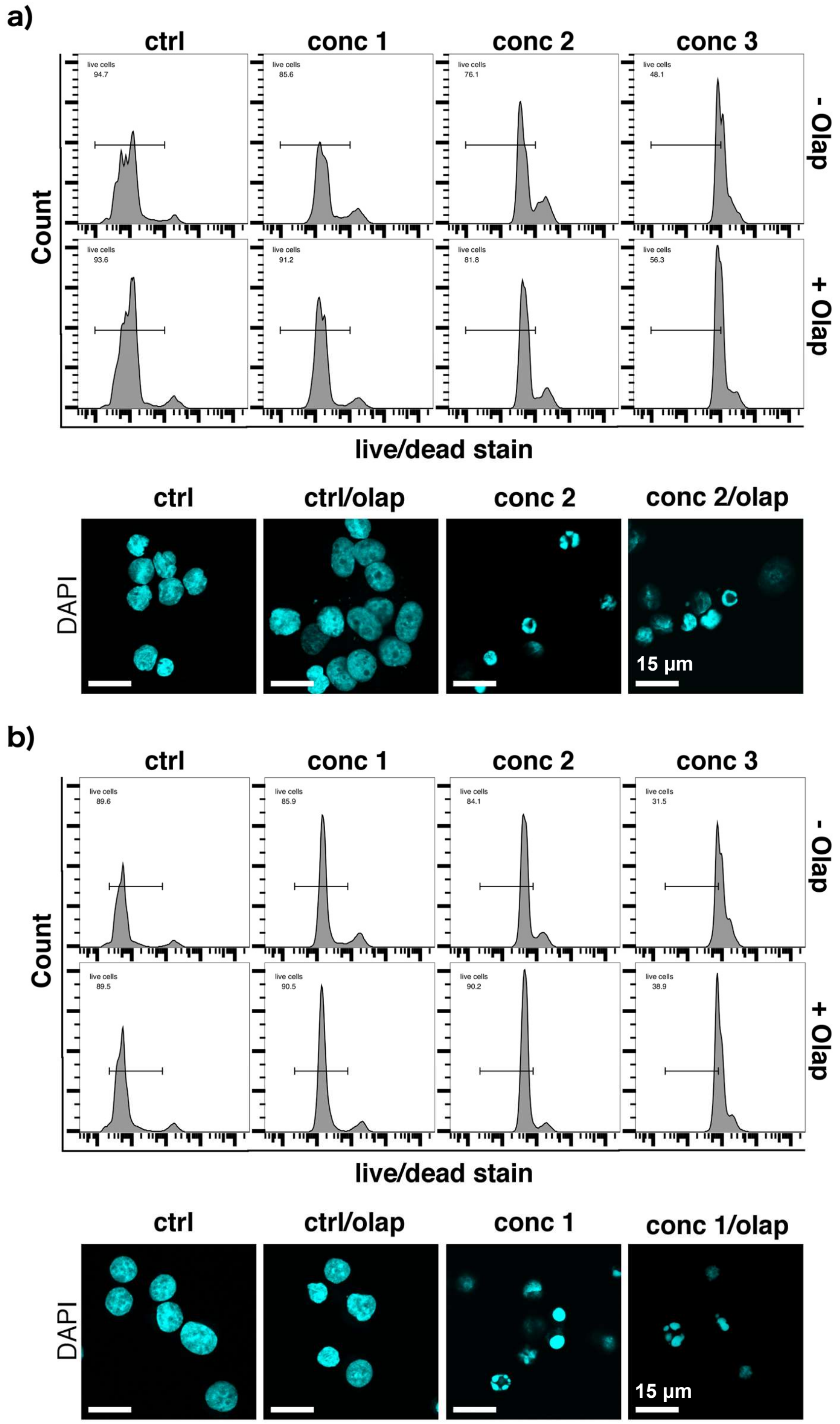


**Fig. D3** Parthanatos features in primary cells from AML donors according to toxicity rescue by Olaparib (Olap) and the presence of ring-shaped nuclei examined by DAPI staining. **a)** 5 /15-105 and **b)** 6 / 16-092. Pretreatment: 1 µM Olaparib o/n; drug treatment: 24 h. Conc 1: 5 µM ara-C + 0.3 µM ida, conc 2: 15 µM ara-C + 0.9 µM ida, conc 3: 30 µM ara-C + 1.8 µM ida.

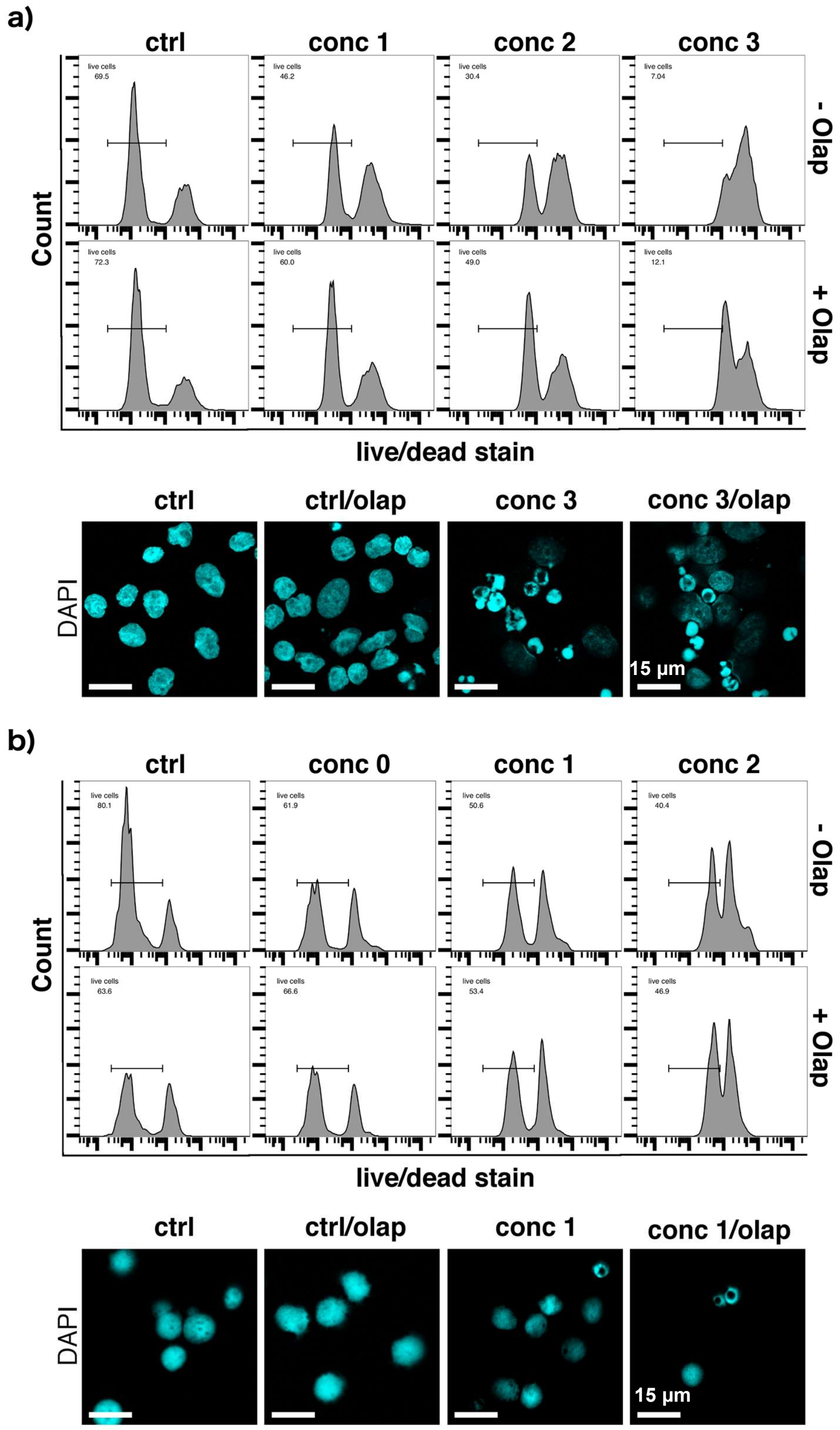


**Fig. D4** Parthanatos features in primary cells from AML donors according to toxicity rescue by Olaparib (Olap) and the presence of ring-shaped nuclei examined by DAPI staining. **a)** 7 / 17-008 and **b)** 8 / 17-016. Pretreatment: 1 µM Olaparib o/n; drug treatment: 24 h. Conc 0: 1 µM ara-C + 0.06 µM ida, conc 1: 5 µM ara-C + 0.3 µM ida, conc 2: 15 µM ara-C + 0.9 µM ida, conc 3: 30 µM ara-C + 1.8 µM ida.

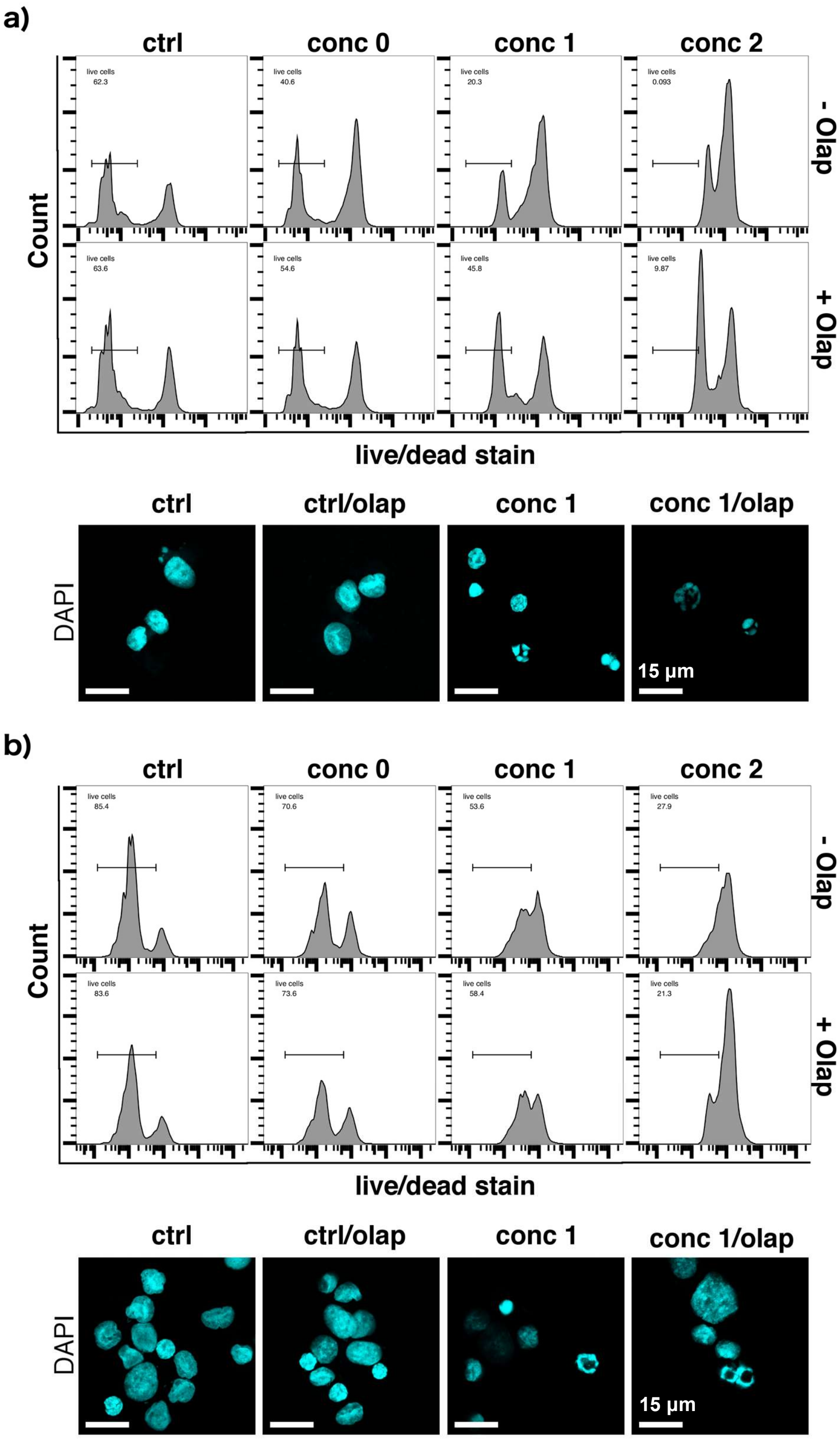


**Fig. D5** Parthanatos features in primary cells from AML donors according to toxicity rescue by Olaparib (Olap) and the presence of ring-shaped nuclei examined by DAPI staining. **a)** 9 / 17-036 and **b)** 10 / 17-063. Pretreatment: 1 µM Olaparib o/n; drug treatment: 24 h. Conc 0: 1 µM ara-C + 0.06 µM ida, conc 1: 5 µM ara-C + 0.3 µM ida, conc 2: 15 µM ara-C + 0.9 µM ida.

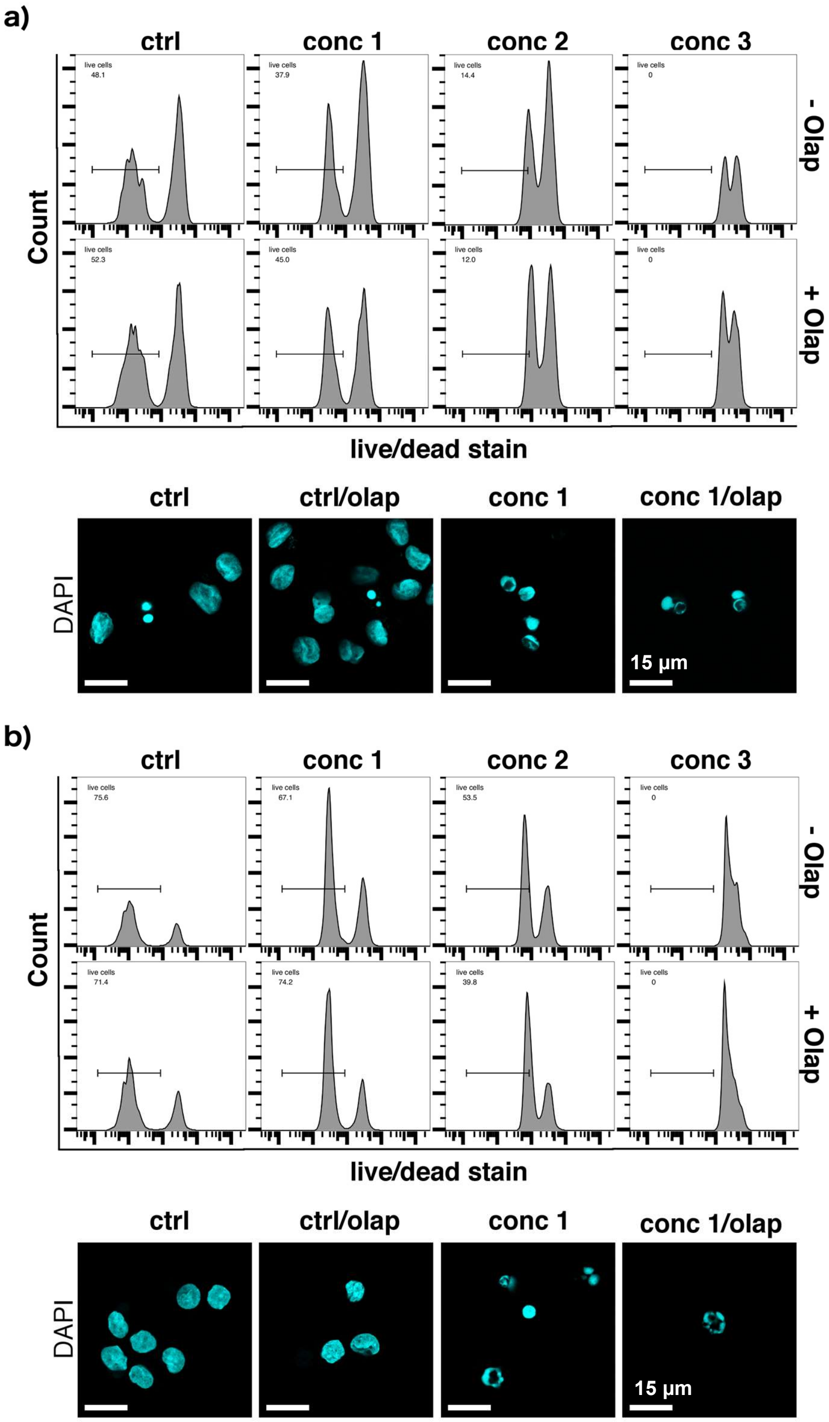


**Fig. D6** Parthanatos features in primary cells from AML donors according to toxicity rescue by Olaparib (Olap) and the presence of ring-shaped nuclei examined by DAPI staining. **a)** 11* / PID 145 and **b)** 12* / PID 160. Pretreatment: 1 µM Olaparib o/n; drug treatment: 24 h. Conc 1: 5 µM ara-C + 0.3 µM ida, conc 2: 15 µM ara-C + 0.9 µM ida, conc 3: 30 µM ara-C + 1.8 µM ida. *Bone marrow isolates.

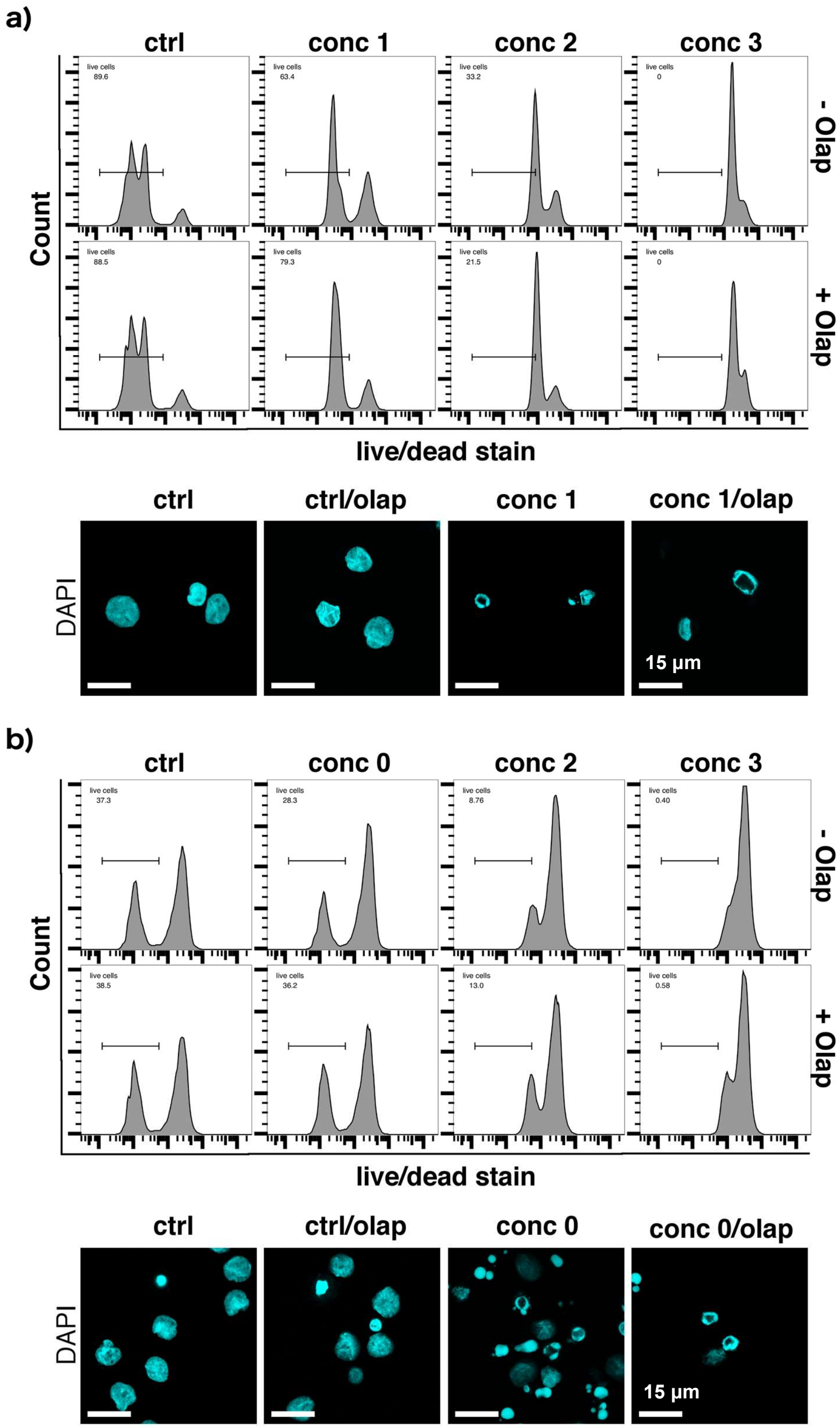


**Fig. D7** Parthanatos features in primary cells from AML donors according to toxicity rescue by Olaparib (Olap) and the presence of ring-shaped nuclei examined by DAPI staining. **a)** 13* / PID 198 and **b)** 14 / PID 752. Pretreatment: 1 µM Olaparib o/n; drug treatment: 24 h. Conc 0: 1 µM ara-C + 0.06 µM ida, conc 1: 5 µM ara-C + 0.3 µM ida, conc 2: 15 µM ara-C + 0.9 µM ida, conc 3: 30 µM ara-C + 1.8 µM ida. *Bone marrow isolates.

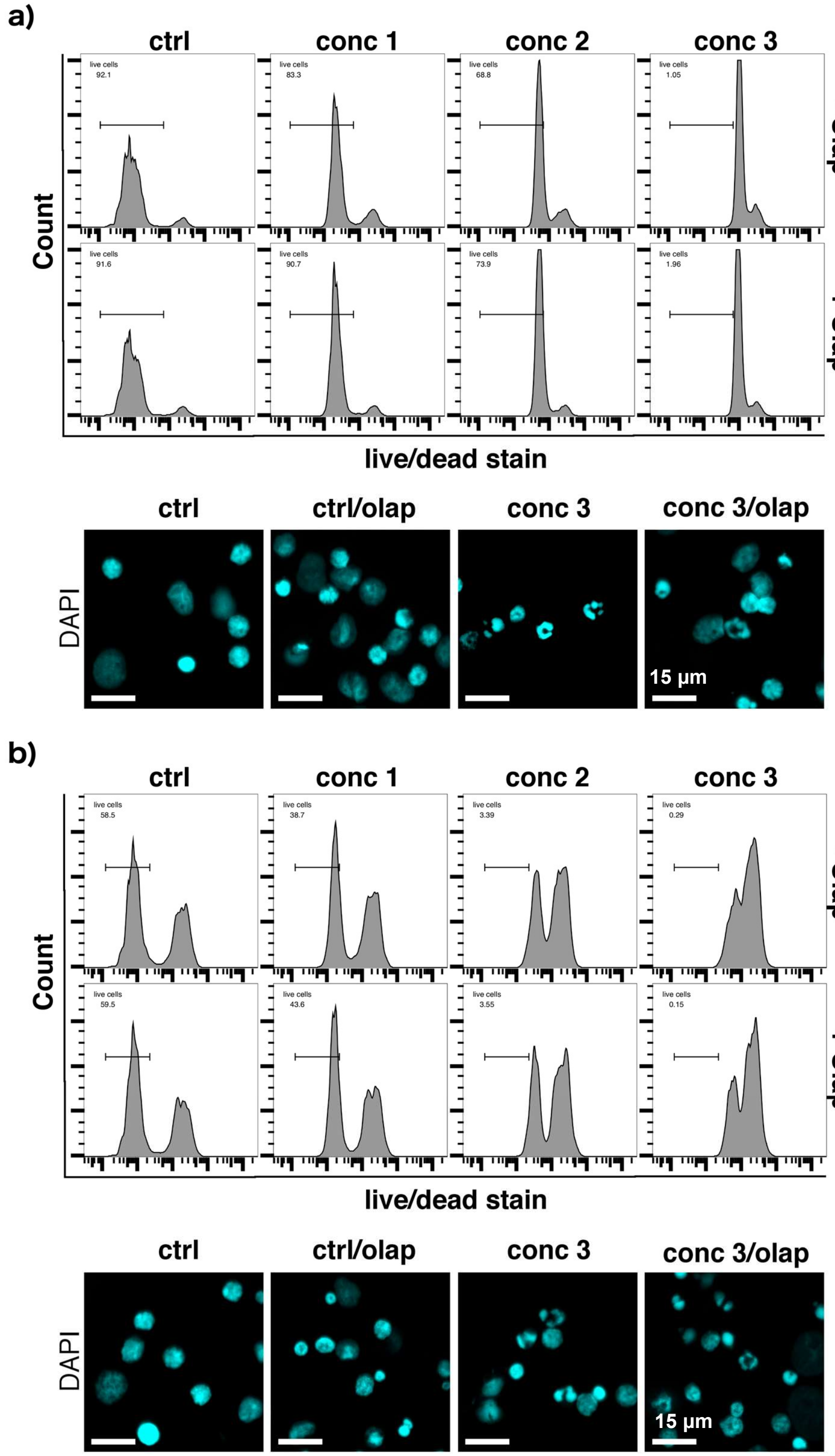


**Fig. D8** Parthanatos features in primary cells from AML donors according to toxicity rescue by Olaparib (Olap) and the presence of ring-shaped nuclei examined by DAPI staining. **a)** 15 / PID 52 and **b)** 16 / PID 528. Pretreatment: 1 µM Olaparib o/n; drug treatment: 24 h. Conc 1: 5 µM ara-C + 0.3 µM ida, conc 2: 15 µM ara-C + 0.9 µM ida, conc 3: 30 µM ara-C +1.8 µM ida.

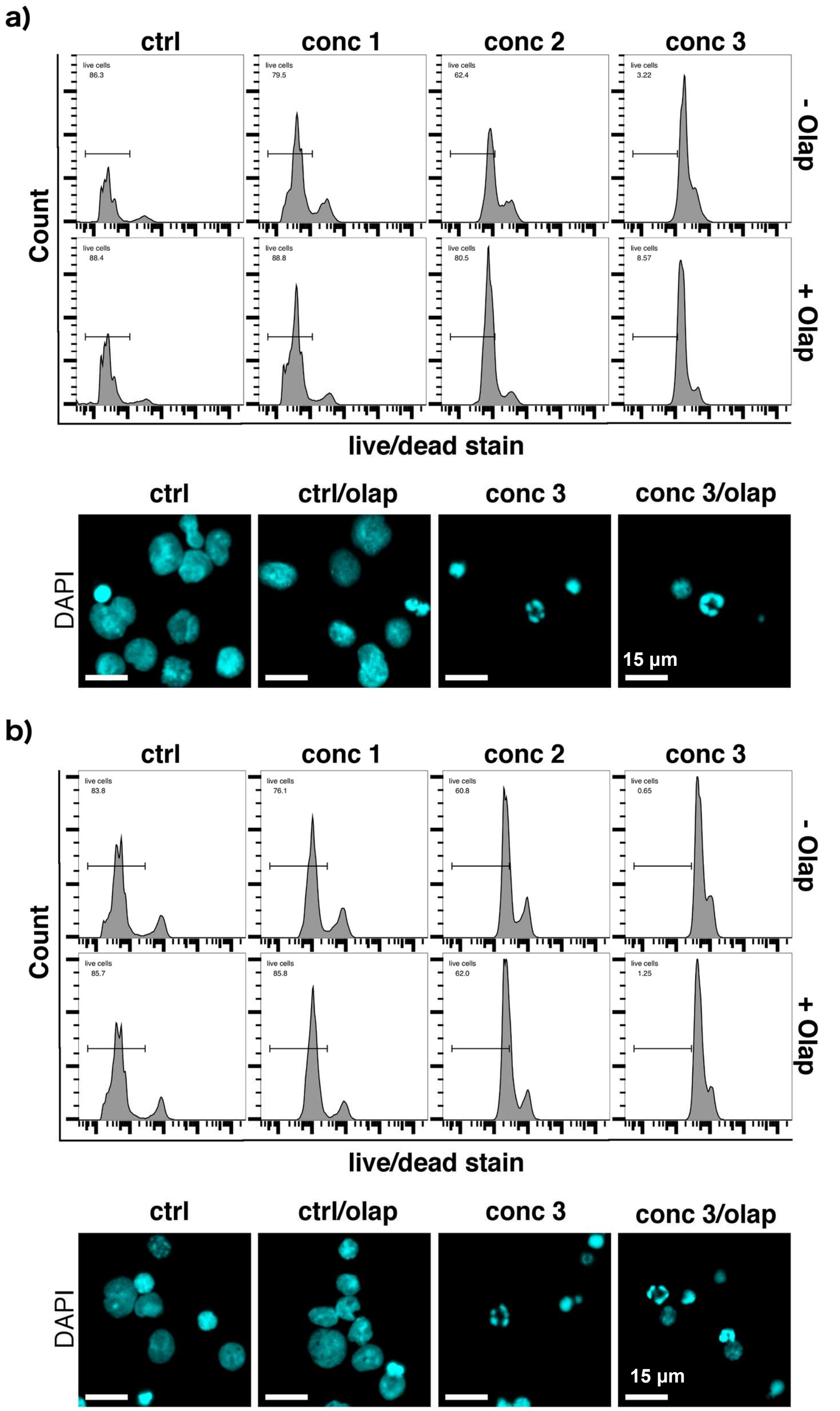


**Fig. D9** Parthanatos features in primary cells from AML donors according to toxicity rescue by Olaparib (Olap) and the presence of ring-shaped nuclei examined by DAPI staining. **a)** 17 / PID 242 and **b)** 18 / PID 127. Pretreatment: 1 µM Olaparib o/n; drug treatment: 24 h. Conc 1: 5 µM ara-C + 0.3 µM ida, conc 2: 15 µM ara-C + 0.9 µM ida, conc 3: 30 µM ara-C + 1.8 µM ida.

### Supplementary Figure E: Primary samples from AML patients exhibiting zero or one parthanatos features.

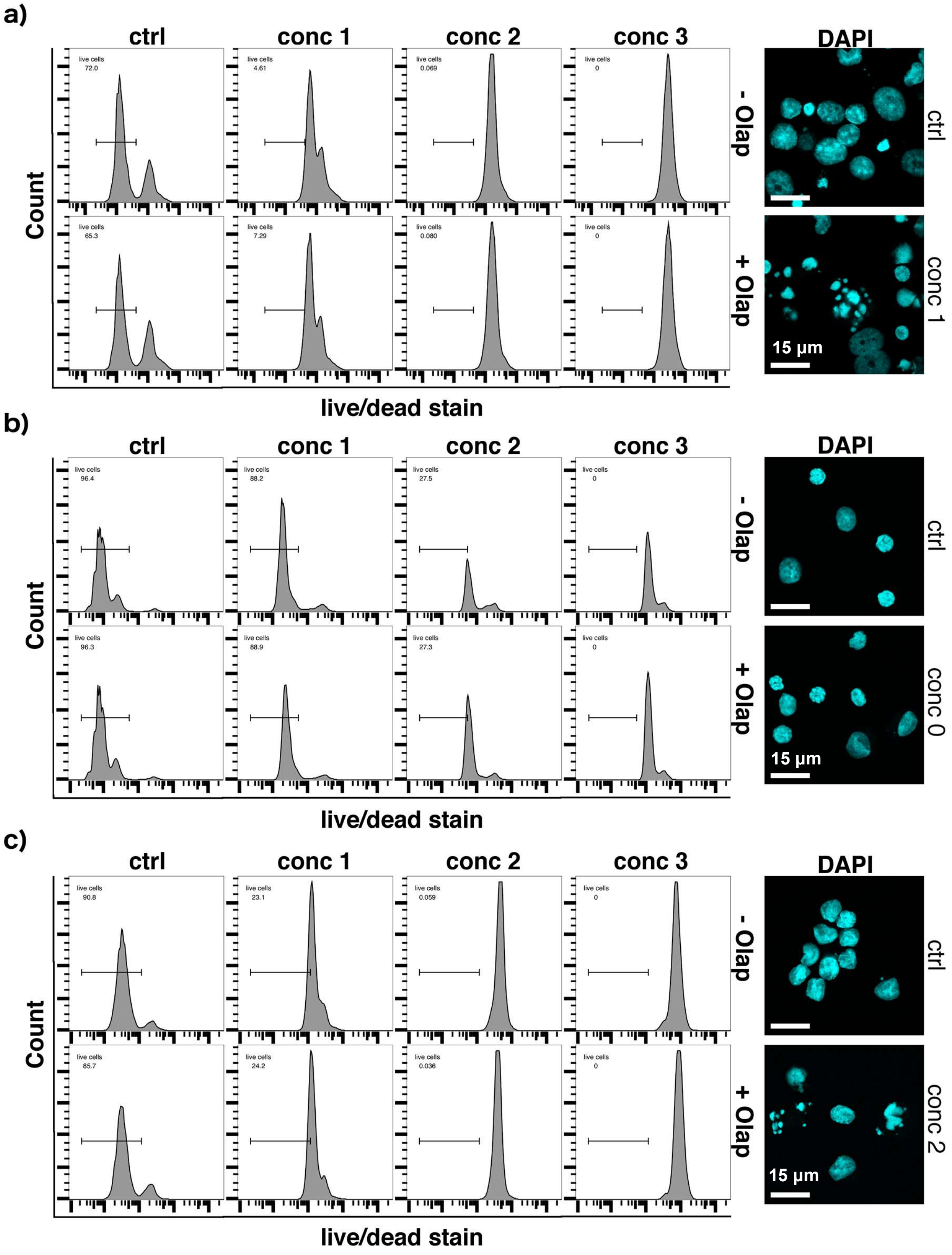


**Fig. E1** No apparent parthanatos features in primary cells from AML donors according to toxicity rescue by Olaparib (Olap) and the presence of ring-shaped nuclei examined by DAPI staining. **a)** 19 / 04-015, **b)** 20 / 15-119 and **c)** 21 / 15-130. Pretreatment: 1 µM Olaparib o/n; drug treatment: 24 h. Conc 0: 1 µM ara-C + 0.06 µM ida, conc 1: 5 µM ara-C + 0.3 µM ida, conc 2: 15 µM ara-C + 0.9 µM ida, conc 3: 30 µM ara-C + 1.8 µM ida.

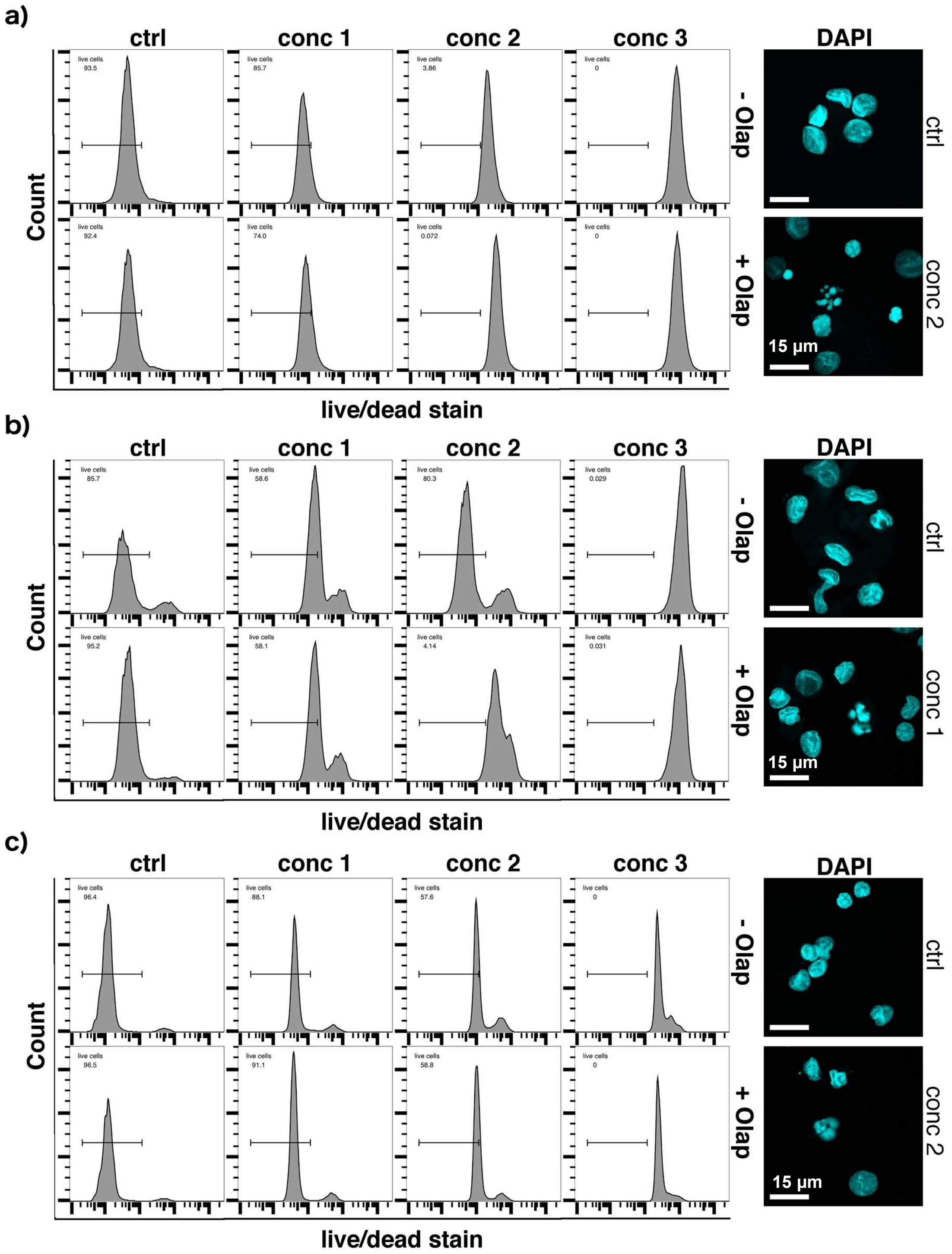


**Fig. E2** No apparent parthanatos features in primary cells from AML donors according to toxicity rescue by Olaparib (Olap) and the presence of ring-shaped nuclei examined by DAPI staining. **a)** 22 / 16-007, **b)** 23 / 16-062 and **c)** 24 / 16-068. Pretreatment: 1 µM Olaparib o/n; drug treatment: 24 h. Conc 1: 5 µM ara-C + 0.3 µM ida, conc 2: 15 µM ara-C + 0.9 µM ida, conc 3: 30 µM ara-C + 1.8 µM ida.

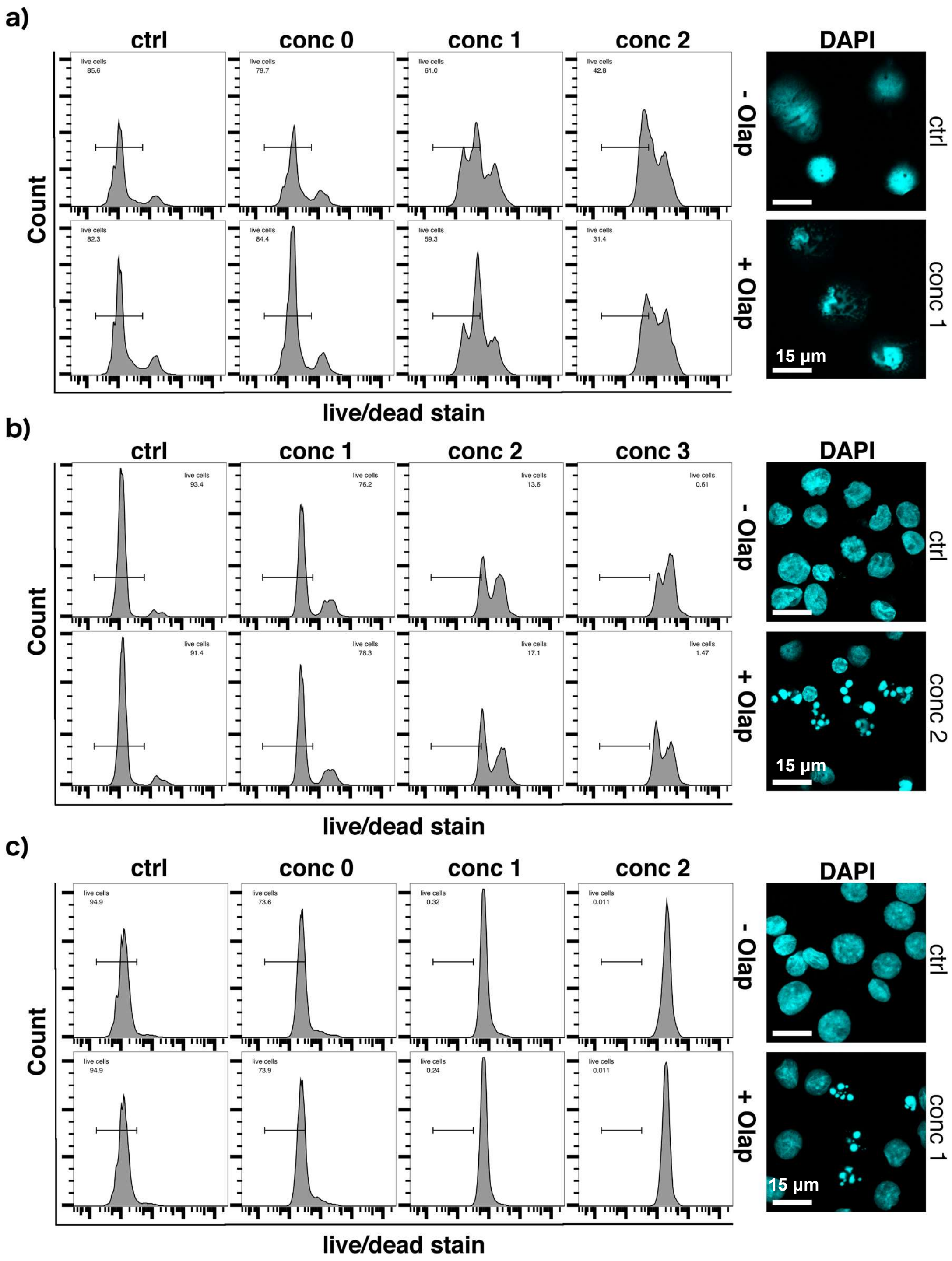


**Fig. E3** No apparent parthanatos features in primary cells from AML donors according to toxicity rescue by Olaparib (Olap) and the presence of ring-shaped nuclei examined by DAPI staining. **a)** 25 / 17-003, **b)** 26 / 17-024 and **c)** 27* / 17-039. Pretreatment: 1 µM Olaparib o/n; drug treatment: 24 h. Conc 0: 1 µM ara-C + 0.06 µM ida, conc 1: 5 µM ara-C + 0.3 µM ida, conc 2: 15 µM ara-C + 0.9 µM ida, conc 3: 30 µM ara-C + 1.8 µM ida. *Bone marrow isolate.

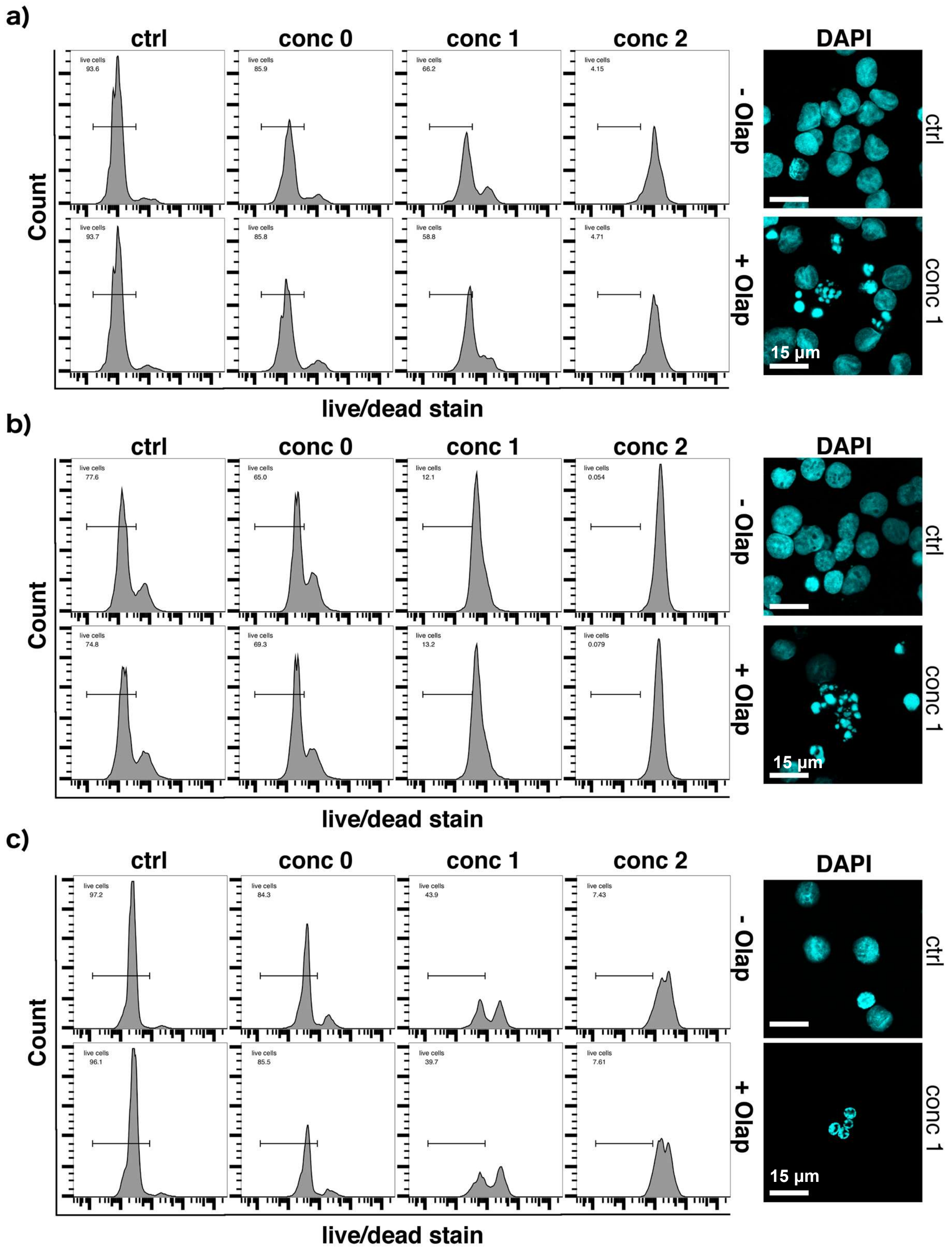


**Fig. E4** Zero (**a**,**b**) or one (**c**) apparent parthanatos feature in primary cells from AML donors according to toxicity rescue by Olaparib (Olap) and the presence of ring-shaped nuclei examined by DAPI staining. **a)** 28 / 17-045, **b)** 29* / 17-014 and **c)** 30* / PID 624. Pretreatment: 1 µM Olaparib o/n; drug treatment: 24 h. Conc 0: 1 µM ara-C + 0.06 µM ida, conc 1: 5 µM ara-C + 0.3 µM ida, conc 2: 15 µM ara-C + 0.9 µM ida. *Bone marrow isolates.

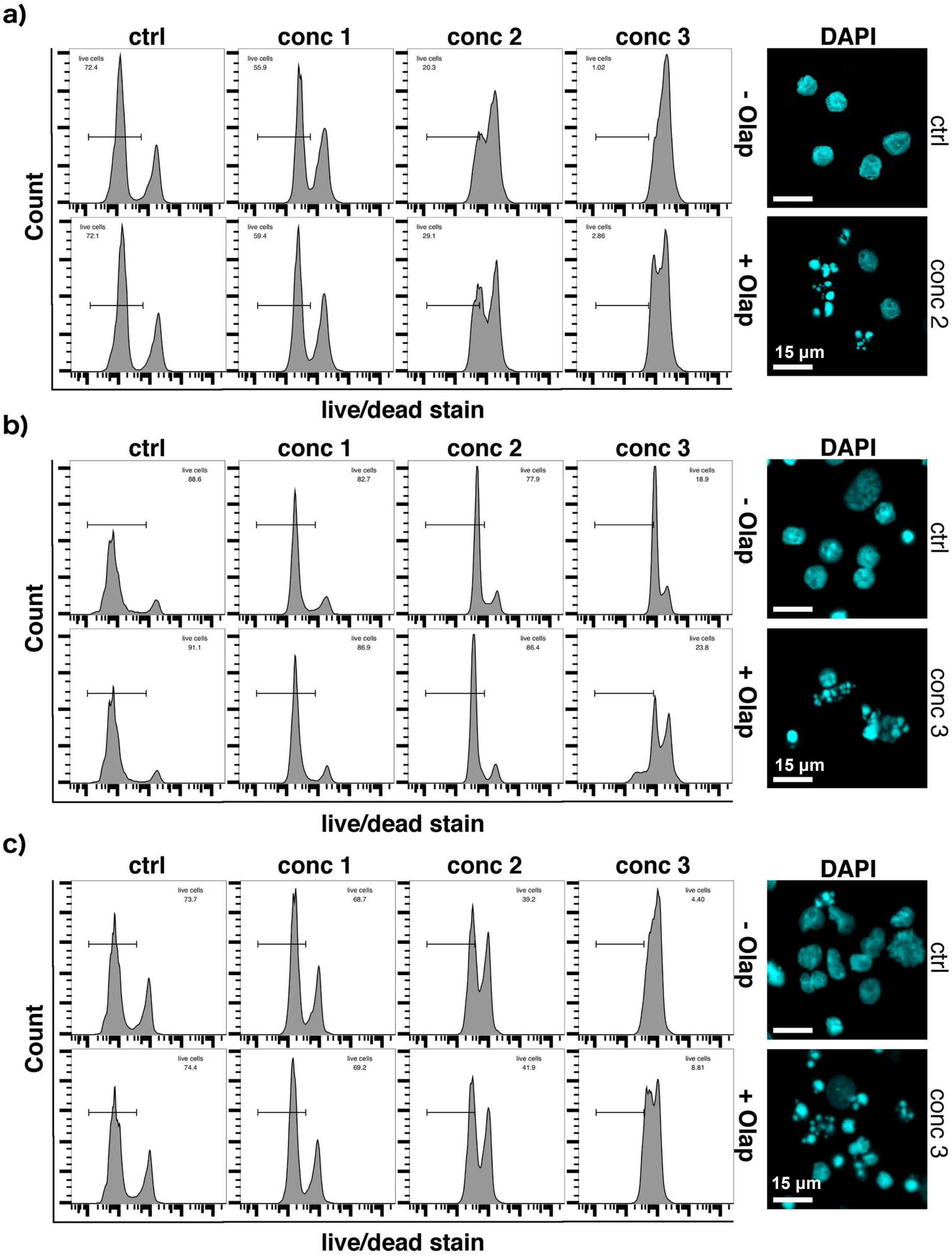


**Fig. E5** One (**a,b**) or zero (**c**) apparent parthanatos features in primary cells from AML donors according to toxicity rescue by Olaparib (Olap) and the presence of ring-shaped nuclei examined by DAPI staining. **a)** 31 / PID 625, **b)** 32 / PID 766 and **c)** 33 / PID 154. Pretreatment: 1 µM Olaparib o/n; drug treatment: 24 h. Conc 1: 5 µM ara-C + 0.3 µM ida, conc 2: 15 µM ara-C + 0.9 µM ida, conc 3: 30 µM ara-C + 1.8 µM ida.

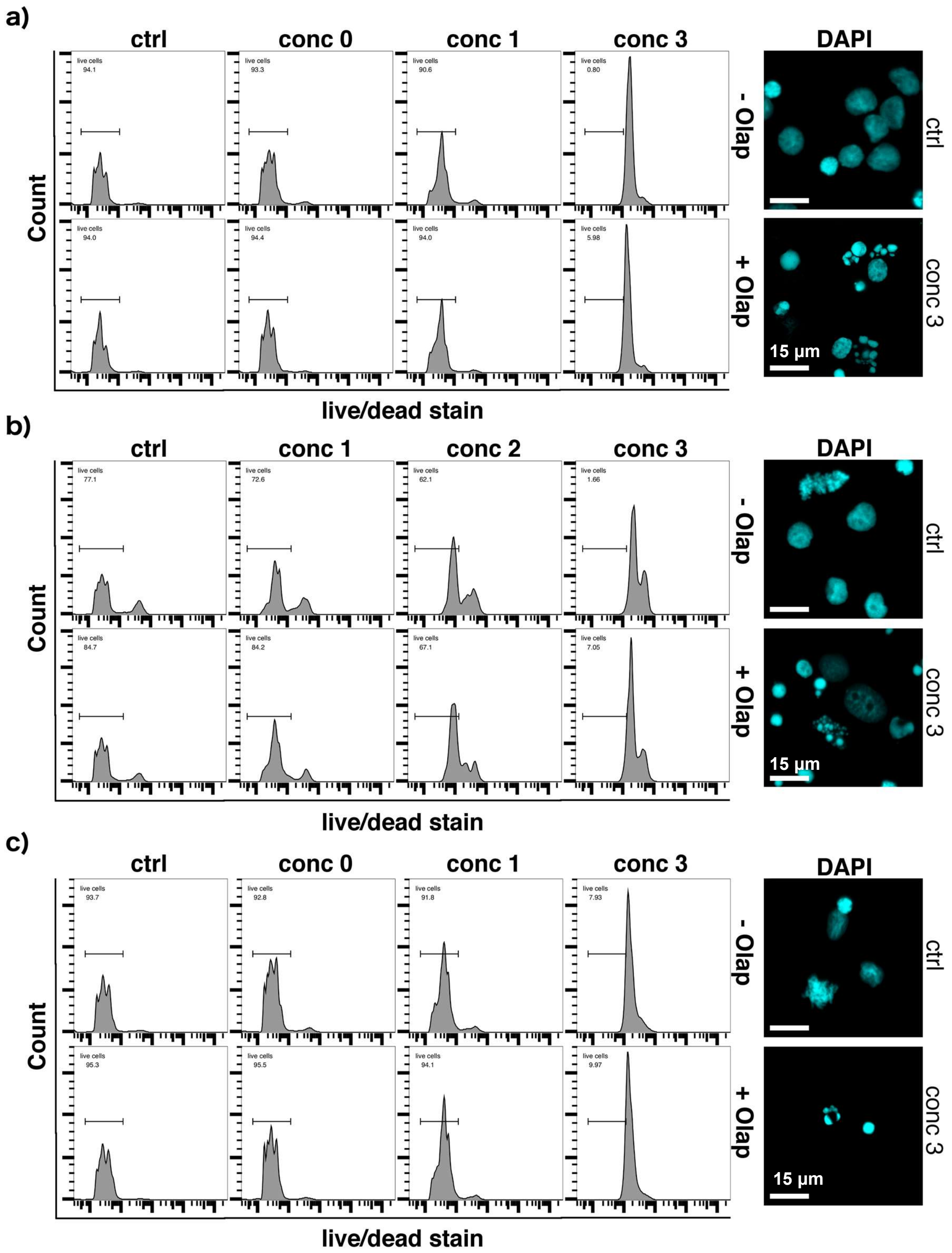


**Fig. E6** Zero (**a**) or one (**b**,**c**) apparent parthanatos feature in primary cells from AML donors according to toxicity rescue by Olaparib (Olap) and the presence of ring-shaped nuclei examined by DAPI staining. **a)** 34 / PID 218, **b)** 35 / PID 469 and **c)** 36 / PID 103. Pretreatment: 1 µM Olaparib o/n; drug treatment: 24 h. Conc 0: 1 µM ara-C + 0.06 µM ida, conc 1: 5 µM ara-C + 0.3 µM ida, conc 3: 30 µM ara-C + 1.8 µM ida.

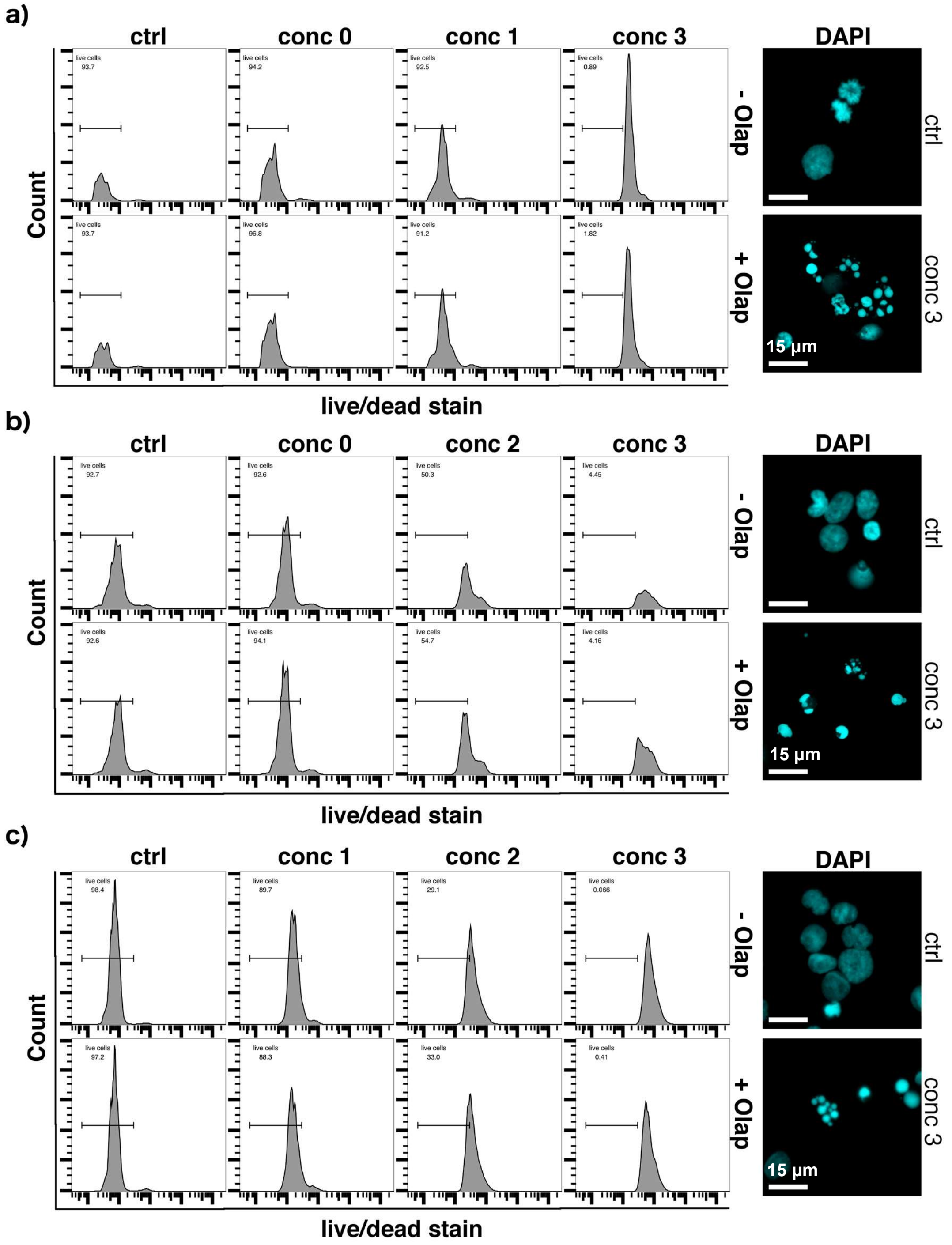


**Fig. E7** No apparent parthanatos features in primary cells from AML donors according to toxicity rescue by Olaparib (Olap) and the presence of ring-shaped nuclei examined by DAPI staining. **a)** 37/ PID 176, **b)** 38 / PID 171 and **c)** 39 / PID 517. Pretreatment: 1 µM Olaparib o/n; drug treatment: 24 h. Conc 0: 1 µM ara-C + 0.06 µM ida, conc 1: 5 µM ara-C + 0.3 µM ida, conc 2: 15 µM ara-C + 0.9 µM ida, conc 3: 30 µM ara-C + 1.8 µM ida.

### Supplementary Figure F: Overall % survival analysis

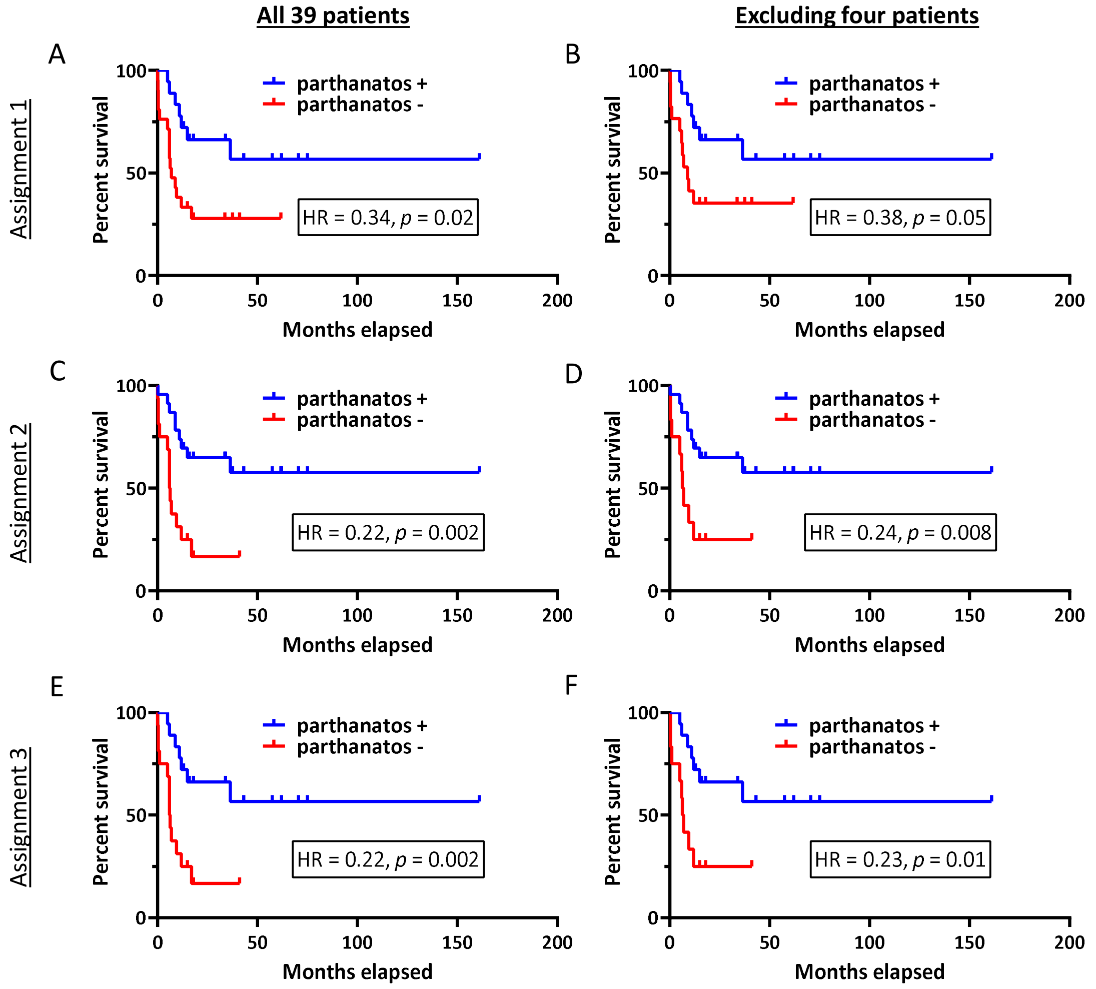


**Fig. F. Overall % survival (OS) analysis of the parthanatos positive (+) versus negative (-) groups. A)** Kaplan-Meier OS curves comparing +/- groups using Assignment 1 of both parthanatos features (n = 18) versus one or zero features (n = 21). **B)** Kaplan-Meier OS curves comparing +/- groups using Assignment 1 and excluding four patients who did not receive curative chemotherapy with ara-C. **C)** Kaplan-Meier OS curves comparing +/- groups using Assignment 2 of one or both parthanatos features (n = 23) versus zero features (n = 16). **D)** Kaplan-Meier OS curves comparing +/- groups using Assignment 2 and excluding four patients who did not receive curative chemotherapy with ara-C. **E)** Kaplan-Meier OS curves comparing +/- groups using Assignment 3 of both parthanatos features (n = 18) versus zero features (n = 16). **F)** Kaplan-Meier OS curves comparing +/- groups using Assignment 3 and excluding four patients who did not receive curative chemotherapy with ara-C.

### Supplementary Figure G: Event-free % survival analysis

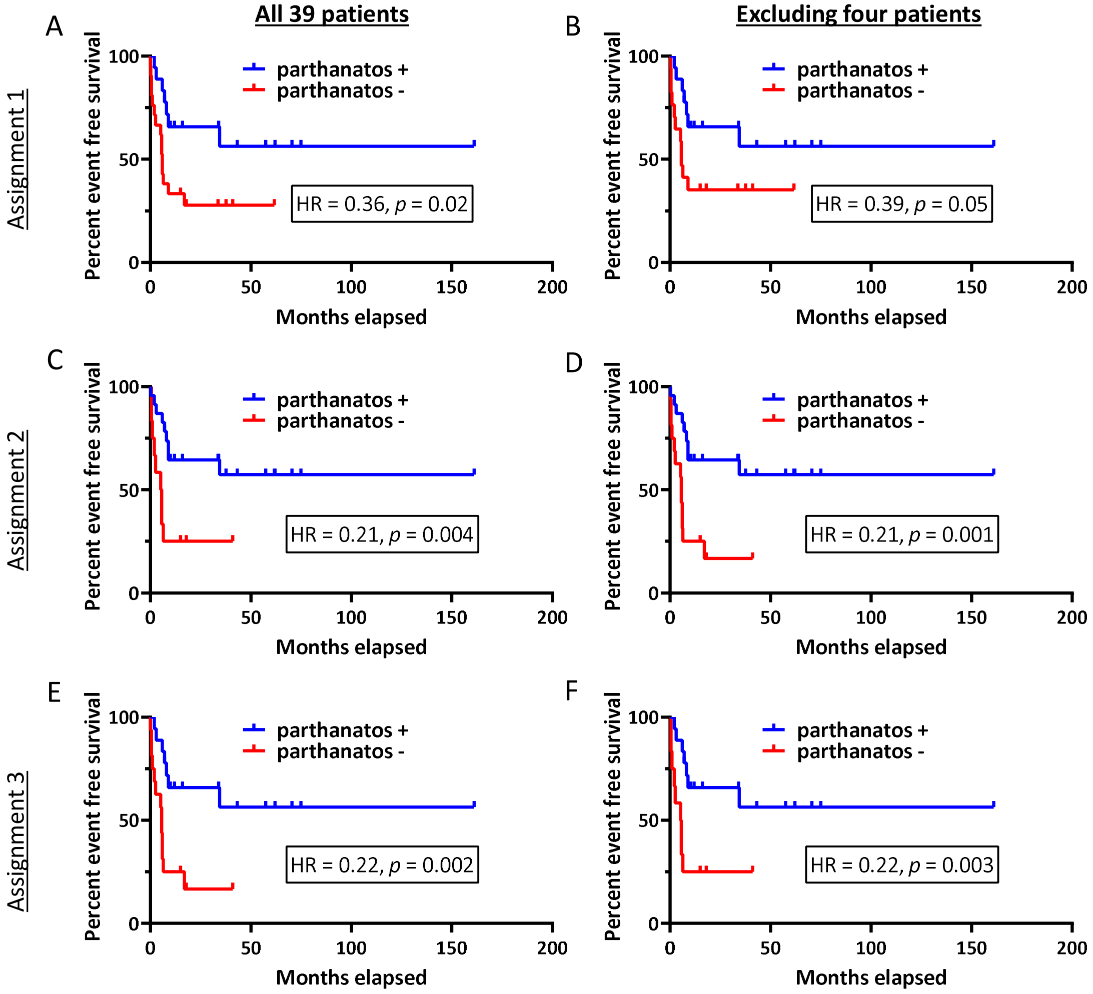


**Fig. G. Event free % survival (EFS) analysis of parthantos positive (+) versus negative (-) groups. A)** Kaplan-Meier estimate EFS curves comparing +/- groups using Assignment 1 of both parthanatos features (N = 18) versus one or zero features (N = 21). **B)** EFS curves comparing +/- groups using Assignment 1 and excluding four patients who did not receive curative chemotherapy with ara-C. **C)** EFS curves comparing +/- groups using Assignment 2 of one or both parthanatos features (N = 23) versus zero features (N = 16). **D)** EFS curves comparing +/- groups using Assignment 2 and excluding four patients who did not receive curative chemotherapy with ara-C. **E)** EFS curves comparing +/- groups using Assignment 3 of both parthanatos features (N = 18) versus zero features (N = 16). **F)** EFS curves comparing +/- groups using Assignment 3 and excluding four patients who did not receive curative chemotherapy with ara-C.

### Supplementary Figure H: mRNA expression of parthanatos-associated genes

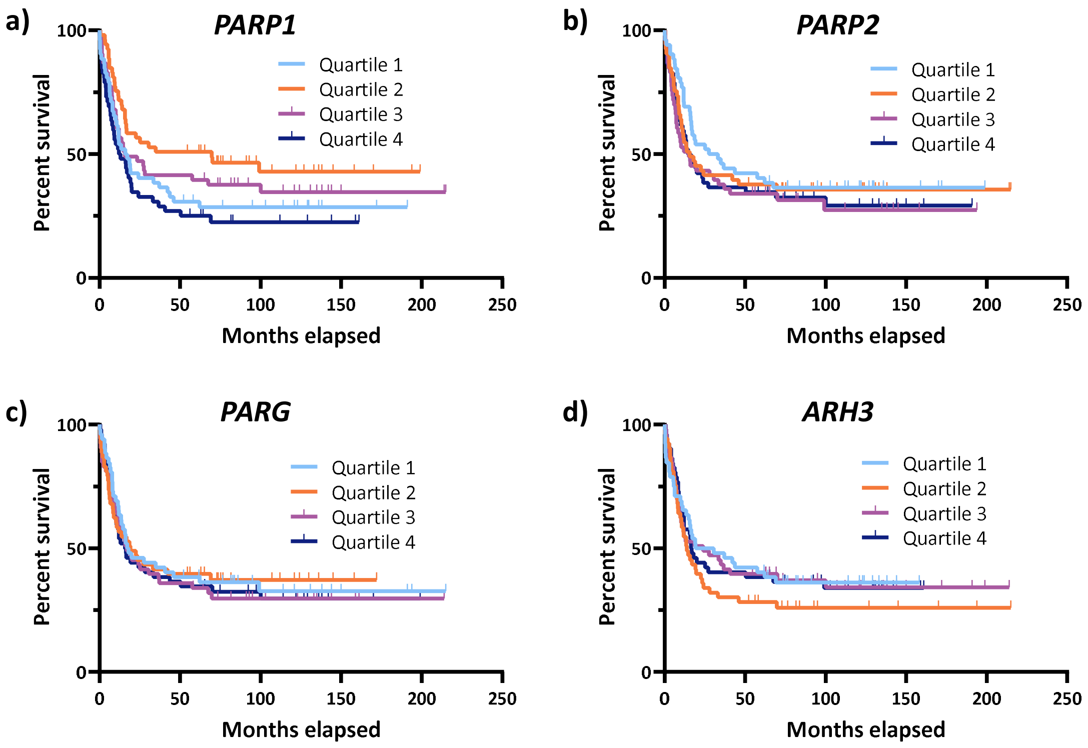


**Fig. H. Analysis of mRNA expression in samples from 210 AML patients (FAB subtypes M4 and M5) receiving curative treatment with ara-C and idarubicin.** Patients were grouped into four equal quartiles according to the relative mRNA expression from each gene (quartile 1, <25%; quartile 2, 25-50%; quartile 3, 50-75%; and quartile 4, >75%) and overall percentage survivals were plotted using Kaplan-Meier survival estimates. **a)** *PARP1* **b)** *PARP2*; **c)** *PARG*; and **d)** *ARH3*.

### Supplementary Figure I: PARP-1 expression analysis

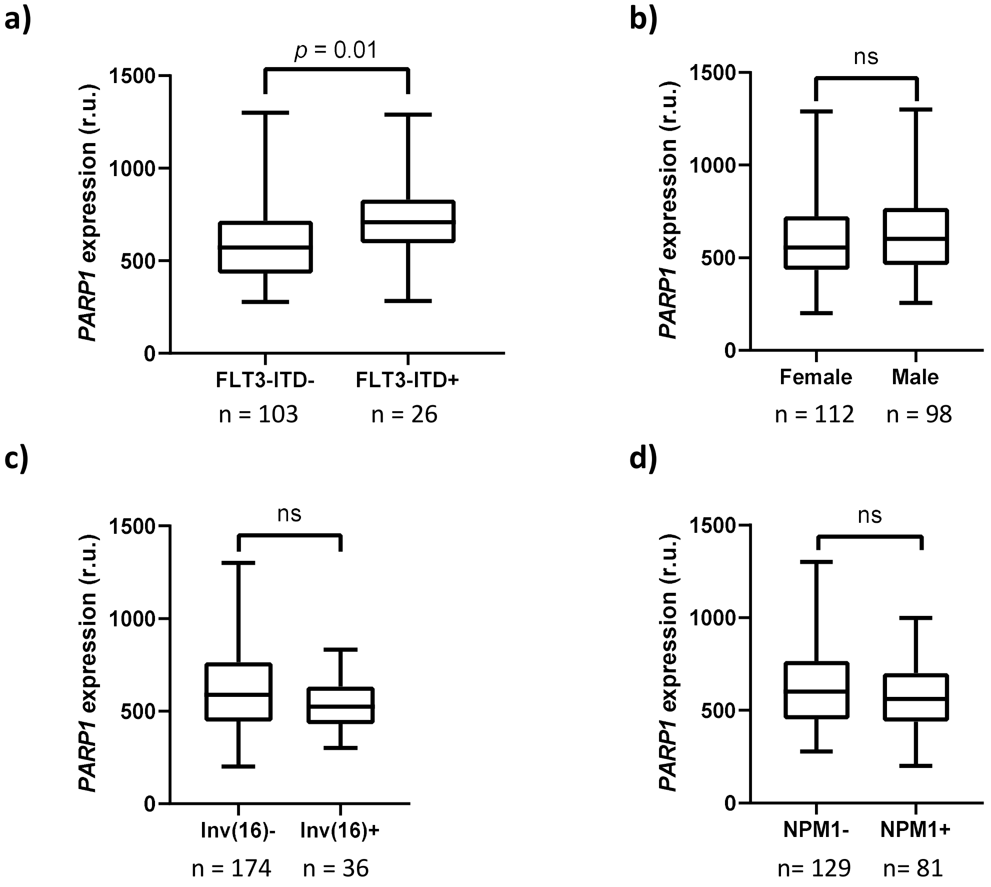


**Figure I. Analysis of mRNA expression (AML FAB subtypes M4 and M5) receiving curative treatment with ara-C and idarubicin.** *PARP1* expression vs AML mutations and sex. **a)** *PARP1* mRNA was significantly higher in patients with the FLT3-ITD mutation – even when NPM1+ mutants are removed from the analysis as shown here. *PARP1* expression was independent of **b)** sex (n = 210 total); **c)** inv(16) status (n= 210 total); and **d)** NPM1 mutational status (n = 210 total).
